# Egocentric Asymmetric Coding in Sensory Cortical Border Cells

**DOI:** 10.1101/2021.03.11.434952

**Authors:** Xiaoyang Long, Bin Deng, Jing Cai, Zhe Sage Chen, Sheng-Jia Zhang

## Abstract

Both egocentric and allocentric representations of space are essential to spatial navigation. Although some studies of egocentric coding have been conducted within and around the hippocampal formation, externally anchored egocentric spatial representations have not yet been fully explored. Here we record and identify two subtypes of border cell in the rat primary somatosensory cortex (S1) and secondary visual cortex (V2). Subpopulations of S1 and V2 border cells exhibit rotation-selective asymmetric firing fields in an either clockwise (CW) or counterclockwise (CCW) manner. CW- and CCW-border cells increase their firing rates when animals move unidirectionally along environmental border(s). We demonstrate that both CW- and CCW-border cells fire in an egocentric reference frame relative to environmental borders, maintain preferred directional tunings in rotated, stretched, dark as well as novel arenas, and switch their directional firings in the presence of multi-layer concentric enclosures. These findings may provide rotation-selective egocentric reference frames within a larger spatial navigation system, and point to a common computational principle of spatial coding shared by multiple sensory cortical areas.

**Highlights:**

- Egocentric border cells are present in rat S1 and V2
- Subtypes of border cells display egocentric asymmetric coding
- Egocentric and allocentric streams coexist in sensory cortices
- Rotation-selective asymmetric firing is robust with environmental manipulations

## INTRODUCTION

Goal-directed spatial navigation is a complex multi-function task that requires memory and sensory processing, sensorimotor integration, planning, and decision making. In the past few decades, spatially-tuned neurons have been discovered across multiple brain areas, starting with the traditional hippocampal-entorhinal network and the neighboring structures (Gofman et al., 2019; Hafting et al., 2005; O’Keefe and Dostrovsky, 1971; Peyrache et al., 2017; Wang et al., 2018) and extending to other high-order association and sensory cortical areas (Alexander et al., 2020; Long et al., 2021; Long et al., 2020; Long and Zhang, 2021). Allocentric (world-centered, viewpoint-invariant) and egocentric (self-centered) representations of space define two distinct reference frames and coordinate systems for coding environmental features (such as the distance, direction, position, boundaries, and objects) (Bicanski and Burgess, 2020; Wang et al., 2020). The classical place cells, grid cells, and head-direction cells are thought to primarily represent allocentric information in the hippocampal-entorhinal system, whereas egocentric information relative to the body, eye or head, has been observed in sensorimotor representations in the posterior parietal cortex (PPC), postrhinal cortex, retrosplenial cortex (RSC), and secondary motor cortex (M2) (Alexander et al., 2020; Hinman et al., 2019; LaChance et al., 2019; Olson et al., 2020). Multiple lines of evidence have suggested that egocentric sensory information derived from sensory and motor cortical areas may be transformed into allocentric representations of space that interact with the allocentric hippocampal-entorhinal system (Olson et al., 2020; Whitlock et al., 2012; Wilber et al., 2014). Additionally, the primarily allocentric hippocampal-entorhinal network also displays egocentric spatial representations, including egocentric goal-direction cells (Sarel et al., 2017) and egocentric head-direction cells (Jercog et al., 2019). Recent rat experiments have demonstrated functionally distinct spatial cell types (i.e., place cells, border cells, head-direction cells and grid cells) in the primary somatosensory cortex (S1) and secondary visual cortex (V2) from freely foraging rats, suggesting the existence of a compact spatial map in the sensory cortices (Long et al., 2021; Long et al., 2020; Long and Zhang, 2021). Importantly, these four distinct types of spatially tuned sensory cortical cell showed robust responses to the environmental geometry, size, orientation, height and lighting conditions. In addition to the allocentric stream, we hypothesize that egocentric representations and coordinate transformation of space also exist in the sensory cortices, independent of or in parallel with the classical hippocampal formation.

Here we sought to identify and characterize egocentric spatial signals from layers IV-VI of the rat S1 and V2. We not only found allocentric representations of space in place cells, head-direction cells, grid cells, and boundary-vector/border cells, but also discovered egocentric representations of space in border cells, which respond to the environmental border or walls. We identified two novel subtypes of border cell in the rat S1 and V2, both of which increased their firing rates only at the edges of the navigation environment when animals moved unidirectionally in an either clockwise (CW) or counterclockwise (CCW) direction---which were referred to as CW-border cells and CCW-border cells, respectively. We demonstrated that, unlike classical allocentric border cells in the medial entorhinal cortex (MEC) and subiculum (Lever et al., 2009; Savelli et al., 2008; Solstad et al., 2008; Stewart et al., 2014), these CW- and CCW-border cells recorded from sensory cortices) are purely egocentric and process a unique egocentric-by-allocentric conjunctive coding of reference frames. Furthermore, it was surprising to find that the border cells in S1 and V2 shared similar robust firing properties across various environmental manipulations and controls of vestibular (whisker removal) and visual (light off) signals. The identification of egocentric asymmetric rotation coding of CW- and CCW-border cells in sensory cortices (S1 and V2) not only supports the hypothesis that sensory systems play a vital role in representing egocentric reference frames, but also suggests a shared map-based reference system and computation mechanism across distributed brain areas to perform coordinated spatial navigation.

## RESULTS

### Extracellular Recordings in Rat S1 and V2

We collected *in vivo* extracellular recordings from the left or right hemisphere of implanted male Long-Evans adult rats (*n* = 20) using self-assembled movable micro-drives connected to four tetrodes. As documented previously (Long et al., 2021; Long et al., 2020; Long and Zhang, 2021), recording electrodes were targeted into either the primary somatosensory cortex (S1) or the secondary visual cortex (V2). Postmortem histological reconstruction confirmed that all of the recording electrodes were located in either S1 or V2 (**Figure S1A**).

We recorded the spiking activity from either S1 or V2 as rats foraged freely in a standard 1 × 1 m^2^ open square arena with 50-cm-high walls. From 735 recording sessions, we identified a total of 2485 and 1870 well-isolated single units from S1 and V2, respectively. Similar to those spatially tuned cells previously identified within the hippocampal formation (Hafting et al., 2005; Lever et al., 2009; O’Keefe and Dostrovsky, 1971; Sargolini et al., 2006; Solstad et al., 2008; Taube et al., 1990), the S1 and V2 neurons from cortical layers IV-VI consisted of place cells, head-direction cells, border cells, grid cells, and conjunctive cells (Long et al., 2021; Long and Zhang, 2021).

### Regular, CW- and CCW-Border Cells Coexisted in S1 and V2

Border cells increase their firing rates exclusively along or near one or multiple borders of an open arena (Solstad et al., 2008). We classified a recorded S1 or V2 unit as a border cell if its border score was greater than the 99^th^ percentile of population shuffled scores (**Figure S1B**). In total, 148 out of 2485 (5.96%) S1 as well as 161 out of 1870 (8.61%) V2 units passed the border score classification threshold and were classified as border cells. This percentage was significantly higher than expected by random selection from the entire shuffled population (**Figure S1B**; *Z* = 40.43, *p* < 0.001; binomial test with expected *p*_0_ of 0.01 among large samples). We found no significant difference in border tuning property or firing property between S1 and V2 border cells (**Figure S1C**; two-sided unpaired *t*-test, *n* = 148 and 161, border score: *p* = 0.2; mean firing rate: *p* = 0.3) and therefore combined data for further analysis.

Similar to MEC border cells in MEC, a large proportion of the S1 and V2 border cells increased their firing rates near one or multiple borders of the open arena in a bidirectional manner (**Figure 1A**). We defined these cells, which exhibited symmetric firing patterns with respect to animals’ head direction (**Figure 1A**), as regular border cells. Additionally, a subset of recorded S1 and V2 border cells presented asymmetric rotation tuning, and displayed unidirectional firing in an either CW or CCW manner (**Figure S2**). To classify this subset of S1 and V2 border cells, we calculated the ratio of each neuron’s firing rate during clockwise versus counterclockwise movement. If the ratio was between 0.5 and 2, then the border cell was classified as a regular border cell (**Figure 1A**); in contrast, if the ratio was greater than 2, the border cell was classified as a CW-border cell (**Figure 1B**); if the ratio was less than 0.5, the border cell was classified as a CCW-border cell (**Figure 1C**). Accordingly, 27.18% (n = 84/309) of identified S1+V2 border cells were classified as CW-border cells, and 23.62% (n = 73/309) of identified S1+V2 border cells were classified as CCW-border cells, and 49.49% (n = 152/309) were classified as regular border cells (**Figure 1D**). The distribution of regular, CW- and CCW-border cells had the tendency to cluster in layer-V (**Figure 1G**) but showed no significant difference across cortical layers.

**Figure 1.**
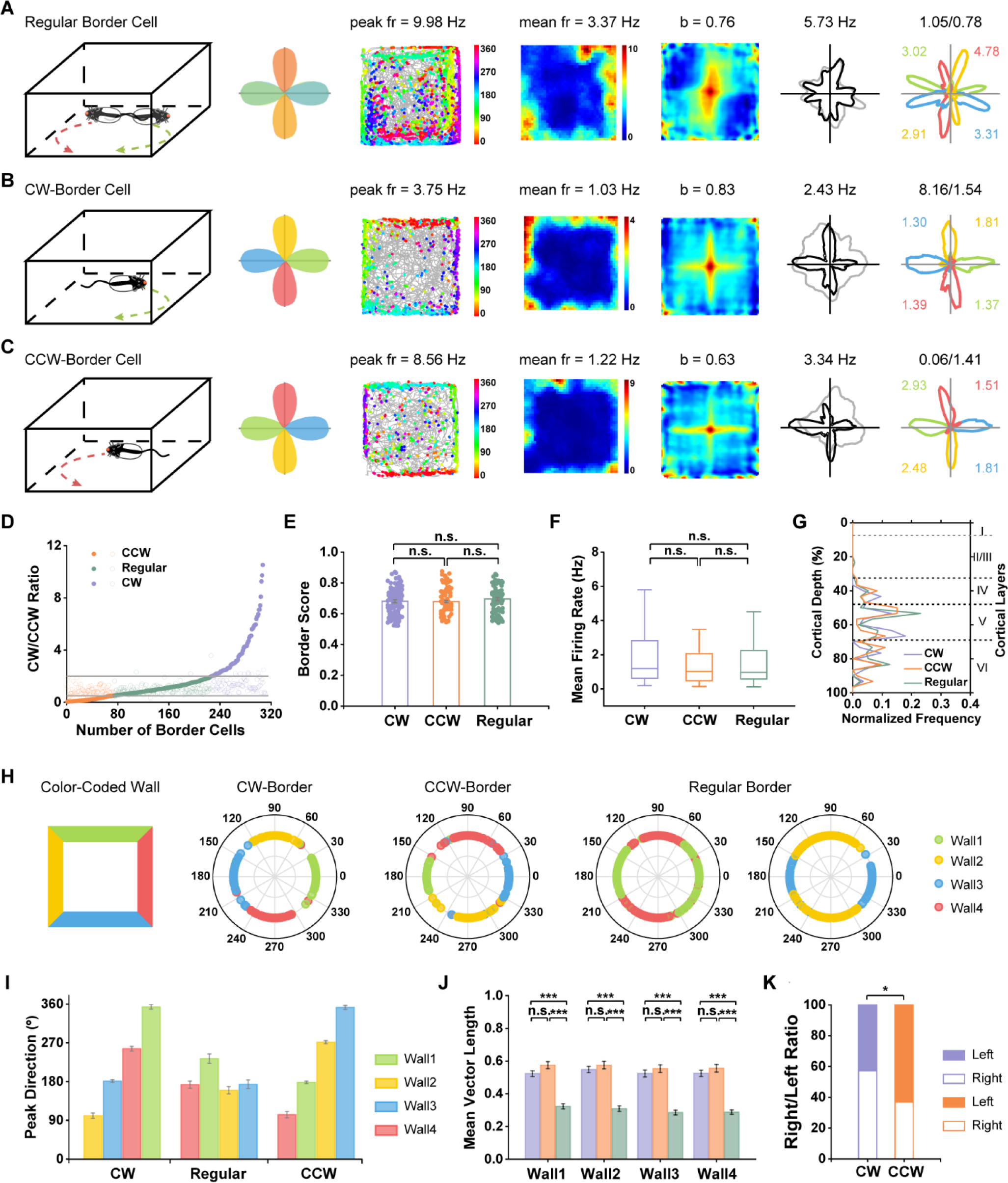
Border Cells from the Rat S1 and V2 Showed Symmetric and Asymmetric Firing Patterns in Either Bidirectional or Unidirectional Manner. (**A-C**) Representative subtype examples of border cells from the rat S1 and V2. Schematic of symmetric head direction tuning for regular (A), CW- (B) and CCW- (C) border cells (1st column); schematic diagrams of heading direction tuning along four color-coded walls of the square enclosure (2nd column); color-coded cross trajectory (grey line) with superimposed directional spike locations (color circles indicate the corresponding head direction; color bar shows the directional range 0°-360°) (3rd column); heat maps of spatial firing rate (4th column); autocorrelation diagrams (5th column); head-direction tuning curves (black) plotted against dwell-time polar plot (grey) (6th column); head-direction tuning curves for the corresponding color-coded four walls with peak firing rates labelled (7th column). Firing rate was color-coded with dark blue (red) indicating minimal (maximal) firing rate. The scale of the autocorrelation maps was twice that of the spatial firing rate maps. Peak firing rate (fr), mean firing rate, border score (b), peak angular rate, the ratios of total time and firing rate between animal’s heading in a clockwise direction and counterclockwise directions (time_cw/ccw and fr_cw/ccw), for each border cell are labelled at the top of the panels. **(D)** The distribution of time_cw/ccw (open dot) and fr_cw/ccw (solid dot) for three subtypes of border cells. **(E)** The border score of three subtypes of border cells. All data are represented as mean ± SEM. **(F)** The mean firing rate of three subtypes of border cells. **(G)** Layer distribution of three subtypes of border cells. **(H)** Polar plot showing the distribution of peak direction of head-direction tuning for four color-coded walls. Left to right: CW-, CCW- and regular border cells. **(I)** The summary of peak directions of head-direction tuning for three subtypes of border cells. **(J)** Comparison of mean vector length calculated in four separate walls for three subtypes of border cells **(K)** The ratio of CCW- and CW-border cells recorded in the right versus left hemisphere. See also **Figures S1-S4**.

To distinguish the relative composition of putative excitatory and inhibitory neurons, we classified all recorded cells from the S1 and V2 based on their spike widths and firing rates (**Figure S3D**). Putative fast-spiking interneurons were classified as cells with an average firing rate above 3 Hz and a peak-to-trough spike width below 300 µs (Peyrache et al., 2012). According to our criterion, 68.29% (n= 211/309) of recorded border cells were classified as regular-spiking (RS), 5.5% (n= 17/309) were classified as fast-spiking (FS) and 26.21% (n = 81/309) were unclassified (**Figure S3D**). FS border cells exhibited moderately lower border score than the other two types (**Figure S1E**, one-way ANOVA test, post hoc Bonferroni multiple comparisons test, Wall1: F(3) = 3.401, *p* = 0.035, RS versus FS: *p* = 0.06; FS versus Unclassified: *p* = 0.36; RS versus Unclassified: *p* = 0.44). Notably, all three classes of border cells consisted of both putative RS and FS cell types (**Figures S1, A-C** and **Figure S1F**).

There were no significant differences in border score (**Figure 1E**, one-way ANOVA test, F(3) = 1.03, *p* = 0.39, post hoc Bonferroni multiple comparisons test, *p* > 0.05) or mean firing rate (**Figure 1F**, one-way ANOVA test, F(3) = 1.03, *p* = 0.85, post hoc Bonferroni multiple comparisons test, *p* > 0.05) among CW-, CCW- and regular border cells. Although both CW- and CCW-border cells can be recorded in either hemisphere, they exhibited a distribution bias. Specifically, 57.17% (n = 48/84) of CW-border cells were located in the right hemisphere, whereas 63.01% (n = 46/73) of CCW-border cells were recorded from the left hemisphere. The ratios of CW- and CCW-border cells recorded in the right versus the left hemisphere were significantly different (**Figure 1K**, Pearson’s chi-squared test, *p* = 0.012).

To quantify the directional tuning of the three subtypes of border cells, we calculated the mean vector lengths along four individual walls of the arena. CW- and CCW-border cells showed strong tunings to the geometric walls, obtained higher mean vector length than regular border cells (**Figure 1J**, one-way ANOVA test, all with post hoc Bonferroni multiple comparisons test, Wall1: F(3) = 61.22, *p* < 1 × 10^-6^, CCW versus Regular: *p* < 1 × 10^-6^; CW versus Regular: *p* < 1 × 10^-6^; CCW versus CW: *p* = 0.19; Wall2: F(3) = 65.08, *p* < 1 × 10^-6^, CCW versus Regular: *p* < 1 × 10^- 6^; CW versus Regular: *p* < 1 × 10^-6^; CCW versus CW: 1.00; Wall3: F(3) = 66.95, *p* < 1 × 10^-6^, CCW versus Regular: *p* < 1 × 10^-6^; CW versus Regular: *p* < 1 × 10^-6^; CCW versus CW: *p* = 0.93; Wall4: F(3) = 78.74, *p* < 1 × 10^-6^, CCW versus Regular: *p* < 1 × 10^-6^; CW versus Regular: *p* < 1 × 10^-6^; CCW versus CW: *p* = 0.81), and showed preferred unidirectional firing patterns (**Figure 1H**). The summary of preferred direction in the head-direction tunings of CCW- and CW-border cells displayed a 90° step increase along four walls (**Figure 1I**). In contrast, regular border cells exhibited similar peak directions for two opposite walls (**Figure 1I**) as a consequence of symmetric firing in heading directions.

We also sought to identify putative monosynaptic connections (Bartho et al., 2004; Csicsvari et al., 1998; Latuske et al., 2015) for establishing the functional links between CW-, CCW-border cells and other spatially-tuned cells in the S1 and V2. We found a small number of CW- and CCW-border cells shared local excitatory or inhibitory (and sometimes reciprocal) connections to either simultaneously recorded regular border cells or between each other (**Figure S4**, 12/102 pairs; 11.76%). However, from 18 simultaneously recorded cell pairs, there were no identified putative monosynaptic connections between CW- or CCW-border cells and head-direction cells.

### CW- and CCW-Border Cells in Response to New Inside Wall

To verify whether CW- and CCW-border cells showed orientation-sensitive border responses, we recorded the activity of CW-border cells (*n* = 8) and CCW-border cells (*n* = 7) within a square enclosure in five successive trials (**Figures 2B** and **2D**, see also **Figures S5** and **S6**), starting with a baseline recording, continuing with the additional introduction of one discrete wall (orientation order: horizontal◊vertical◊diagonal), and ending with another baseline recording (**Figure 2A**). The newly inserted horizontal or vertical wall generated new border fields of CW- and CCW-border cells on both the proximal and distal sides of the wall insert. Notably, the newly evoked border fields displayed parallel head-direction tunings to those of the corresponding distal wall (**Figure 2C**), in contrast to the reverse head-direction tuning to that of the corresponding proximal wall (**Figures 2B** and **2D**). However, the preferred directions of newly evoked border fields on the same side of the new insert were opposite for CW- and CCW-border cells. For the diagonal inserts, the newly formed firing fields of CW-border cells only emerged during animal’s counterclockwise movement along either 45° or 225° direction (where 0° denotes the east direction). Additionally, the opposite phenomenon was found for CCW-border cells. Together, CW- and CCW-border cells showed opposite directional tunings along the same side of the new insert (**Figure 2E**), and their peak directions were nearly 180° apart (**Figures 2F** and **2G**). Additionally, they shared similar mean vector length along the two sides of the newly inserted wall (**Figures 2H** and **2I**, two-sided Wilcoxon signed rank test, CW-border: *n* = 8, H, Z = 1.26 and *p* = 0.21; D, Z = 1.68 and *p* = 0.09; V, Z = 0.70 and *p* = 0.48; CCW-border: *n* = 7, H, Z = 1.21 and *p* = 0.23; D, Z = 0 and *p* = 1.00; V, Z = 0.94 and *p* = 0.35).

**Figure 2.**
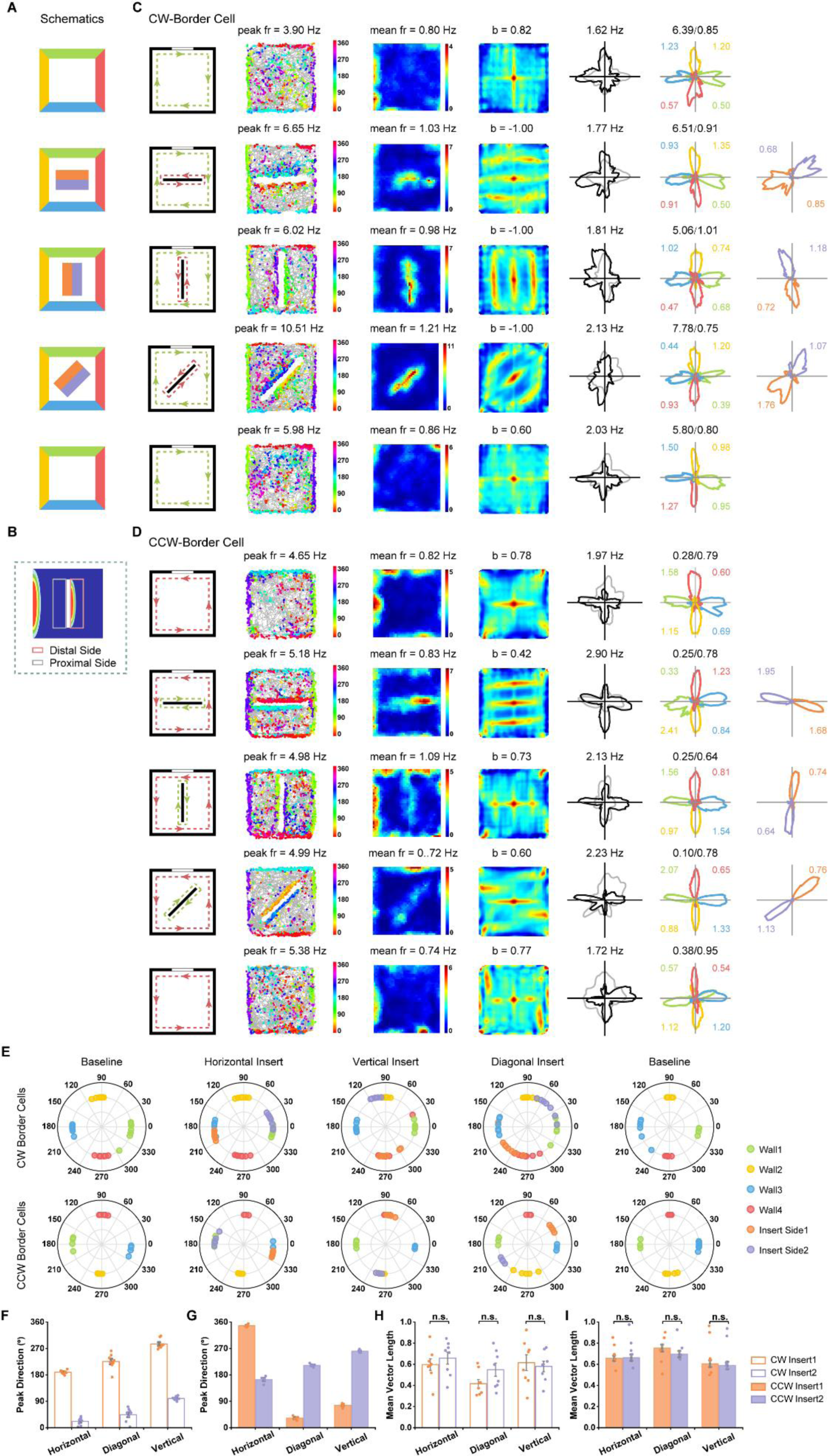
CCW- and CW-Border Cells in Response to New Inside Wall. **(A)** Diagram of experimental procedure with the baseline, inserted wall (order: horizontal→vertical→diagonal), and second baseline conditions. **(B)** Schematic showing spatial responses to the insertion of a new wall (distal vs. proximal side) relative to the original border responses. **(C**, **D**) Spatial responses of representative rat cortical CW- and CCW- border cells, respectively, in the baseline, inserted wall (order: horizontal→vertical→diagonal), and second baseline conditions. Notations and symbols are similar to Figure 1. (**E**) Distribution of peak directions for four color-coded sides in five consecutive conditions. Note the peak direction of head-directional tuning for the two sides of the insert were parallel to the orientation of the new discrete wall. (**F**, **G**) Summary of peak directions along the two sides of the new insert for CW- and CCW-border cells, respectively. The peak directions along the two sides of the new insert varied from different inserts. The CW- and CCW-border cells exhibited opposite directional tuning along the same side of the insert. (**H**, **I**) Distribution of mean vector length of border cells showing that the head direction maintained the tuning properties along the two sides of the insert. See also **Figures S5 and S6**.

### Spatial Responses of CW- and CCW-Border Cells in Environments with Two Geometric Borders

To confirm asymmetric rotational firing and border specificity in CW- and CCW-border cells, we introduced two geometric borders within the running enclosure. We first placed a small 50 × 50 × 30 cm^3^ (L × W × H) square box in the center of a larger 150 × 150 cm^2^ arena, creating an outer ring and an inner ring two separate geometric borders. Rats could not climb over the small box during foraging. Within the outer ring, CW-border cells fired only when animals moved along the boundary in a clockwise direction (**Figure 3A**). In contrast, within the inner ring, the same CW- border cells only fired in the counterclockwise direction, displaying egocentric border firing specificity. The transformation from a CW-border cell within the outer ring into a CCW-border cell along the inner ring, therefore, confirmed the egocentric coding of border firing patterns (**Figures 3A** and **S7**). Similar patterns were also found in CCW-border cells (**Figures 3B** and **S8**). The distribution of preferred direction in head-direction tunings along the four walls of the outer ring was similar to that of the baseline but opposite to that of the inner ring (**Figure 3C**). Additionally, the distribution of peaks had four dominant modes, showing 90° step increases in the peak directions for CW- and CCW-border cells (**Figures 3D** and **3E**, respectively). For both CW- and CCW-border cells, the CW/CCW firing rate ratio displayed an inverse pattern between the outer and inner rings despite the similar CW/CCW ratios during occupancy (two-sided Wilcoxon signed rank test, CW-border: *n* = 5, ratio in firing rate, Z = 2.02 and *p* = 0.043; ratio in occupancy, Z = 0.14 and *p* = 0.89; CCW-border: *n* = 7, ratio in firing rate, Z = 2.37 and *p* = 0.018; ratio in occupancy, Z = 1.52 and *p* = 0.13), suggesting that the firing asymmetry was not caused by the difference in animal’s dwell time (**Figures 3F** and **3G**). Although the head-direction tunings of CW- and CCW-border cells were inverted between the outer ring and inner ring, their mean vector lengths maintained stable between two rings (**Figures 3H** and **3I**). Similar asymmetric coding and border sensitivity were also detected in another spatial apparatus with three geometric borders (**Figure S9**). The rings displayed an alternating pattern, where the outer and inner rings shared one selectivity, and the middle ring took the other. This switch between CW- and CCW-border firing patterns within the same asymmetric border cells provides strong evidence of egocentric spatial coding.

**Figure 3.**
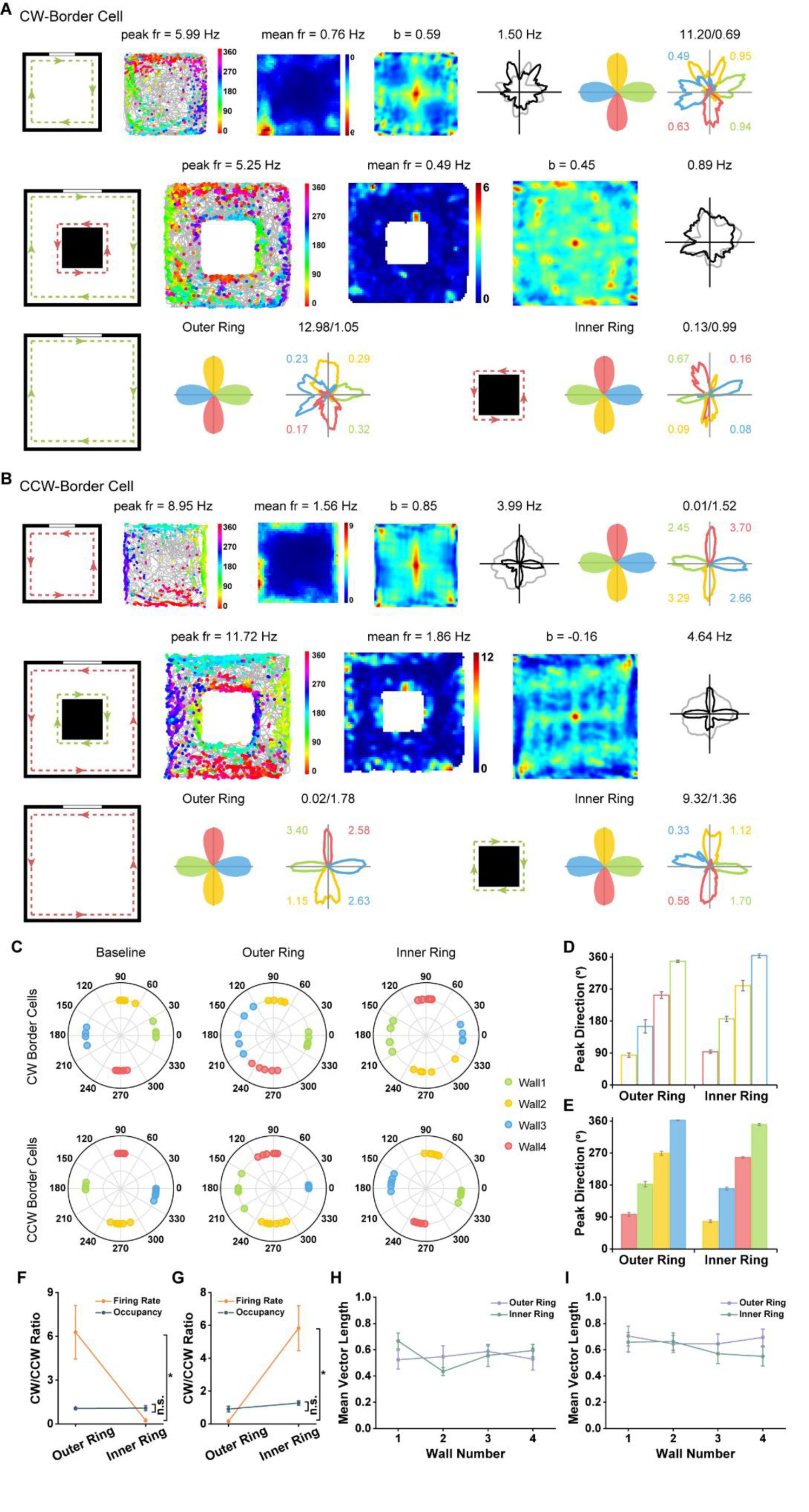
Spatial Responses of CCW- and CW-Border Cells in Environment with Two Geometric Borders. (**A**, **B**) Spatial responses of representative CW- and CCW-border cells, respectively, in running boxes with two geometric borders. A small 50 × 50 × 30 cm^3^ square box was placed in the center of a large 150 ×150 cm^2^ box which created an outer ring and an inner ring of geometric borders. The rat could not climb over the box. Notations and symbols are similar to **Figure 1**. (**C**) Distribution of peak directions of head direction tuning for four color-coded walls in baseline and manipulation conditions. For outer and inner rings in the same box, the head direction of the four sides showed opposite tuning properties. (**D**, **E**) Summary of the peaks of directional tuning for the outer ring and inner ring for CW- and CCW-border cells, respectively. (**F**, **G**) CW/CCW ratio in firing rate (red) and occupancy (blue) for outer ring and inner ring for CW- and CCW- border cells, respectively. (**H**, **I**) Mean vector length of directional responses for four sides of the outer ring and inner ring for CW- and CCW- border cells, respectively. Note that the average value of mean vector length for each side was higher than 0.4, a typical value considered to indicate tuning to head direction. See also **Figures S7-S9**.

### CW- and CCW-border Cells Are Anchored to Local Environmental Borders

If the properties of CW- and CCW-border cells were stably anchored to the layout of the local borders rather than distal cues within a larger environment, the rotation of the local environment would cause change in border tuning. To address this possibility, we conducted four consecutive experimental manipulations: starting with a baseline recording session, then one session with counterclockwise 45° rotation of the square box, followed by another session with clockwise 45° rotation of the square box, and a last baseline recording session (**Figure 4** and **Figures S10 and S11**). Together with CW- or CCW-border cells (**Figures 4A** and **4C**), head-direction cells were also simultaneously recorded and detected from the same tetrode (**Figures 4B** and **4D**). Following the local cue card rotation, there was no significant change in either border score, CW/CCW firing rate ratio, or mean vector length along four single walls among the identified eight CW- and six CCW-border cells (**Figures 4E-4G**). In contrast, the preferred head-direction tunings of CW- and CCW-border cells rotated coherently with the local cue card (**Figures 4H** and **4I**). Notably, simultaneously recorded head-direction cells also showed a significant shift in response to the rotation of local boundaries (**Figures 4B** and **4D**), as verified by the strong correlation in angular peak direction shift between head-direction tuning of CW- and CCW-border cells and head-direction cells (**Figure 4J**, HD-Wall1, *r* = 0.92, *p* = 1.67×10^-5^; HD-Wall2, *r* = 0.91, *p* = 2.12×10^-4^; HD-Wall3, *r* = 0.91, *p* = 7.25×10^-6^; HD-Wall4, *r* = 0.93, *p* = 1.07×10^-6^). The CW/CCW ratio in firing rate remained almost unchanged (two-sided Wilcoxon signed rank test, CW-border: *n* = 8, B-CCW45, Z = 1.54 and *p* = 0.12; B-CW45, Z = 1.4 and *p* = 0.16; B-B’, Z = 0.28 and *p* = 0.78; CCW-border: *n* = 6, B-CCW45, Z = 0.73 and *p* = 0.46; B-CW45, Z = 0.31 and *p* = 0.75; B-B’, Z = 0.73 and *p* = 0.46). Besides, border scores were not significantly different across cue manipulation conditions for both CW- and CCW-border cells (two-sided Wilcoxon signed rank test, n.s., not significant, CW-border: *n* = 8, B-CCW45, Z = 0.49 and *p* = 0.62; B- CW45, Z = 0.98 and *p* = 0.33; B-B’, Z = 0 and *p* = 1.00; CCW-border: *n* = 6, B-CCW45, Z = 0.11 and *p* = 0.92; B-CW45, Z = 0.73 and *p* = 0.46; B-B’, Z = 1.15 and *p* = 0.25). Additionally, we examined CW- and CCW-border cells that fired along either one or two walls (**Figure S11C**) in response to a 90° (clockwise and counterclockwise) rotation of the cue card. As expected, the directional tuning of the conjunctive CW- and CCW-border cells rotated coherently with the local borders of the open field. Together, these results indicate that egocentric CW-and CCW-border cells respond to local boundaries and exhibit a significant shift in rotated environments, suggesting that two independent streams of egocentric and allocentric reference frames coexist within the rat S1 and V2.

**Figure 4.**
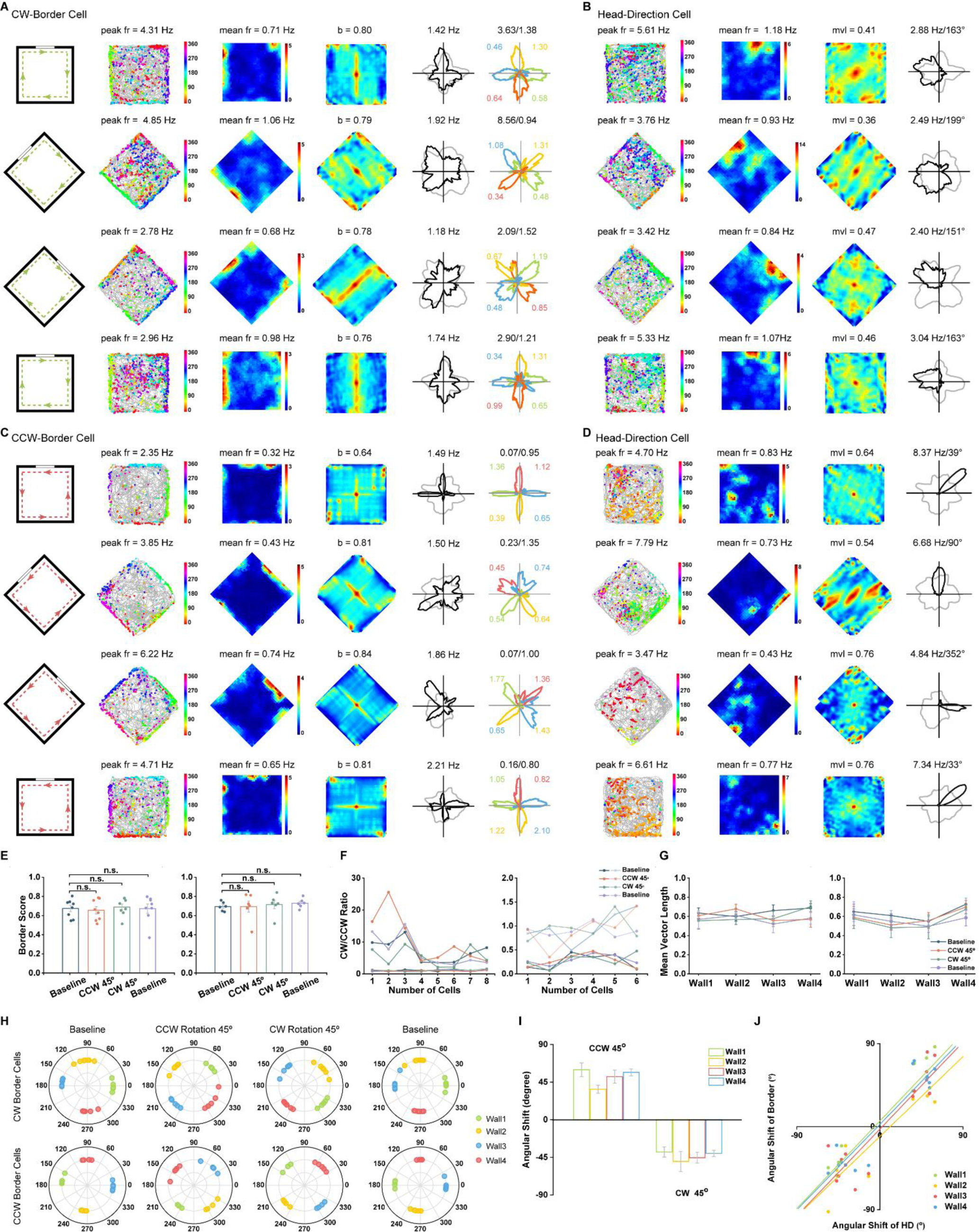
Visual Landmark Control of CCW- and CW-Border Cells. (**A**, **B**) Spatial responses of a representative CW-border cell and a simultaneously recorded V2 head-direction cell, respectively, in the baseline (B), counterclockwise rotation 45°(CCW45), counterclockwise rotation 45°(CW45) and second baseline conditions (B’). Notations and symbols are similar to **Figure 1**. (**C**, **D**) Same as (**A**, **B**) except for a representative CCW-border cell and a simultaneously recorded head direction cell. **(E)** Border score was not significantly different across cue manipulation conditions for both CW- (left) and CCW- (right) border cells. **(F)** The CW/CCW firing rate ratio remained nearly unchanged in different cue rotation experiments for both CW- and CCW-border cells. Curves in light and dark color represent time_cw/ccw and fr_cw/ccw, respectively. **(G)** Distribution of mean vector length of the head directional responses along the four walls for both CW- and CCW- border cells. Head directionality was well preserved under the different cue manipulation conditions. **(H)** Distribution of peak direction of head-direction tuning for four color-coded walls in the baseline and rotation conditions. **(I)** Angular shift of corresponding peak directions of head-direction tuning for four color-coded walls in the clockwise and counterclockwise 45° rotation conditions. **(J)** Correlation of angular peak direction shift between head-direction tuning of CW- and CCW-border cells along four single walls and head-direction cells. See also **Figures S10 and S11**.

### CW- and CCW-Border Cells Maintained Egocentric Tunings in Darkness

To determine whether visual input was required to retain the rotation-selective asymmetry for CW- and CCW-border cells, we further recorded the spike activity of CW- and CCW-border cells during the sequential light-dark-light (L-D-L’) sessions (**Figures 5A** and **5B**; see also **Figures S12** and **S13**). The preferred direction of head-direction tuning along the four walls persisted during the L-D-L’ sessions (**Figure 5C**). The CW/CCW ratio in firing rate remained nearly unchanged during these sessions for both CW- and CCW-border cells (**Figure 5D**) (two-sided Wilcoxon signed rank test, CW-border: *n* = 11, L-D, Z = 0.53 and *p* = 0.59, D-L’, Z = 0.36 and *p* = 0.72; L-L’, Z = 1.25 and *p* = 0.21; CCW-Border: *n* = 8, L-D, Z = 1.72 and *p* = 0.09, D-L’, Z = 1.84 and *p* = 0.07; L-L’, Z = 0 and *p*= 1.00). Darkness had no significant influence on these S1 or V2 neurons’ mean firing rates (**Figure 5E**, CW-border: L-D, Z = 0.09 and *p* = 0.93; D-L’, Z = 0.27 and *p* = 0.79; L-L’, Z = 0.45 and *p* = 0.66; CCW-Border: L-D, Z = 0.14 and *p* = 0.89; D-L’, Z = 1.26 and *p* = 0.21; L-L’, Z = 0.42 and *p* = 0.67) or border scores (**Figure 5F**, CW-border: L-D, Z = 0.27 and *p* = 0.79; D-L’, Z = 0.31 and *p* = 0.76; L-L’, Z = 0.15 and *p* = 0.88; CCW-Border: L- D, Z = 0.56 and *p* = 0.58; D-L’, Z = 0.42 and *p* = 0.67; L-L’, Z = 0 and *p* = 1.00). Meanwhile, the switch from the light condition to darkness did not affect their direction tunings (**Figure 5G**). Overall, CW- and CCW-border cells sustained their rotation-selective asymmetry in the absence of visual input, suggesting that egocentric tuning properties of these CW- and CCW-border cells are not dependent on allocentric cues in spatial navigation.

**Figure 5.**
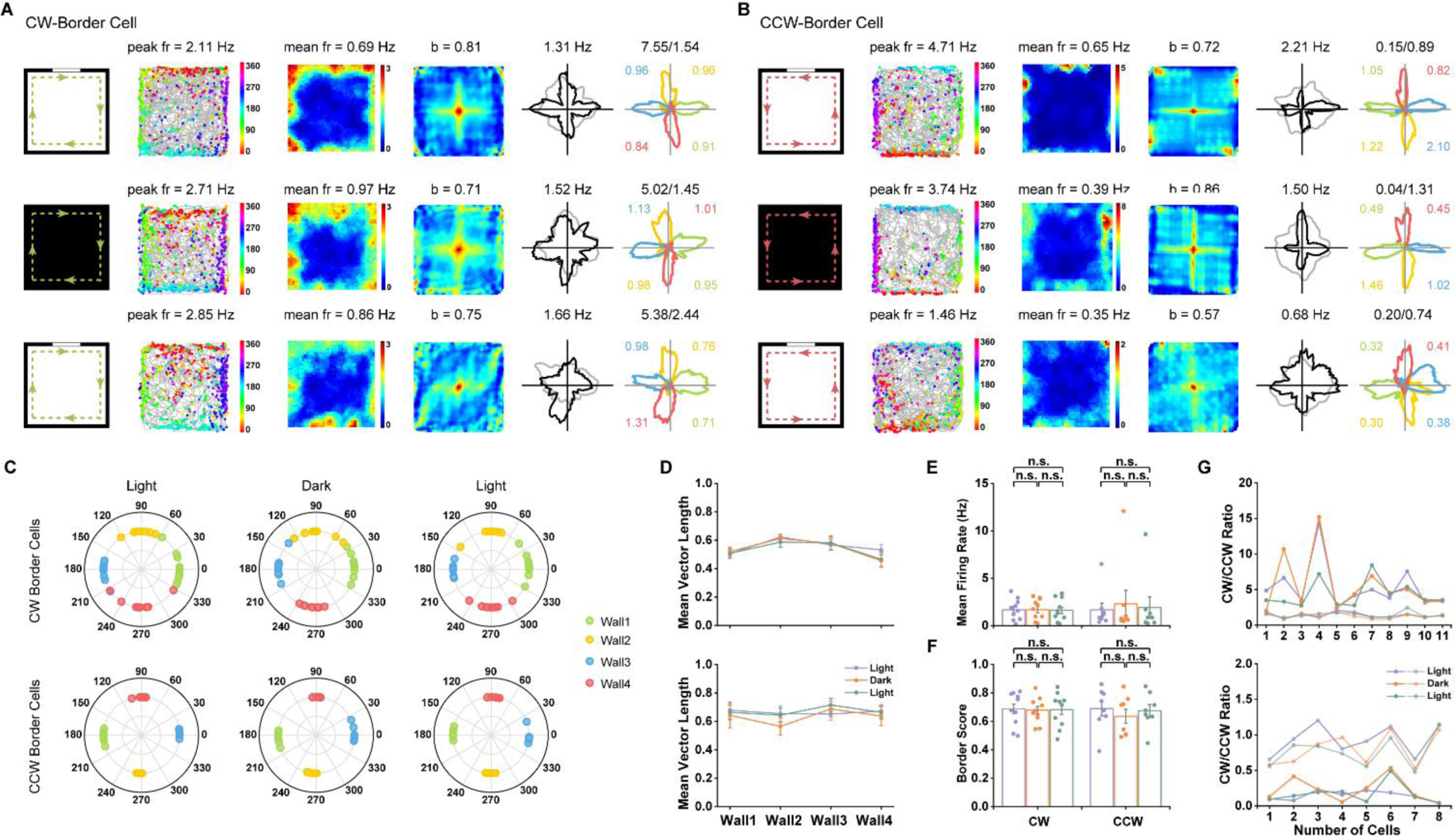
Somatosensory CCW- and CW-border Cells Persist in Darkness. (**A**, **B**) Spatial responses of representative cortical CW- and CCW-border cells, respectively, in the light (L), dark (D) and light (L’) conditions. Notations and symbols are similar to Figure 1. **(C)** Distribution of peak directions for four color-coded sides in L-D-L’ conditions. **(D)** Mean vector length along the four sides of the square enclosure indicated that head-directional tuning persisted in the darkness for both CW- (top) and CCW- (bottom) border cells. **(E)** Mean firing rate was not significantly different between darkness and light conditions. **(F)** Border score was not significantly different between darkness and light conditions. **(G)** The CW/CCW firing rate ratio remained almost unchanged in total darkness compared to the light condition for both CW- (top) and CCW- (bottom) border cells. Curves in light color represents time_cw/ccw and curves in dark color represents fr_cw/ccw. See also **Figures S12 and S13**.

### Spatial Responses of CW- and CCW-Border Cells to Environmental Wall and Geometry

Next, we investigated whether CW- and CCW-border cells were specialized to respond to environmental borders rather than physical walls, or specifically, whether the egocentric border cells continued to evoke border-responsive asymmetric rotational firing in a wall-less recording enclosure. To do so, we removed all four external walls of the recording arena, leaving a 50-cm high drop-edge to the floor (**Figures 6A** and **6B**). The animals ran on the open running surface without direct somatosensory information of physical walls. Both CW- and CCW-border cells lost their head-direction tuning along four edges in the wall-less elevated platform (**Figure 6C**; see also **Figures S14** and **S15**), and the preferred direction along four sides of running arena was no longer discretized (**Figure 6D**). Furthermore, the mean vector length along four sides of running arena also decreased after wall removal (**Figure 6E**), yet there was no significant change in mean firing rate after wall removal (**Figure 6F**). Consequently, the CW/CCW ratio in firing rate decreased for both CW- and CCW-border cells (**Figure 6H**,CW-border: B-E, Z = 2.80 and *p* = 0.005; E-B’, Z = 2.80 and *p* = 0.005; B-B’, Z = 1.58 and *p* = 0.11; CCW-Border: B-E, Z = 2.67 and *p* = 0.008; E-B’, Z = 2.67 and *p* = 0.008; B-B’, Z = 0.18 and *p* = 0.86). Consistent with the border score decrease in entorhinal border cells (Solstad et al., 2008), egocentric CW- and CCW-border cells in the rat S1 and V2 also decreased their border scores in the wall-less elevated platform (**Figure 6G**, two-sided Wilcoxon signed rank test, CW-border: *n* = 10, B-E, Z = 2.70 and *p* = 0.007; E-B’, Z = 2.60 and *p* = 0.009; B-B’, Z = 0.77 and *p* = 0.44; CCW-Border: *n* = 9, B-E, Z = 2.67 and *p* = 0.008; E-B’, Z = 2.67 and *p* = 0.008; B-B’, Z = 0.42 and *p* = 0.68). Together, these results indicated that the physical walls are required to maintain the egocentric border signals.

**Figure 6.**
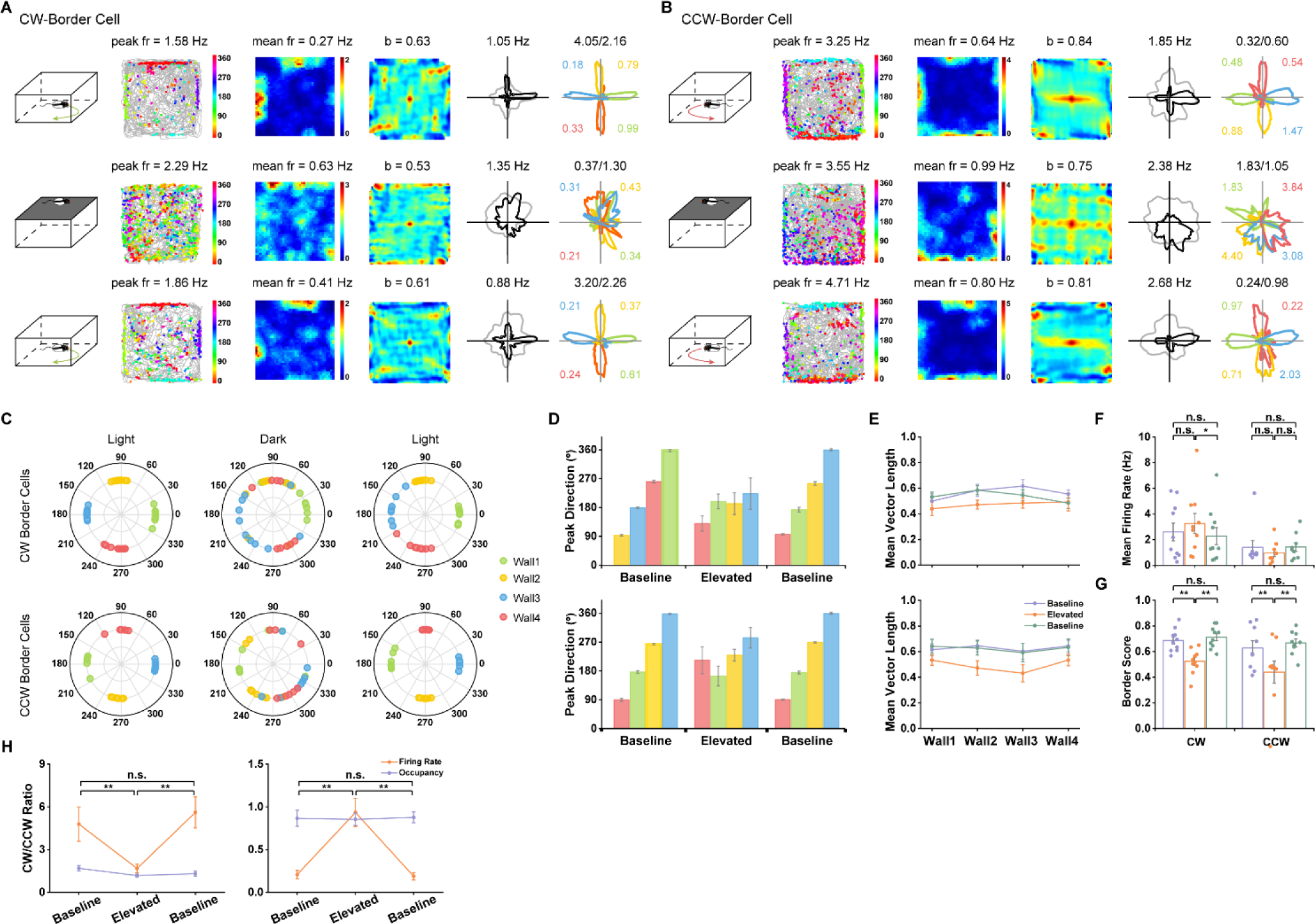
CCW- and CW-border cells Requires Geometric Physical Borders. (**A**, **B**) Spatial responses of representative cortical CW- and CCW-border cells, respectively, in the baseline (B), elevated platform without walls (E) and baseline (B’) sessions. Notations and symbols are similar to **Figure 1**. **(C)** Distribution of peak directions for four color-coded sides in the baseline, elevated platform without walls, and baseline sessions. Note the disrupted regularity of the 90° step increase of peak directions along the four borders in the elevated platform. **(D)** The summary of peak directions of head-direction tuning for CW- (top) and CCW- (bottom) border cells. The peak directions of head direction tuning in the elevated platform lost their unidirectional firing properties compared to those in the baseline sessions. **(E)** Mean vector length along the four sides of the elevated platform showed that the head directional tuning deteriorated in the elevated wall-less platform for both CW- (top) and CCW- (bottom) border cells, respectively. **(F)** Comparison of mean firing rates of CW- and CCW-border cells between baseline and elevated platform. **(G)** Border score was significantly reduced in elevated wall-less platform for both CW- and CCW-border cells. **(H)** CW- (left) and CCW- (right) border cells didn’t maintain CW/CCW firing rate ratio in circumstances without physical borders. See also **Figures S14 and S15**.

**Figure 7.**
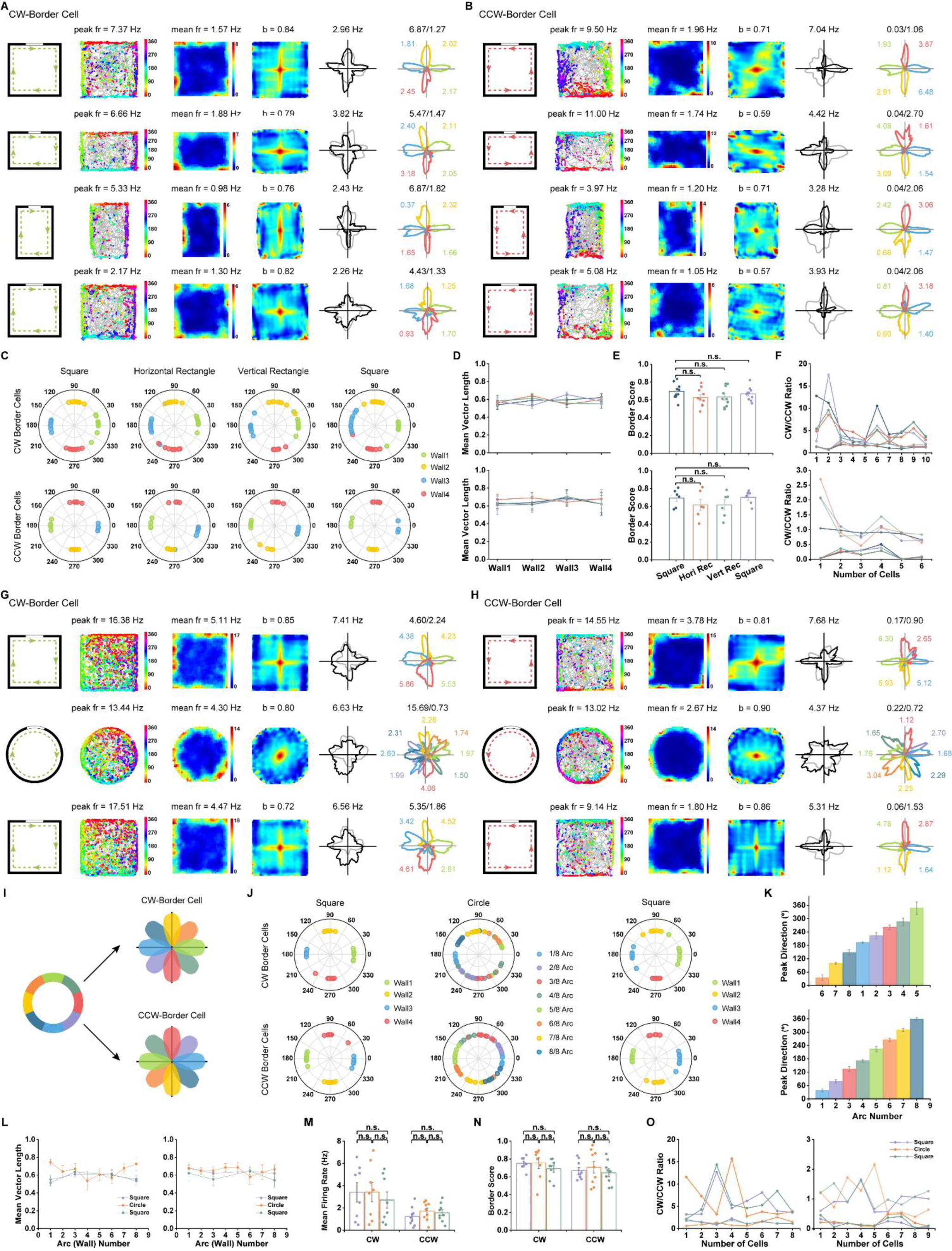
CCW- and CW-Border Cells Preserve Unidirectional Firing Patterns in Rectangular and Circular Environments. (**A**, **B**) Spatial responses of representative cortical CW- and CCW-border cells, respectively, in the 100 × 100 cm^2^ square enclosure (S), 100 × 70 cm^2^ horizontal rectangle (R_h_), 70 × 100 cm^2^ vertical rectangle (R_v_) and back in the 100 × 100 cm^2^ square enclosure (S’). Notations and symbols are similar to Figure 1. **(C)** Distribution of peak directions for four color-coded sides in the baseline, elevated wall-less platform and baseline sessions. The regularity of the 90° step increase of peak directions along the four borders persisted in the rectangle. **(D)** Mean vector length showed that the head directional tuning was maintained in the rectangles for both CW- (top) and CCW- (bottom) border cells. **(E)** Border scores of CW- (top) and CCW- (bottom) border cells showed no significant difference between square and rectangle boxes. **(F)** CW- (top) and CCW- (bottom) border cells maintained their CW/CCW firing rate ratios in different running boxes. (**G**, **H**) Spatial responses of representative cortical CW- and CCW-border cells, respectively, in the square (S), circle (C) and back in the square (S’) enclosure. **(I)** Schematic diagram showing the representative responses of eight divided arcs in circular environment for CW- and CCW-border cells. **(J)** Distribution of peak directions for four color-coded sides in the square enclosure and eight radian in the circular arena. **(K)** Summary of peak directions in eight arcs of circular boxes for CW- (top) and CCW- (bottom) border cells. The peak directions gradually changed and CW- and CCW-border cells exhibited opposite directional tuning in the same arc. **(L)** Mean vector length along four walls in the square enclosure and along eight arcs in the circular arena for CW- and CCW-border cells. Mean vector length showed that both the CW- (orange) and CCW- (blue) border cells maintained their head direction tuning properties in the circle enclosure. **(M)** Mean firing rate showed no significant change between the square and circle enclosures. **(N)** Border score was not statistically different between distinct geometric shapes. **(O)** CW- and CCW-border cells maintained CW/CCW ratio in firing rate in different shapes of boxes. See also **Figures S16**-**S19**.

Furthermore, we determined whether the activity of egocentric CW- and CCW-border cells was influenced by the environmental geometry. We inserted a 1-m long and 50-cm high plate into the center of the 1-m square box either in parallel or orthogonal to the original cue card, stretching the 1 × 1 m^2^ square into a 0.7 × 1 m^2^ (R_h_) or 1 × 0.7 m^2^ (R_v_) rectangle. The firing fields of CW- and CCW-border cells were insensitive to the geometric shapes (**Figures 7A** and **7B**, see also **Figures S16** and **S17**). CW- and CCW-border cells yielded opposite head-direction tunings along the four walls, which were aligned with the orientation of the walls in the stretched rectangular enclosure (**Figure 7C**) with high mean vector length along all four borders (**Figure 7**). There was no significant difference in border scores (**Figure 7E**, two-sided Wilcoxon signed rank test, CW-border: *n* = 10, S-R_h_, Z = 1.88 and *p* = 0.06; S-R_v_, Z = 1.68 and *p* = 0.10; S-S’, Z = 0.87 and *p*= 0.39; CCW-Border: *n* = 6, S-R_h_, Z = 0.94 and *p* = 0.35; S-R_v_, Z = 1.78 and *p* = 0.08; S-S’, Z = 0.11 and *p* = 0.92) or CW/CCW firing rate ratio (**Figure 7F**, CW-border: S-R_h_, Z = 1.88 and *p* = 0.06; S-R_v_, Z = 1.88 and *p* = 0.06; S-S’, Z = 1.27 and *p* = 0.20; CCW-Border: S- R_h_, Z = 0.73 and *p* = 0.46; S-R_v_, Z = 1.15 and *p* = 0.25; S-S’, Z = 0.31 and *p* = 0.75).

In addition, we tested the effect of environmental geometry on the orientation-specific tuning of CW- and CCW-border cells in square-circle-square (S-C-S’) recording sessions. As a result of gradual change of geometric borders in the circular environment, egocentric border cells showed directionally uniform tuning (**Figures 7G** and **7H**, see also **Figures S18** and **S19**). To quantify the direction tuning in the circular arena, we evenly divided the full 360° into eight sectors and calculated the corresponding head-direction tuning curves for each arc (**Figure 7I**). The preferred direction of head-direction tuning showed a gradual angular shift (**Figures 7J** and **7K**) with high mean vector length value (**Figure 7L**). Notably, the geometric shape did not change the mean firing rate (**Figure 7M**, two-sided Wilcoxon signed rank test, CW-border: *n* = 8, S-C, Z = 0 and *p* = 1.00; C-S’, Z = 0.70 and *p* = 0.48; S-S’, Z = 0.77 and *p* = 0.44; CCW-border: *n* = 10, S-C, Z = 1.48 and *p* = 0.14; C-S’, Z = 0.41 and *p* = 0.68; S-S’, Z = 1.84 and *p* = 0.07), border score (**Figure 7N**, CW-border: S-C, Z = 0.42 and *p* = 0.67; C-S’, Z = 1.26 and *p* = 0.21; S-S’, Z = 1.26 and *p* = 0.21; CCW-border: S-C, Z = 0.77 and *P* = 0.44; C-S’, Z = 0.33 and *p* = 0.33; S-S’, Z = 0.65 and *p* = 0.52) or CW/CCW firing rate ratio (**Figure 7O**, CW-border: S-C, Z = 0.14 and *p* = 0.89; C-S’, Z = 0 and *p* = 1.00; S-S’, Z = 0.42 and *p* = 0.67; CCW-border: S-C, Z = 1.48 and *p* = 0.14; C-S’, Z = 0.41 and *p* = 0.68; S-S’, Z = 1.48 and *p* = 0.14).

Finally, we tested whether CW- and CCW-border cells preserved their egocentric properties in novel environments in a familiar-novel-familiar (A-B-A’) setting (**Figures S20-23**). CW- and CCW-border cells sustained their preferred direction tunings along all four borders in new environment (**Figures S20C** and **S20G**). There was no change between the novel and familiar environments in their mean firing rate (**Figure S20E**, two-sided Wilcoxon signed rank test, CW- border: *n* = 6, A-B, Z = 0.94 and *p* = 0.35; B-A’, Z = 1.15 and *p* = 0.25; A-A’, Z = 0.31 and *p* = 0.75; CCW-Border: *n* = 6, A-B, Z = 1.78 and *p* = 0.08; B-A’, Z = 1.36 and *p* = 0.17; A-A’, Z = 0.52 and *p* = 0.60), border score (**Figure S20F**, CW-border: A-B, Z = 0.73 and *p* = 0.46; B-A’, Z = 0.94 and *p* = 0.35; A-A’, Z = 0.94 and *p* = 0.35; CCW-Border: A-B, Z = 0.11 and *p* = 0.92; B-A’, Z = 1.36 and *p* = 0.17; A-A’, Z = 0.94 and *p* = 0.35) or CW/CCW ratio in firing rate (**Figure S20G**, CW-border: A-B, Z = 0.31 and *p* = 0.75; B-A’, Z = 0.52 and *p* = 0.60; A-A’, Z = 0.73 and *p* = 0.46; CCW-Border: A-B, Z = 1.36 and *p* = 0.17; B-A’, Z = 0.73 and *p* = 0.46; A-A’, Z = 0.94 and *p* = 0.35). These results demonstrated that these egocentric border cells are insensitive to environmental geometry and novel circumstances.

### CW- and CCW-Border Cells in Response to Whisker Trimming

Whiskers are an important component for the somatosensory system in rodent navigation. To evaluate how whiskers affected the border firing patterns in the rat S1 or V2 neurons, we also recorded their spiking activities before and after whisker trimming (**Figures S23-25**). For CW- border cells, whiskers on the left side were first trimmed and then after one recording session, whiskers on the right side were trimmed (**Figure S23A**). For CCW-border cells, the order was reversed such that the whiskers on the right side were trimmed before those on the left side (**Figure S23B**). There was no significant change in mean firing rate (**Figure S23E**, two-sided Wilcoxon signed rank test, CW-border: *n* = 9, W-O, Z = 0.85 and *p* = 0.40; O-N, Z = 1.01 and *p* = 0.31; W-N, Z = 1.35 and *p* = 0.18; CCW-Border: *n* = 7, W-O, Z = 0.53 and *p* = 0.59; O-N, Z = 0.18 and *p* = 0.86; W-N, Z = 0.18 and *p* = 0.86), border score (**Figure S23F**, CW-border: W-O, Z = 0.42 and *p* = 0.68; O-N, Z = 1.36 and *p* = 0.17; W-N, Z = 1.24 and *p* = 0.21; CCW-Border: W-O, Z = 0.68 and *p* = 0.50; O-N, Z = 0.17 and *p* = 0.87; W-N, Z = 0.68 and *p* = 0.50) or CW/CCW ratio in firing rate (**Figure S23G**, CW-border: W-O, Z = 1.24 and *p* = 0.21; O-N, Z = 0.65 and *p* = 0.52; W-N, Z = 1.48 and *p* = 0.14; CCW-Border: W-O, Z = 0.30 and *p* = 1.00; O-N, Z = 0.65 and *p* = 0.87; W-N, Z = 0.30 and *p* = 0.50). The preferred direction of head-direction tunings along four walls remained unchanged (**Figure S23C**), with well-preserved head directionality along four borders (**Figure S23D**) after (either one- or two-side) trimming animals’ whiskers. Together, these findings suggest that there was no effect of the whisker deprivation on the firing properties and spatial responses of both CW- and CCW-border cells in both S1 and V2.

## DISUCSSION

Vector coding (such as distance and direction) in spatial cognition is common for neurons within and around the hippocampal formation, far from the sensory periphery (Bicanski and Burgess, 2020). The spatial receptive fields (“vector codes”) of these neurons, are often expressed in the allocentric coordinates, such as in the cases of boundary vector cells (Lever et al., 2009), border cells (Solstad et al., 2008), landmark vector cells (Deshmukh and Knierim, 2013), object vector cells (Hoydal et al., 2019), visual cue cells (Kinkhabwala et al., 2020), human anchor cells (Kunz et al., 2021), border-elicited theta oscillation (Stangl et al., 2021), primate border cells (Killian et al., 2012) and vector trace cells (Poulter et al., 2021). Recently, egocentric representations of space have been reported in the PPC, RSC, lateral EC (LEC), postrhincal cortex, postsubiculum and dorsomedial striatum (Gofman et al., 2019; LaChance et al., 2019; Peyrache et al., 2017; Wang et al., 2018). However, to our best knowledge, such signals have never been found in the sensory cortices. Our findings about egocentric border cells in the rat S1 and V2 are consistent with the hypothesis that egocentric representations of reference frame are coded across distributed brain regions beyond the classical hippocampal formation. Specifically, egocentric border cells in the rat S1 and V2 discharge unidirectionally when animals move along the border of the local environment in a rotation-specific CW or CCW manner. Therefore, our results support they hypothesis that egocentric and allocentric spatial processing is functionally coupled in these brain areas. The co-existence of allocentric and egocentric border cells accommodates an egocentric × allocentric conjunctive coding strategy utilizing both reference frames within the somatosensory and visual cortices. An important question remains: how are egocentric and allocentric representations co-represented across a broad range of brain areas during spatial navigation, and how do they interact?

In the literature, egocentric and allocentric reference frames have been found in RSC and PPC neurons. The RSC and PPC are interconnected with the hippocampus, subiculum, perihinal, postrhinal, and entorhinal cortices, and also have projections to the sensory cortices, such as the V1, V2 and M2 (Whitlock et al., 2012). Additionally, the RSC and sensory cortices have dense reciprocal projections between their egocentric coordinate system and the allocentric-dominant hippocampal-entorhinal system. A recent report has suggested dissociation between the allocentric reference frame of the MEC and the egocentric reference frame of LEC (Wang et al., 2018). Furthermore, it was shown that RSC border cells specifically encode walls, but not objects, and are sensitive to the animal’s direction to nearby borders (Alexander et al., 2020), suggesting that the egocentric RSC representation of space is generated independently from visual or whisker sensation but is affected by the MEC input that contains allocentric spatial information (van Wijngaarden et al., 2020). Our results indicate that S1 and V2 egocentric border cells maintain their egocentric properties in the absence of whisker stimulation or light condition. Therefore, we envision that these primary and secondary sensory cortices, with possible involvement of the motor cortex, engage in allocentric and egocentric spatial coding in a “sensory-memory-motor” loop within a much larger spatial navigation brain network (Peyrache and Duszkiewicz, 2021). First, sensory cortices convert sensory information into two independent or conjunctive coordinate systems, and then transform the information into a coherent “cognitive map” representation of space. Second, hippocampal-neocortical interactions transform the integrated spatial information into motor output. It is likely that the somatosensory egocentric representation of space is generated from the allocentric spatial map originating from the hippocampal-entorhinal network because of their anatomic interconnections. The pure egocentric CW and CCW-border cells identified in the somatosensory and visual cortices might interact with common postsynaptic targets within the hippocampal-entorhinal network to encode allocentric × egocentric conjunctive spatial signals. These spatial signals might be further used depending on specific locomotion or navigation scenarios (e.g., naturalistic environment vs. virtual reality vs. simulated imagery).

The remapping of neurons’ spatial receptive fields in response to a changing environment, such as in response to alterations of the size or shape of the environment or in response to the addition of walls, reflects the transformation of those neurons’ reference frame. Egocentric border representations would change according to the coordinate such transformations, which would inform the upstream or downstream neurons about the position of external landmarks in a viewpoint-dependent manner. It has been previously shown that the border cells of the MEC, as well as the boundary vector cells of the RSC and dorsal subiculum, share similar context- invariant tuning (Alexander et al., 2020; Hartley and Lever, 2014; Lever et al., 2009; Solstad et al., 2008). In the rat S1 and V2, the switch between the CW- and CCW-border cells with respect to a new wall inside the open arena further confirms their egocentric nature of spatial coding. Additionally, the CW- and CCW-border fields are relatively invariant to the environmental geometry, and maintain their preferred directional tunings in rotated, expanded, elevated, and novel arenas. Importantly, these egocentric sensory cortical border cells are independent on the visual input (light vs. darkness) or whisker stimulation.

The prevalence of egocentric sensory cortical CW- and CCW-border cells, as well as their presence across multiple cortical layers, may suggest a universal or common computational function in sensory cortices. One hypothesis is that these primary or secondary sensory cortices receive egocentric signals from the RSC or PPC, as well as traditional direction-selective input from downstream structures (such as the sensory thalamus or V1); integration of these two sources of signals may contribute to such rotation-selective tuning asymmetry. However, a causal circuit dissection based on optogenetic inactivation of specific circuits is still required to test this hypothesis. Recent discoveries suggest that vector-based egocentric spatial representations emerge from a distributed cortical-subcortical network (Alexander et al., 2020). A new theory proposal, also known as the “Thousand Brains Theory” (Hawkins et al., 2018), suggests that the brain’s model of the world is built using map-like reference frames. In this theory, the neocortex stores thousands of reference frames for the purpose of spatial navigation and goal-directed planning (and thereby thinking), forming the basis for a broad range of brain functions.

To date, several computational models that perform egocentric-to-allocentric transformation have been proposed for spatial memory and navigation (Bicanski and Burgess, 2020; Byrne and Becker, 2008; Byrne et al., 2007; Madl et al., 2015; Rolls and Mills, 2019)). Specifically, neurons that are tuned for both allocentric head direction and egocentric bearing can be wired together to construct the border cells, and such allocentric spatial signals form the basis of hippocampal place cells (Barry et al., 2006; Hartley et al., 2000). Additionally, hippocampal place cells have been hypothesized to be representations of allocentric bearing and distance to specific external landmarks for facilitating vector-based navigation, where such “landmark-vector cells” or “object- vector cells” are constructed from egocentric bearing and allocentric head-direction cells (McNaughton et al., 1995). However, it remains unknown how sensory cortices can facilitate such egocentric-to-allocentric transformation. One possible theory of spatial coordinate transform in the dorsal visual system posits that gain modulation followed by slow short-term memory synaptic trace learning (Andersen et al., 1985; Andersen and Mountcastle, 1983; Duhamel et al., 1997; Rolls and Mills, 2019). Our new findings will provide more opportunities for development of new computational models along that direction.

## SUPPLEMENTAL INFORMATION

Supplemental information includes 1 table (Table S1) and 25 figures (Figures S1-S25).

## ACKNOWLEDGMENTS

We thank Jonathan Gould for reading of the manuscript. We thank Hao Wu, Wan-Neng Tang, Jia-Shun Ren, Cai-Ping Song, De-Guang Qi, Tong-Quan Liao, Hao Chen, Qian Chen, Yang Yang, Hui Yang and Sheng-Qing Lv for their encouragement and support. We thank Edvard Moser, May-Britt Moser, Neil Burgess and Caswell Barry for sharing their analytical codes with us. Special thanks to Ting-Ting Huang, Mi Zhang and A-Xiang Zhou for their technical assistance. S.-J.Z. was supported by the National Natural Science Foundation of China (Grant# 31872775). X.L. is supported by the Chongqing Municipality postdoctoral fellowship (Grant# cstc2019jcyj- bshX0035). Z.S.C. is partly supported by the US National Institutes of Health (R01-MH118928).

## AUTHOR CONTRIBUTIONS

Conceptualization: S.-J.Z.; Investigation: X.L., B.D. J.C. and S.-J.Z.; Formal Analysis: X.L. Resources: Z.S.C. and S.-J.Z.; Software: X.L.; Writing – Original Draft, Z.S.C., X.L. and S.-J.Z.; Writing – Review & Editing, Z.S.C., X.L. and S.-J.Z. Project Administration and Funding Acquisition: S.-J.Z.

## Materials and Methods

Materials and methods in this investigation were similar to those described previously (Long and Zhang, 2021; Zhang et al., 2013).

### Subjects

Twenty male Long-Evans adult rats (aged 2-4 months, weighting 250-450 grams on the day for chronic surgery) were implanted in these recording experiments. Rats were housed individually in transparent plexiglass cages (W × L × H: 35 cm × 45 cm × 45 cm) and kept on a reversed 12- hour light/12-hour dark schedule (lights on from 21:00 p.m. to 09:00 hours). All recording trials were performed during the dark phase. Rats were kept in a temperature-controlled (19-23°C) and humidity-adjusted (55-70%) vivarium. Rats were placed on a partially food-deprived schedule to about 85-90% of free-feeding body weight. Food restriction was imposed 8-24 hours before each training and recording trial. Water was supplied *ad libitum*. The animal experiments were approved by the National Animal Welfare Act under a protocol with the permission license number #SYXK-2017002 in accordance with the Animal Care and Use Committee from the Army Medical University and Xinqiao Hospital.

### Surgery and electrode placement

Tetrodes were constructed with four twisted 17 µm polyimide-coated platinum-iridium (90-10%) wires (#100167, California Fine Wire Company, USA). Tetrodes were plated with a 1.5% platinum solution to reduce electrode impedances to between 150 and 300 kΩ at 1 kHz through electroplating prior to surgery (nanoZ; White Matter LLC, Seattle, Washington, USA). Anesthesia was induced isoflurane mixed with oxygen (1.5-3.0% in O_2_), immobilized in a stereotaxic frame (David Kopf Instruments, Tujunga, California, USA) and kept on feedback-adjusted temperature control pad at 37°C. A self-assembled microdrive loaded with four tetrodes were implanted to target either the secondary visual cortex (V2, ∼2.5 mm lateral to the midline, ∼−4.5 mm anterior-posterior from bregma, ∼0.4-2.0 mm dorsal-ventral below the dura and at an angle of 10° from the medial-to-lateral direction in the coronal plane, ten rats), or the primary somatosensory cortex (S1, anterior-posterior: ∼0.2–2.2 mm posterior to bregma; medial-lateral: ∼2.2–3.4 mm lateral to midline; dorsal-ventral: ∼0.4–2.0 mm below the dura, 10 rats). Dental cement together with 8- 10 anchor screws were used to secure the microdive to the rat brain surface. Two screws were connected to the system ground for serving as the ground electrode.

### Behavioral protocol and data collection

Both unit activity and local field potentials (LFP) were recorded in parallel. The chronically implanted rats were rested at least one week of recovery before experimental recording, tetrode turning and data recording were initiated. Rats were trained to forage in different shapes of enclosure with a white cue card (297 × 210 mm2) mounted on one side of the interior wall. Food pellets were randomly dispersed into the enclosure intermittently to encourage free exploration. In the open filed, each recording trial lasted typically between 20 and 40 min to facilitate full coverage of the testing enclosure. Tetrodes were lowered very slowly in steps of 25 or 50 µm every day until well-separated single units can be identified. Data were acquired by an Axona system (Axona Ltd., St. Albans, U.K.) at 48 kHz, band-pass filtered between 0.8-6.7 kHz and a gain of 5,000-25,000 times. Spikes were digitized with 50 8-bit sample windows. Local field potentials were recorded from one of the electrodes with a low-pass filtered at 500 Hz.

### Spike sorting, cell-type identification and locational firing rate map

To identify well-isolated units, spike sorting was manually performed offline with graphical cluster-cutting software (TINT, version 4.4.16, Axona Ltd, St. Albans, U.K.), and the clustering was primarily based on features of the spike waveform (peak-to-trough amplitude and spike width), together with additional autocorrelations and cross-correlations separation tools and criteria. During the manual cluster cutting, units with similar or identical waveform shapes were strictly counted only once whenever similar or identical individual units were recorded and tracked across two consecutive recording trials. Only recording trials in which rats covered more than 80% of the running enclosure were taken for further analysis. To assess the quality of cluster separation, we calculated the well-established metric (L_ratio_) that measures the isolation distance between clusters (Schmitzer-Torbert et al., 2005).

A pair of small light-emitting diodes (LEDs) were attached to the head stage to track the rats’ speed, position and head orientation via an overhead video camera. Only spikes recorded during animal’s instantaneous running speeds > 2.5 cm/s were chosen for further analysis in order to exclude confounding behaviors such as immobility, grooming or rearing.

To classify firing fields and firing rate distributions, the position data were divided into 2.5 × 2.5 cm2 bins, and the path was smoothed with a 21-sample boxcar window filter (400 ms; 10 samples on each side). Units with > 100 spikes per session and with a coverage of >80% were included for further analyses. Maps for spike numbers and spike times were smoothed with a quasi- Gaussian kernel over the neighboring 5 × 5 bins. Spatial firing rates were calculated by dividing the smoothed map of spike numbers with spike times. The peak firing rate was defined as the highest rate in the corresponding bin within the spatial firing rate map. Mean firing rates were averaged from the whole session data.

### Identification of S1 or V2 border cells

The calculation of the border score was followed by previously publications (Boccara et al., 2010; Solstad et al., 2008; Zhang et al., 2013). Border cells were identified by calculating the border score, or the difference between the maximal length of any of the four walls touching on any single spatial firing field of the cell and the average distance of the firing field to the nearest wall, divided by the sum of those two values. The border score ranged from −1 for cells with perfect central firing fields to +1 for cells with firing fields that exactly line up with at least one entire wall. Firing fields were defined as summation of neighboring pixels with total firing rates higher than 0.3 times the unit’s maximum firing rate that covered a total area of at least 200 cm^2^.

Border cell classification was verified using population shuffling methods. For each permutation trial, the whole sequence of spike trains was time-shifted along the animal’s trajectory by a random period between 20 s and 20 s less than the length of the entire trial, with the end wrapped to the start of the trial. A spatial firing rate map was then obtained, followed by border score estimation.

The distribution of border score was calculated for the entire set of permutation trials from all recorded units, from which the threshold with the 99th percentile was determined. The unit was defined as a border cell if its border score was higher than the 99th percentile threshold derived from the shuffled data. Head direction tuning along four walls of border cells were measured in a 15-cm-wide area along each wall.

### Quantification of head-direction tuning

The calculation of mean vector length (mvl) was followed by previous publication (Boccara et al., 2010; Sargolini et al., 2006; Zhang et al., 2013). The rat’s head-direction was computed by the relative position of a pair of LEDs differentiated through their sizes. The directional tuning curve for each recorded unit was drawn by plotting the firing rate as a function of the rat’s head angle, which is divided into bins of 3° and then smoothed with a 14.5° mean window filter (2 bins on each side). The strength of directionality was measured by computing the mean vector length from circular distributed firing rates.

### Environmental manipulations

For visual landmark rotation, we first recorded S1 or V2 unit activity in the standard recording session followed by a 45° cue-card rotation in the clockwise or counterclockwise orientation. Next, another standard session was performed with the cue-card rotated back to the original position. For recording in the elevated platform without walls, we first recorded S1 or V2 units in the square box, followed by the recording in the elevated platform without walls. For recording in the dark, we first recorded S1 or V2 units in the square box under the light condition, followed by the recording in the dark and back to the light condition. For recording in different geometric shapes, we first recorded S1 or V2 units in the square box, followed by the recording in the circular enclosure and back to the square box. For recording in the rectangle arena, a 1-m long and 50-cm high plate was inserted into the center of the 1-m square box either in parallel or orthogonal to the original cue card, stretching the 1 × 1 m^2^ square into a 0.7 × 1 m^2^ or 1 × 0.7 m^2^ rectangle. For the recording of border cells in the presence of an inserted wall, a baseline recording trial was performed followed by the additional introduction of one discrete wall in three different orientations: from horizontal to vertical to diagonal, then ended with another baseline recording session. For recording in a larger environment with two geometric borders, a small 50 × 50 × 30 cm^3^ (L × W × H) square box was placed in the center of a larger 150 × 150 cm2 arena, creating an outer ring and an inner ring two separate geometric borders. For recording in running boxes with three geometric borders, a small 80 × 80 × 5 cm3 ring (inner wall size: 70 × 70 × 5 cm^3^) was placed in the center of a large 150 × 150 cm2 box, creating an outer ring, a middle ring and an inner ring of geometric borders.

### Histology and tetrode track location

In the end of the final recording session, implanted animals were deeply anaesthetized with overdose of sodium pentobarbital (0.01ml/g) and perfused intracardially with ice-cold 1 x phosphate-buffered saline (PBS) followed by 4% ice-cold paraformaldehyde (PFA) in 1 x PBS. Brains were taken out and post-fixed in 4% PFA in 1 x PBS at 4°C for more than 24 hours. The brain was then transferred into 10, 20 and 30% sucrose/PFA solution sequentially across 72 hours before sectioning by using a cyrostat. Thirty-micron-thick coronal sections were serially cut and obtained through the implanted brain area. Sections were mounted on glass slides and stained with Cresyl Violet (Sigma-Aldrich). The final recording positions were imaged and determined from digitized images of the Nissl-stained sections scanned on the Olympus Slideview VS200 Digital Slide Scanner. Positions of each individual recordings were estimated from the deepest tetrode track according to the daily notebook on tetrode advancement. The tissue shrinkage correction was calculated by dividing the distance between the brain surface and electrode tips by the final advanced depth of the recording electrodes. Electrode traces were confirmed to be located either S1 or V2 from twenty implanted rats based on the reference figures published in the sixth edition of *The Rat Brain in Stereotaxic Coordinates* (Paxinos and Watson, 2007).

### Data availability

Recording dataset will become available in a forthcoming public domain, and reasonable request into acquiring the raw data beforehand should be directed to the corresponding author.

### Code availability

The source codes used in this study are available from the corresponding author upon reasonable request.

## Supplementary Table S1 and Supplementary Figures S1-S25

**Table S1.** Summary of Border Cells from S1 and V2.

| Parameters | S1 | V2 |
| --- | --- | --- |
| Number of Border Cells | 148 | 161 |
| Mean Firing Rate (Hz) | 2.20 ± 0.09 | 1.88 ± 0.08 |
| Border Score | 0.69 ± 0.08 | 0.68 ± 0.08 |
| CW-Border | 40 | 44 |
| CCW-Border | 28 | 45 |
| Regular Border | 72 | 80 |
| CW: R/L Hemisphere | 10/30 | 6/38 |
| CCW: R/L Hemisphere | 21/7 | 25/20 |
| CW: Mean MVL(Wall1-4) | 0.49/0.58/0.56/0.53 | 0.49/0.51/0.47/0.50 |
| CCW: Mean MVL(Wall1-4) | 0.53/0.52/0.52/0.51 | 0.69/0.68/0.66/0.67 |

**Figure S1.**
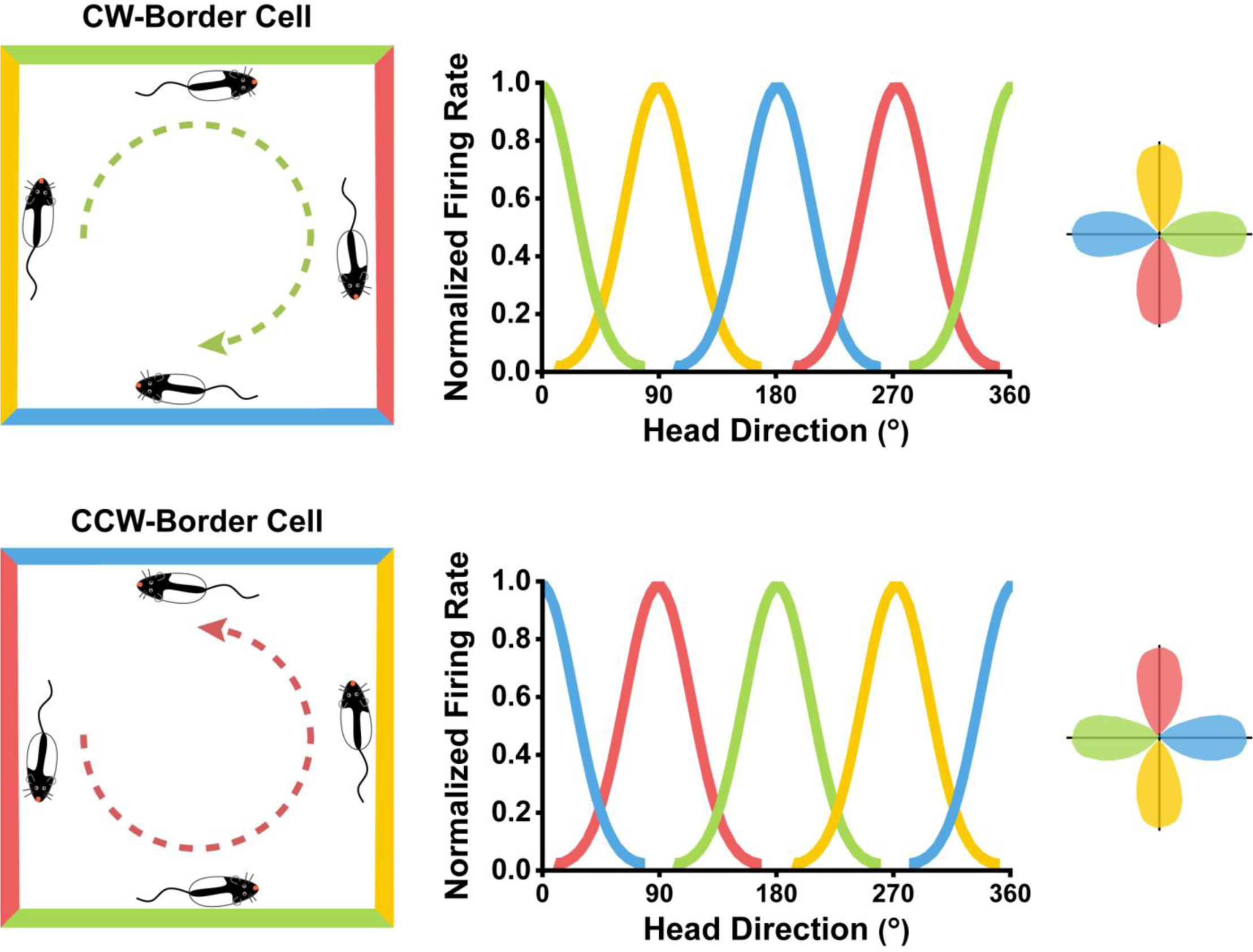
Border Cells Fire in a Clockwise or Counterclockwise Manner. The left panel shows an illustration of the specific heading direction in which CW-border cells (upper panel) and CCW-border cells (bottom panel) increased their firing rates. CW- and CCW-border cells were active only when animals moved unidirectionally in a clockwise and counterclockwise manner, respectively. The middle panel shows the directional tuning curve which resulted from each color-coded wall plotted in the left panel. The peak directions of head-direction tuning curves increased in a 90°-step manner for both CW- and CCW-border cells. However, the corresponding peaks of the directional tuning curves for the same color-coded wall differed 180° for CW- and CCW-border cells. The polar plot in the rightmost column shows the distribution of peak directions of head-direction tuning along the four color- coded walls.

**Figure S2.**
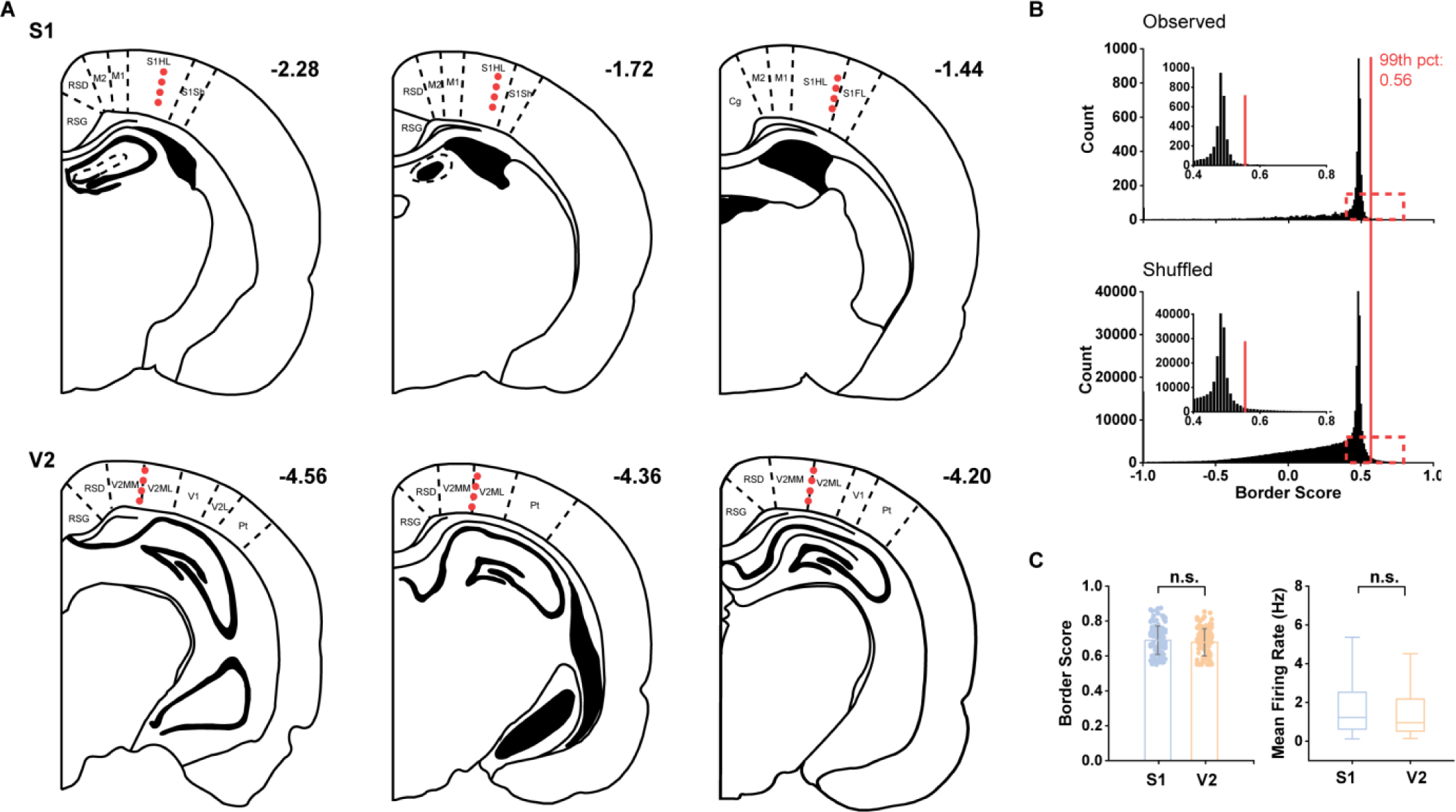
Identification of Border Cells. **(A)** Anatomical location of the electrode tracks in V2 and S1. S1HL: primary somatosensory cortex, hindlimb region; S1FL: primary somatosensory cortex, forelimb region; S1Sh: primary somatosensory cortex, shoulder region; Cg: cingulate cortex; RSG: retrosplenial granular cortex; RSD: retrosplenial dysgranular cortex; M2: secondary motor cortex; M1: primary motor cortex; V2MM: secondary visual cortex, mediomedial area; V2ML: secondary visual cortex, mediolateral area; Pt: parietal cortex, posterior area; V1: primary visual cortex; S1: primary somatosensory cortex. **(B)** Distribution of border score for observed data (top panel) and shuffled data (bottom panel). Red line indicates the 99^th^ percentile for the border score derived from the shuffled data. The inset shows the zoomed-in of the red dashed box. **(C)** Border scores and mean firing rates from the two border populations of S1 and V2 show no significant difference.

**Figure S3.**
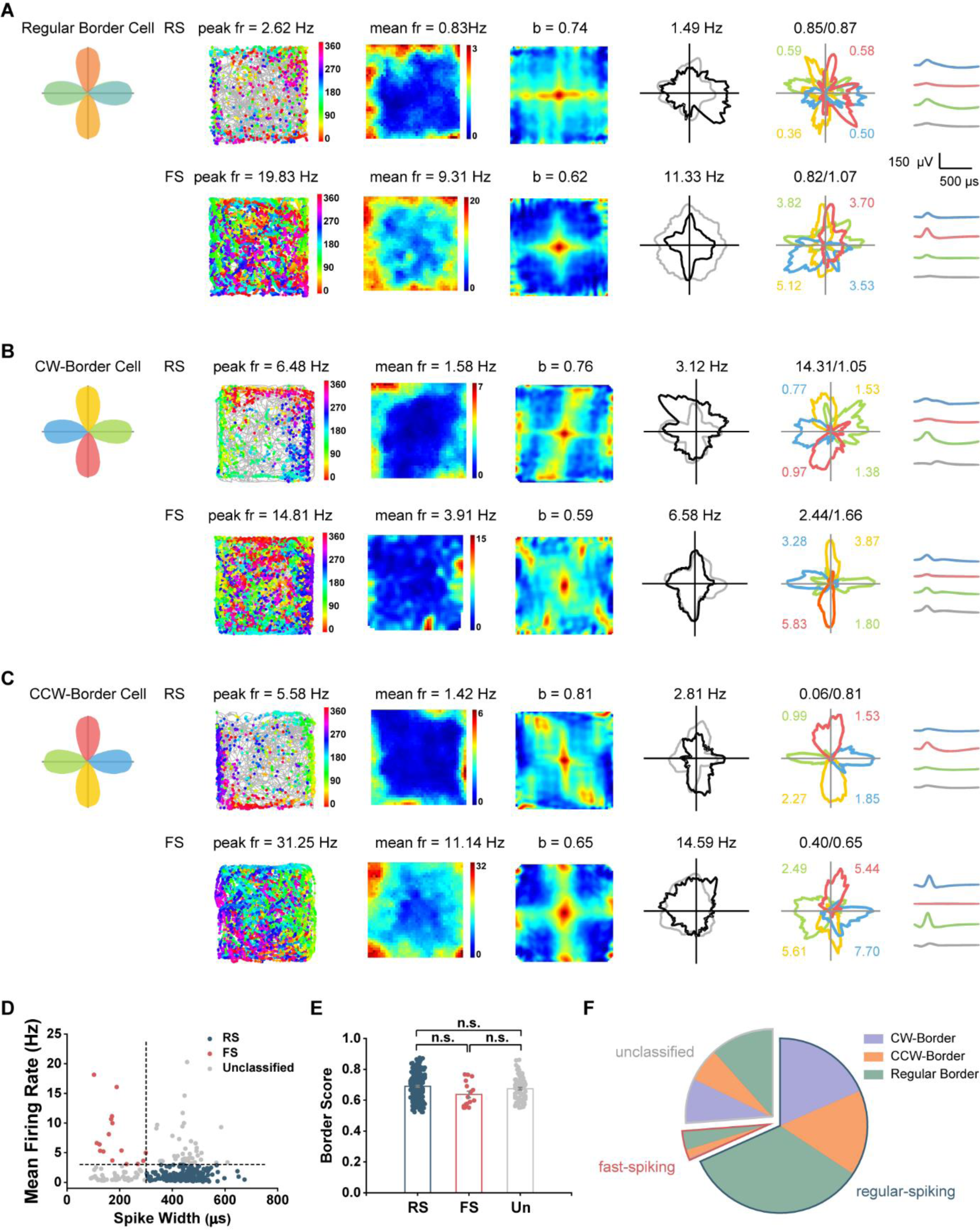
Classification of Subtypes of Border Cells in the Rat S1 and V2. (**A-C**) Representative subtype examples of border cells in the rat S1 and V2. Schematic diagrams showing a representative regular-spiking neuron and a representative fast-spiking neuron with regular border (**A**), CW-border (**B**) and CCW-border (**C**) firing properties, respectively. Notations and symbols are similar to **Figure 1**. **(D)** The distribution of spike width (peak to trough) versus mean firing rate for all identified border cells. RS, regular-spiking; FS, fast-spiking. **(E)** The border score of three subtypes of border cells. **(F)** The composition of regular, CW- and CCW-border cells in each neuronal subtypes.

**Figure S4.**
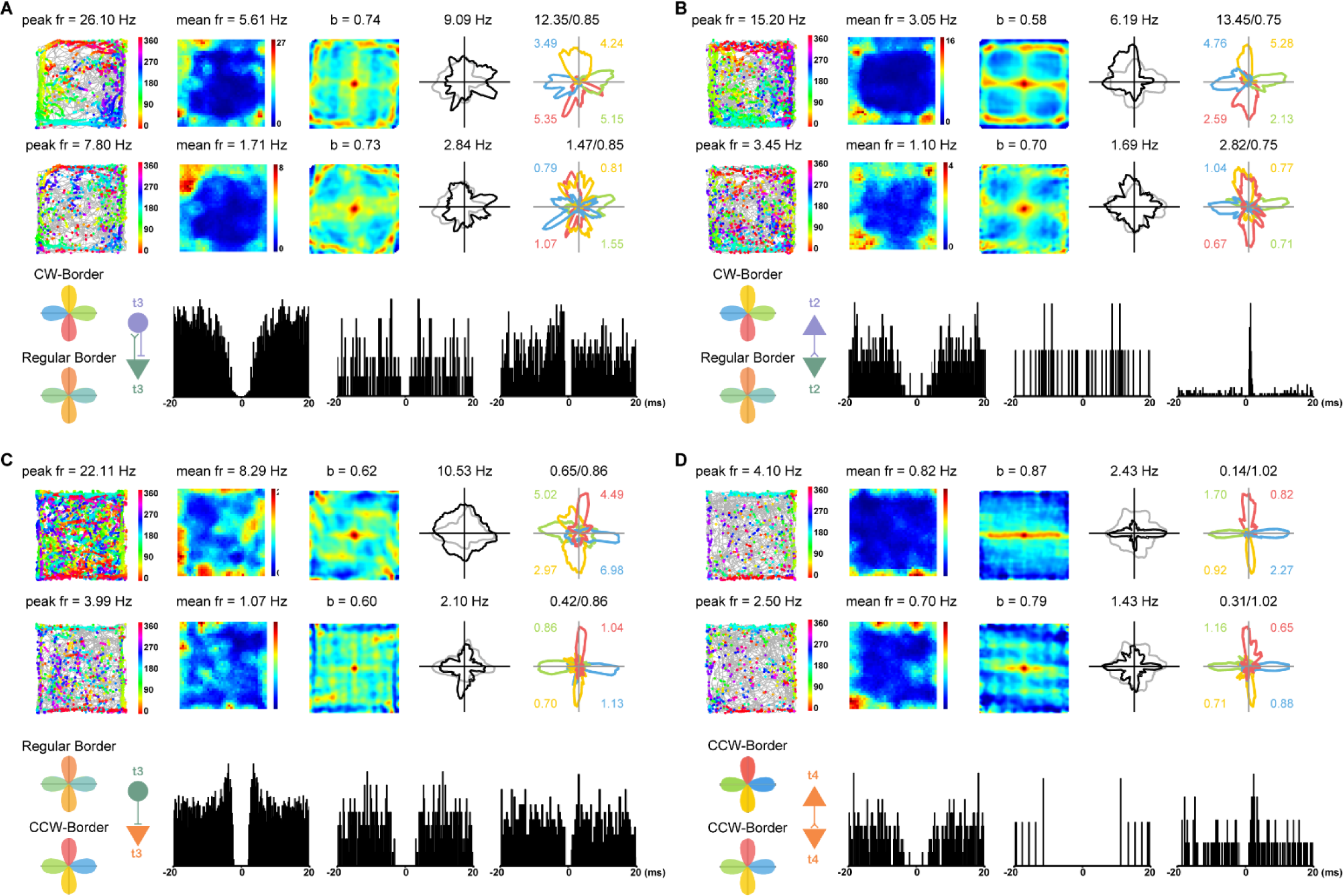
Putative Monosynaptic Connections Between Simultaneously Recorded CW-, CCW- or Regular Border Cells. (**A**, **B**) Spatial responses for representative CW-border cells (upper panels) and co-recorded regular border cells (middle panels). Notations and symbols are similar to **Figure 1**. Putative monosynaptic connections as revealed by the spike-time cross-correlogram (middle panels). Left, hypothesized synaptic connectivity with CW-border cells (purple triangle), regular border cells (green triangle), and putative interneurons (circle); Left middle, autocorrelogram of the reference cell; Right middle, autocorrelogram of the target cell; Right, cross-correlogram. The tetrode number (t1-t4) on which the representative cell was recorded is labeled. (**C**, **D**) Same as (**A**, **B**) except for simultaneously recorded representative CCW-border cells (red) and regular border cells (blue).

**Figure S5.**
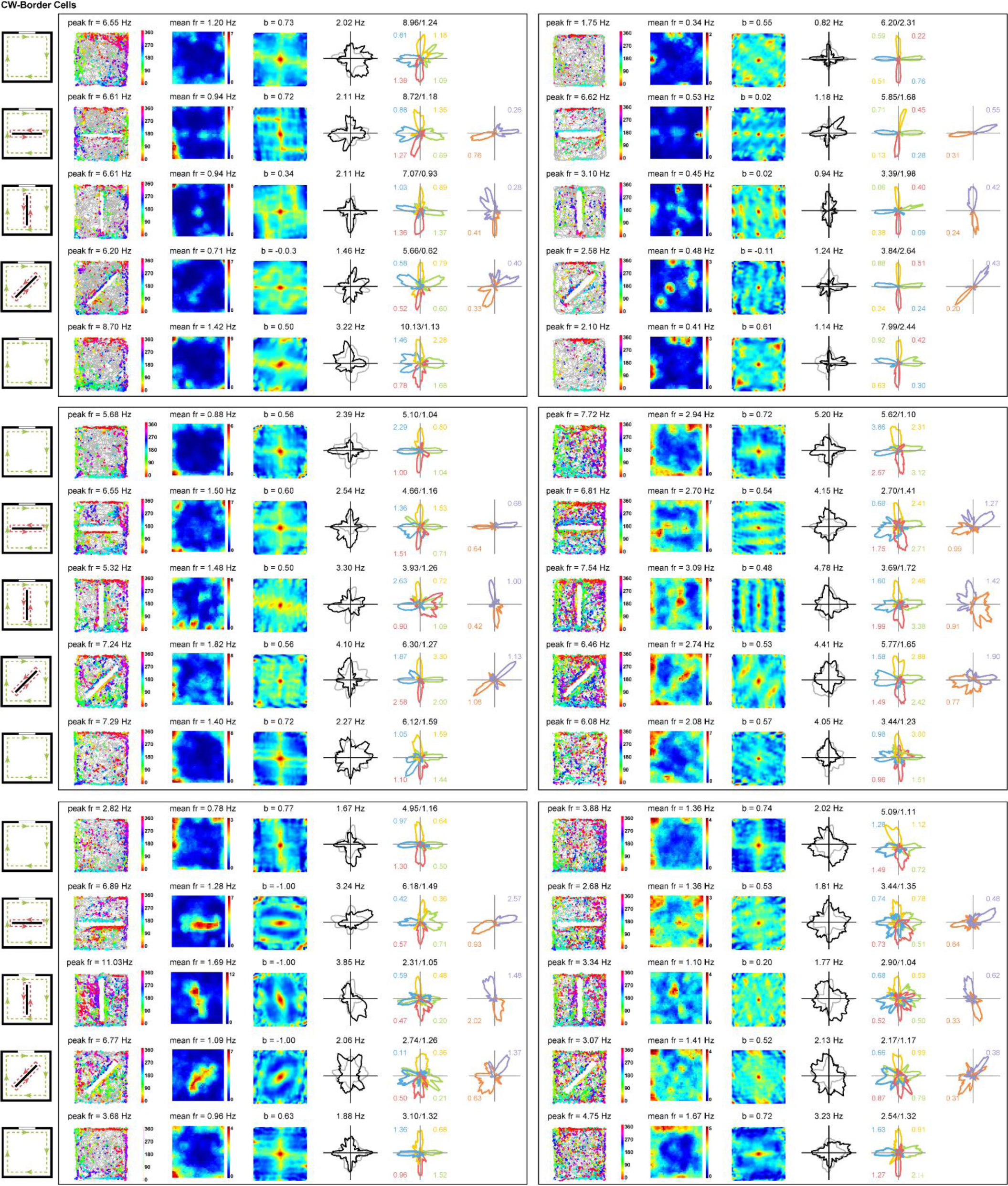
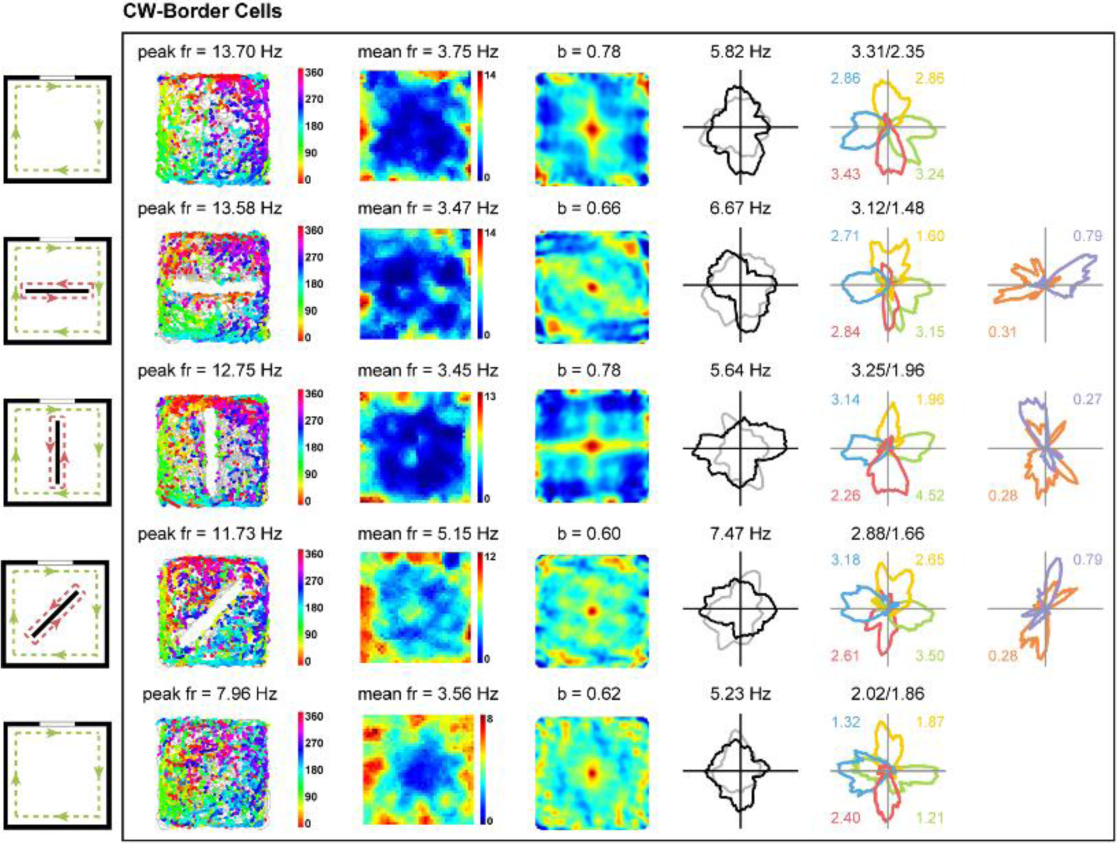
Spatial Firing Patterns for the Entire Sample of CW-Border Cells before and after New Wall Insert. Notations and symbols are similar to **Figure 1**.

**Figure S6.**
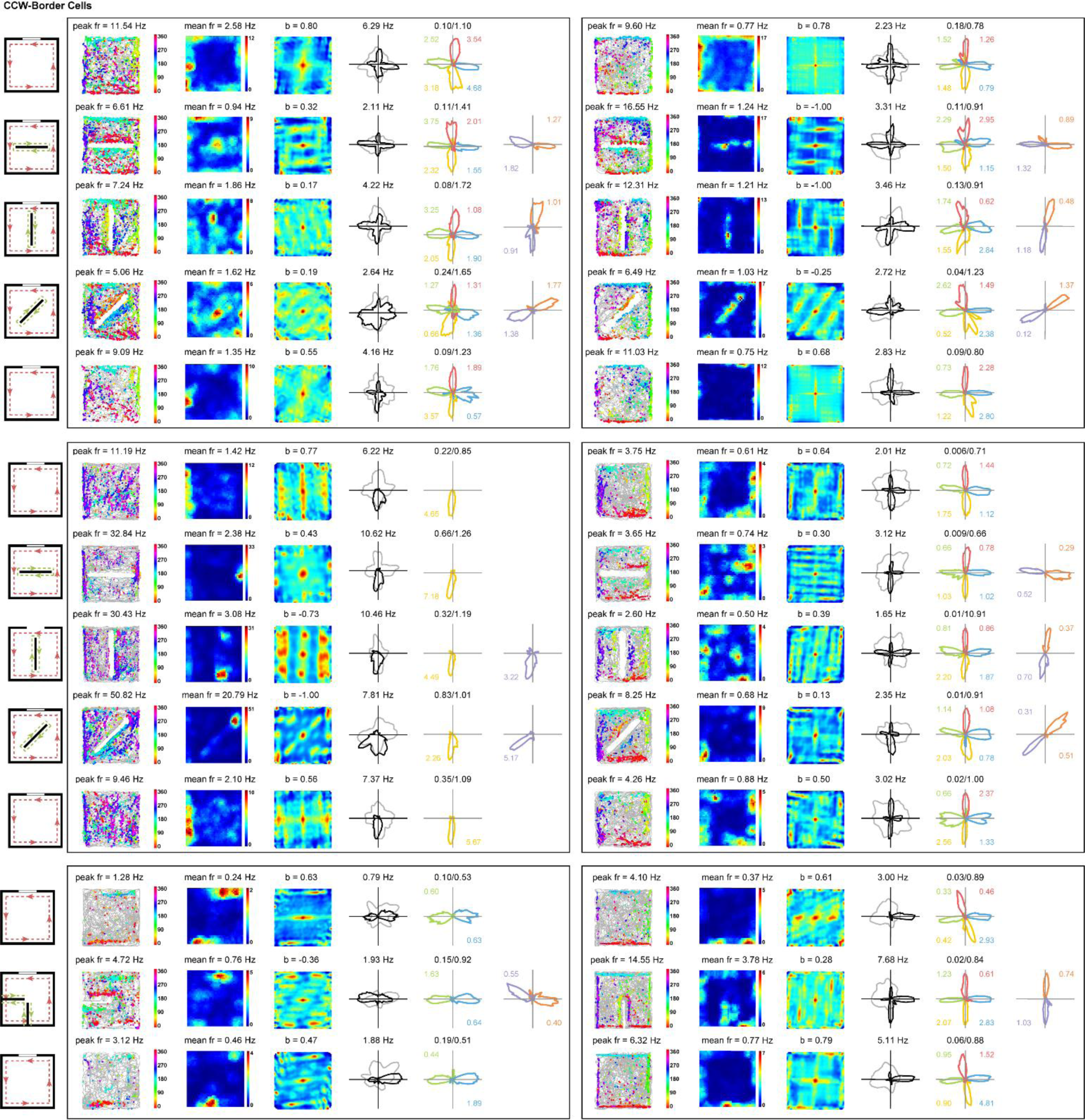
Spatial Firing Patterns for the Entire Sample of CCW-Border Cells before and after New Wall Insert. Notations and symbols are similar to **Figure 1**.

**Figure S7.**
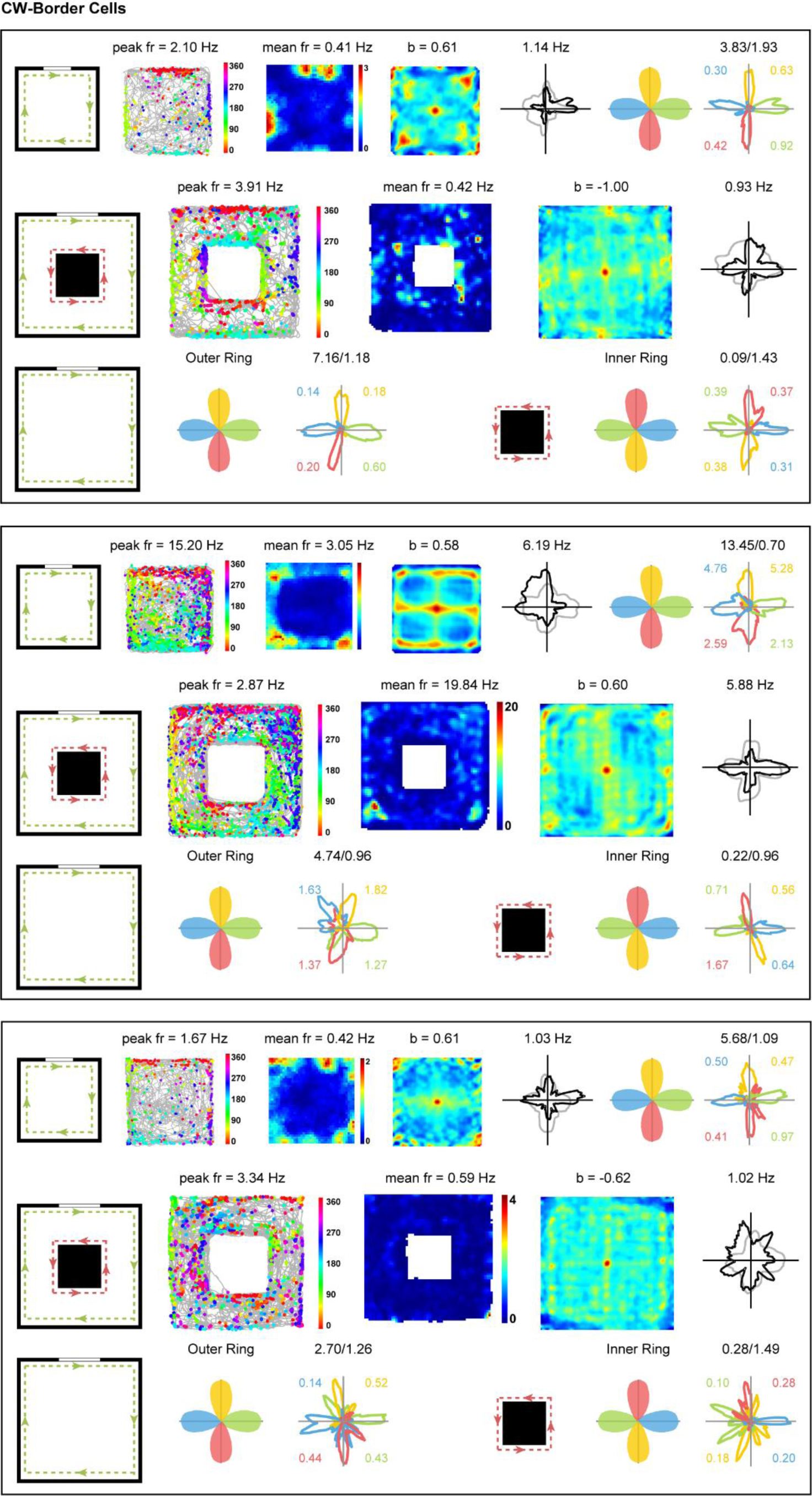
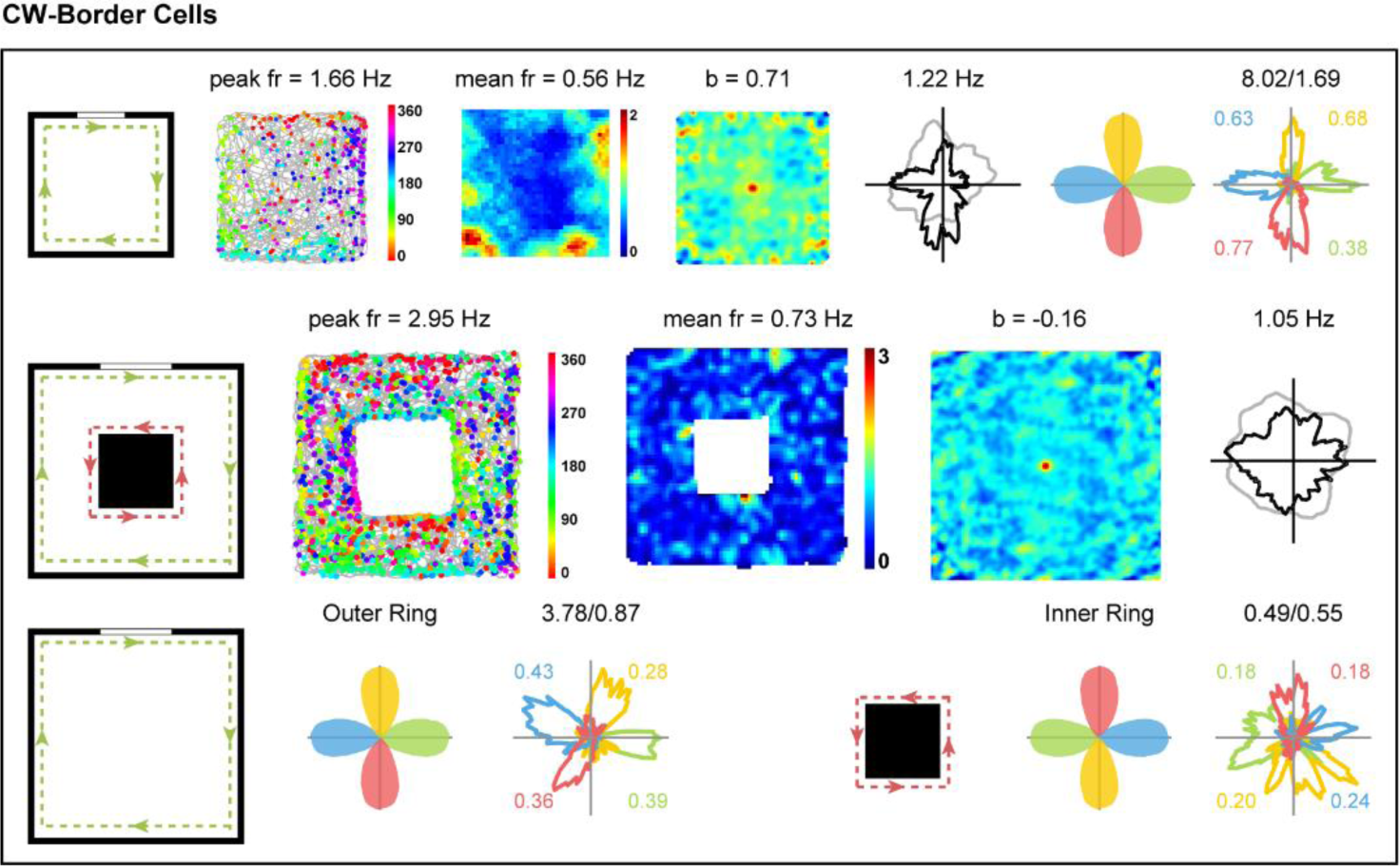
Spatial Firing Patterns for the Entire Sample of CW-Border Cells in Environments with Two Geometric Borders. Notations and symbols are similar to **Figure 1**.

**Figure S8.**
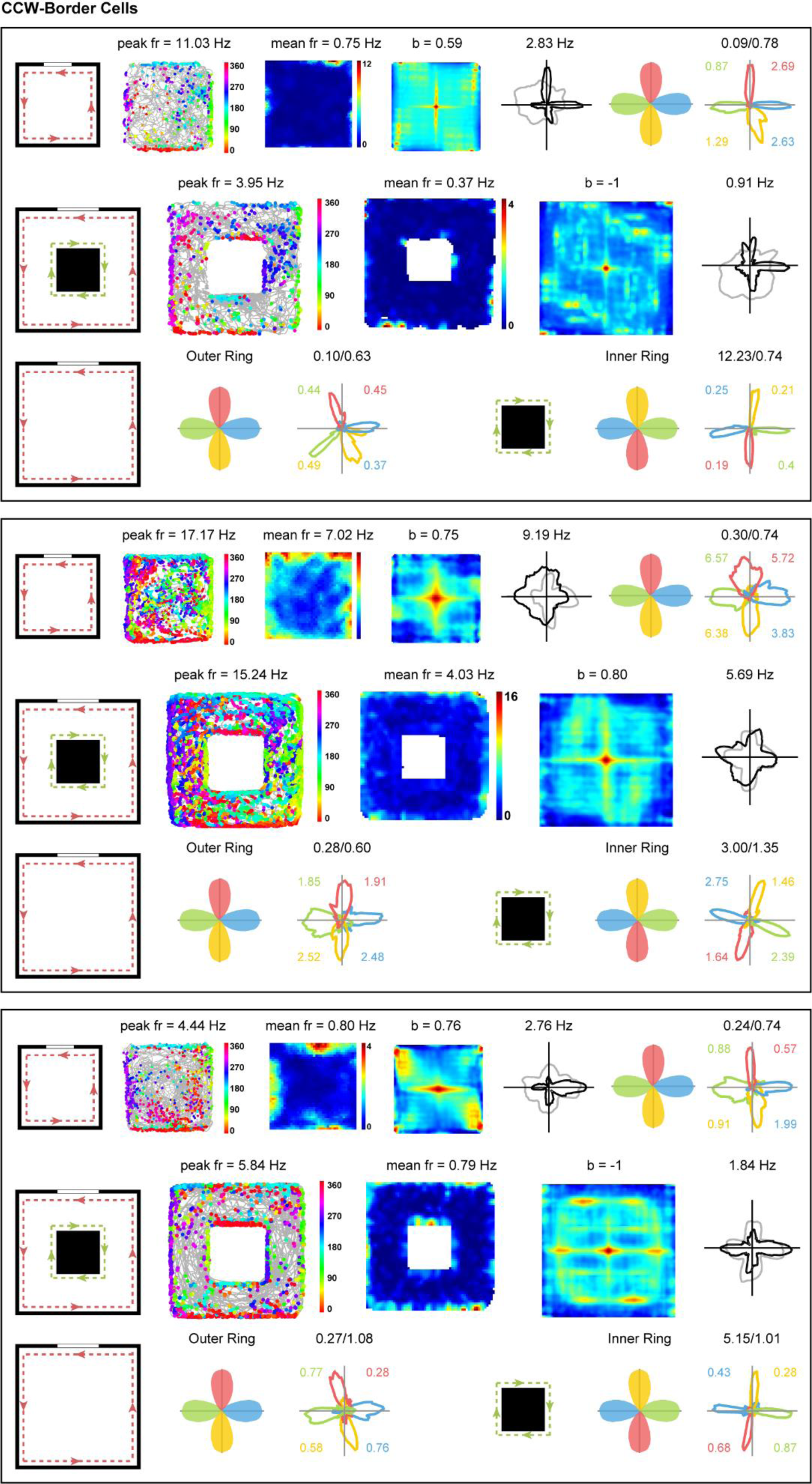
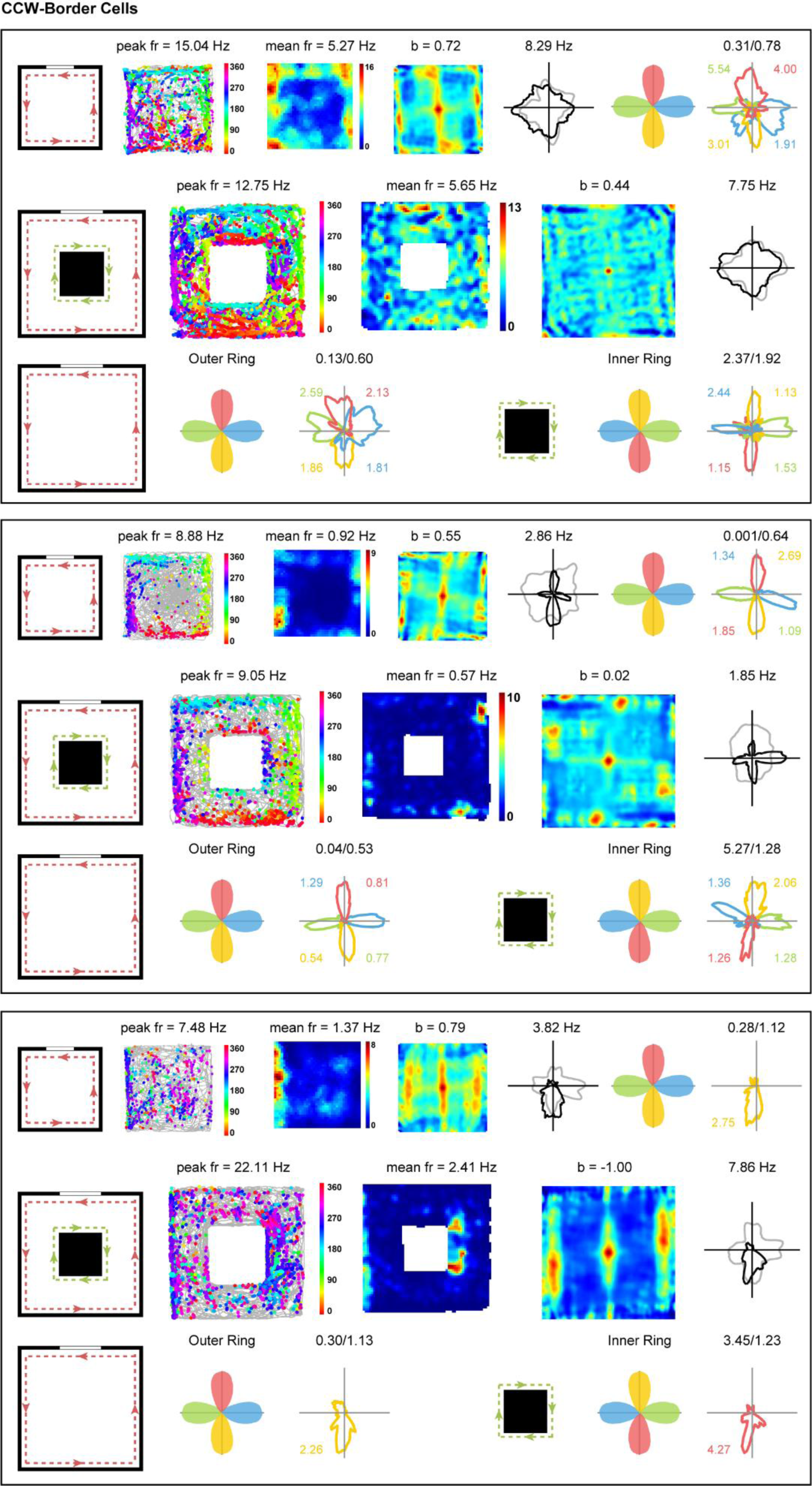
Spatial Firing Patterns for the Entire Sample of CCW-Border Cells in Environments with Two Geometric Borders. Notations and symbols are similar to **Figure 1**.

**Figure S9.**
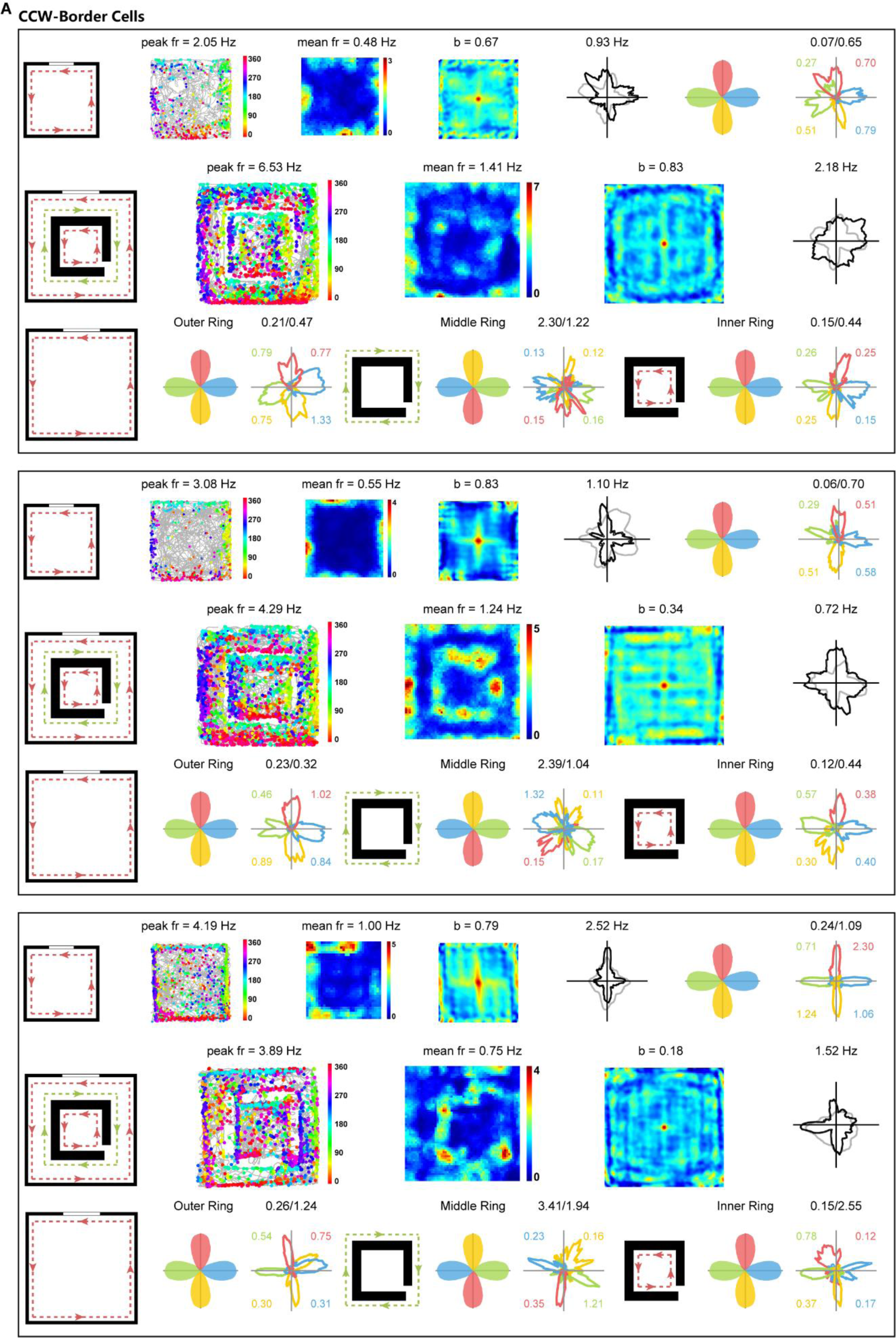
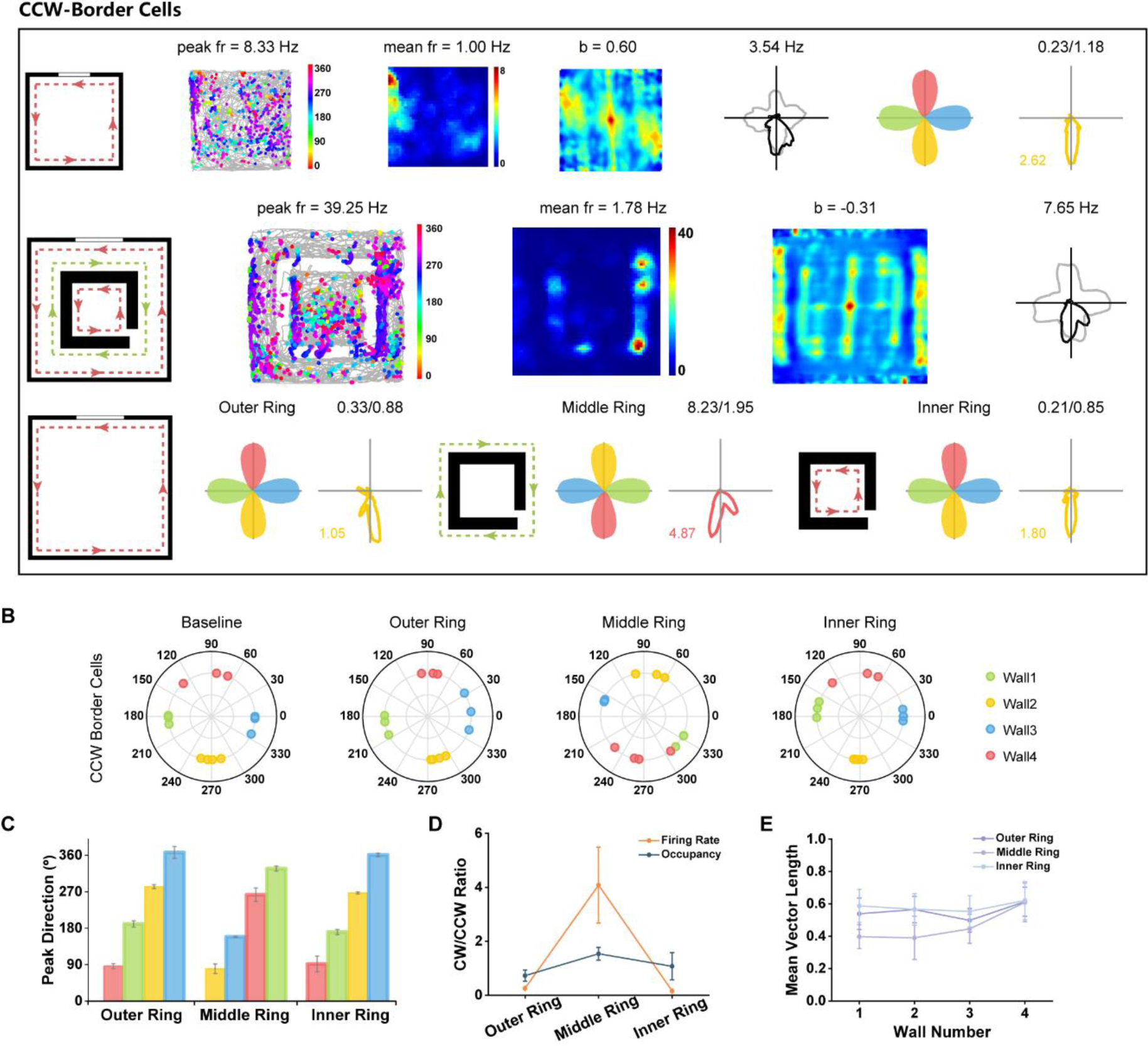
Spatial Responses of CCW-Border Cells in Environments with Three Geometric Borders. **(A)** Spatial responses of representative cortical CCW-border cells in running boxes with three geometric borders. A small 80 × 80 × 5 cm^3^ ring (inner wall size: 70 × 70 × 5 cm^3^) was placed in the center of a large 150 × 150 cm^2^ box which created an outer ring, a middle ring and an inner ring of geometric borders and the rat could climb over the ring. Notations and symbols are similar to **Figure 1**. **(B)** Distribution of peak directions of head direction tuning for four color-coded walls in baseline and boxes with three-ring conditions. Noted that the head directions of the four sides of the middle ring showed opposite tuning patterns to those of outer ring and inner ring. **(C)** Summary of the peaks of directional tuning of the outer ring, middle ring and inner ring for CCW- border cells. **(D)** CW/CCW ratio in firing rate (red) and occupancy (blue) for outer ring, middle ring and inner ring. Note that outer ring and inner ring shared similar CW/CCW ratio in firing rate (<0.5), while middle ring had opposite CW/CCW ratio in firing rate similar to that of CW-border cells (>2). **(E)** Mean vector length of directional responses for four sides of the outer ring, middle ring and inner ring.

**Figure S10.**
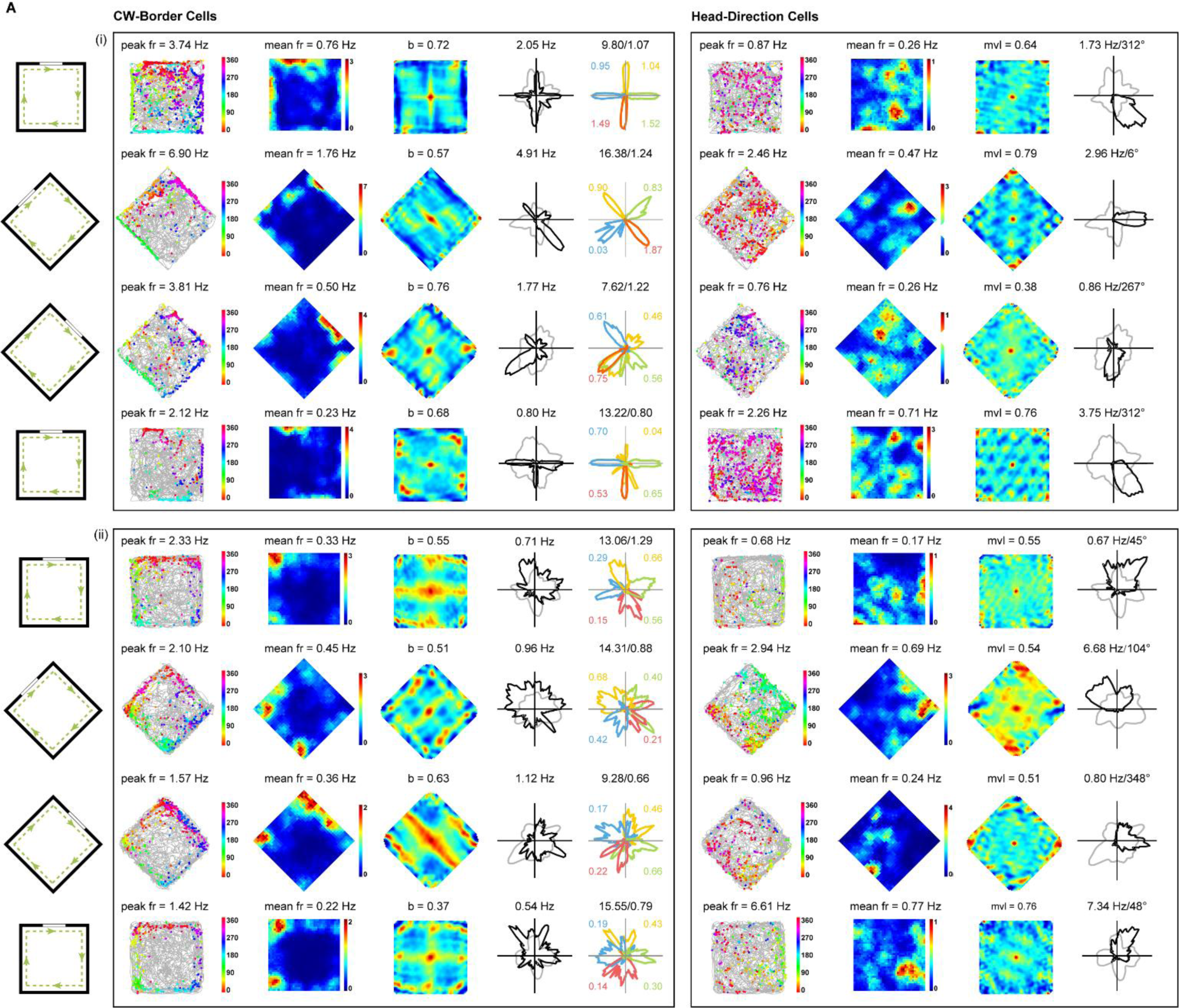
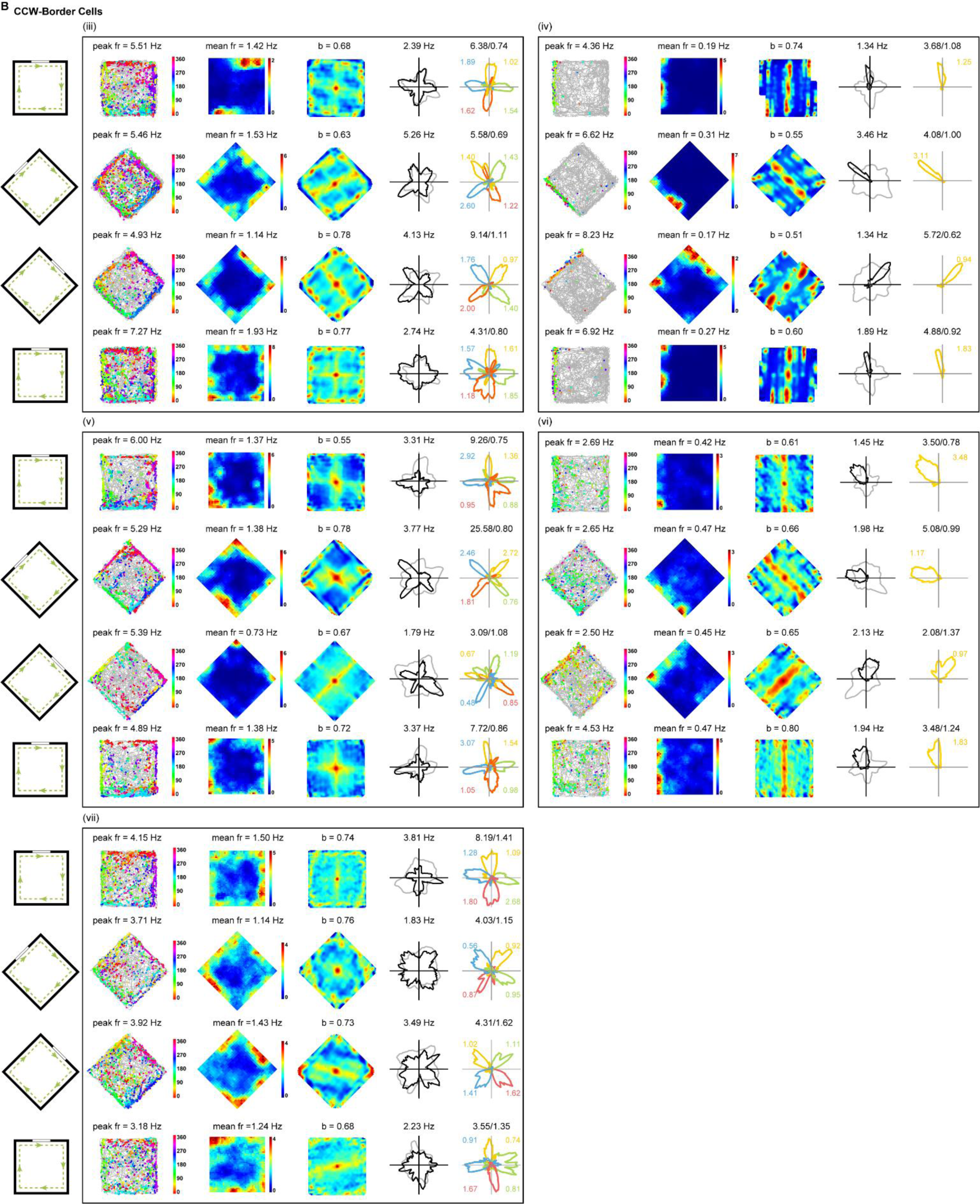
Spatial Firing Patterns for the Entire Sample of CW-Border Cells during Cue Manipulation. (**A**, **B**) Spatial responses for the entire sample of CW-border cells (**A**, **B**) with spatial responses of simultaneously recorded head-direction cells (**A**) during 45° rotation of cue manipulation. Notations and symbols are similar to **Figure 1**. For the simultaneously recorded head-direction cell, only the first four column are shown [(i) in (**A**) and (i)-(v) in (**B**)].

**Figure S11.**
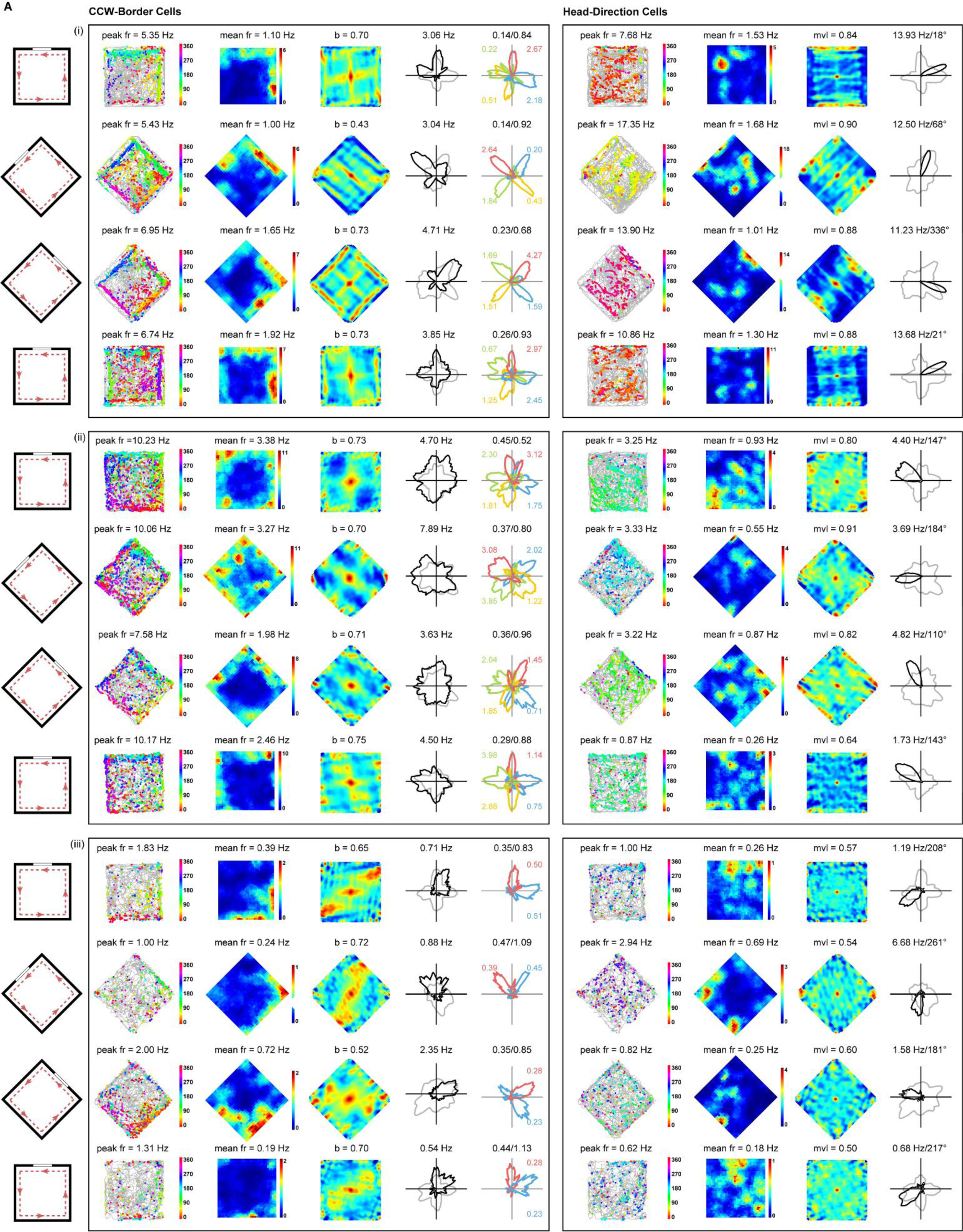
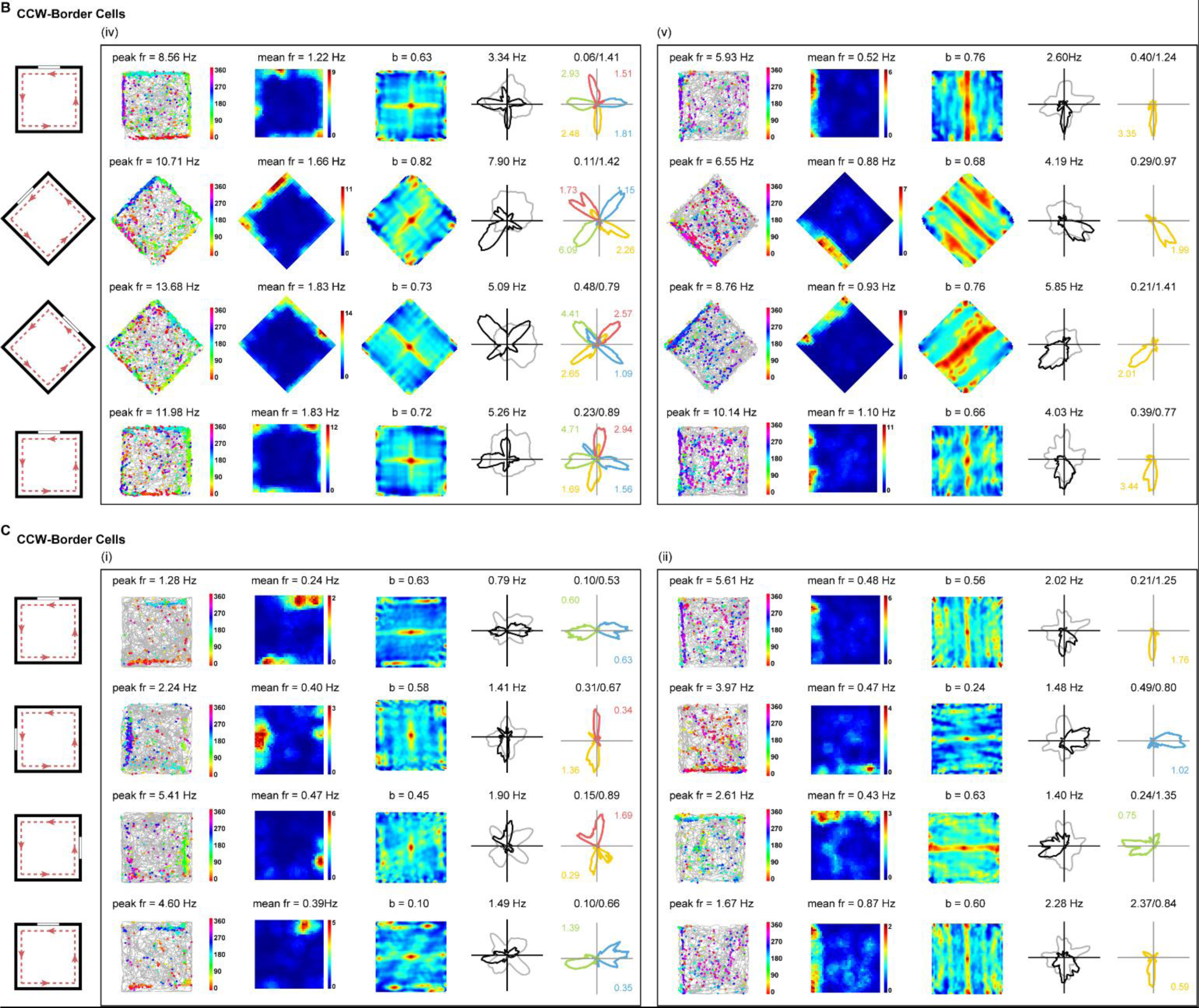
Spatial Firing Patterns for the Entire Sample of CCW- and CW-Border Cells during Cue Manipulation. (**A**, **B**) Spatial responses for the entire sample of CCW-border cells (**A**, **B**) with spatial responses of simultaneously recorded head-direction cells (**A**) during 45° rotation of cue manipulation. Notations and symbols are similar to Figure 1. For the simultaneously recorded head-direction cell, only the first four column are shown [(i) in (**A**) and (i)-(v) in (**B**)]. (**C**) Spatial responses of CCW-border cells firing along two walls (i) and a single wall (ii) during 90° rotation of cue manipulation.

**Figure S12.**
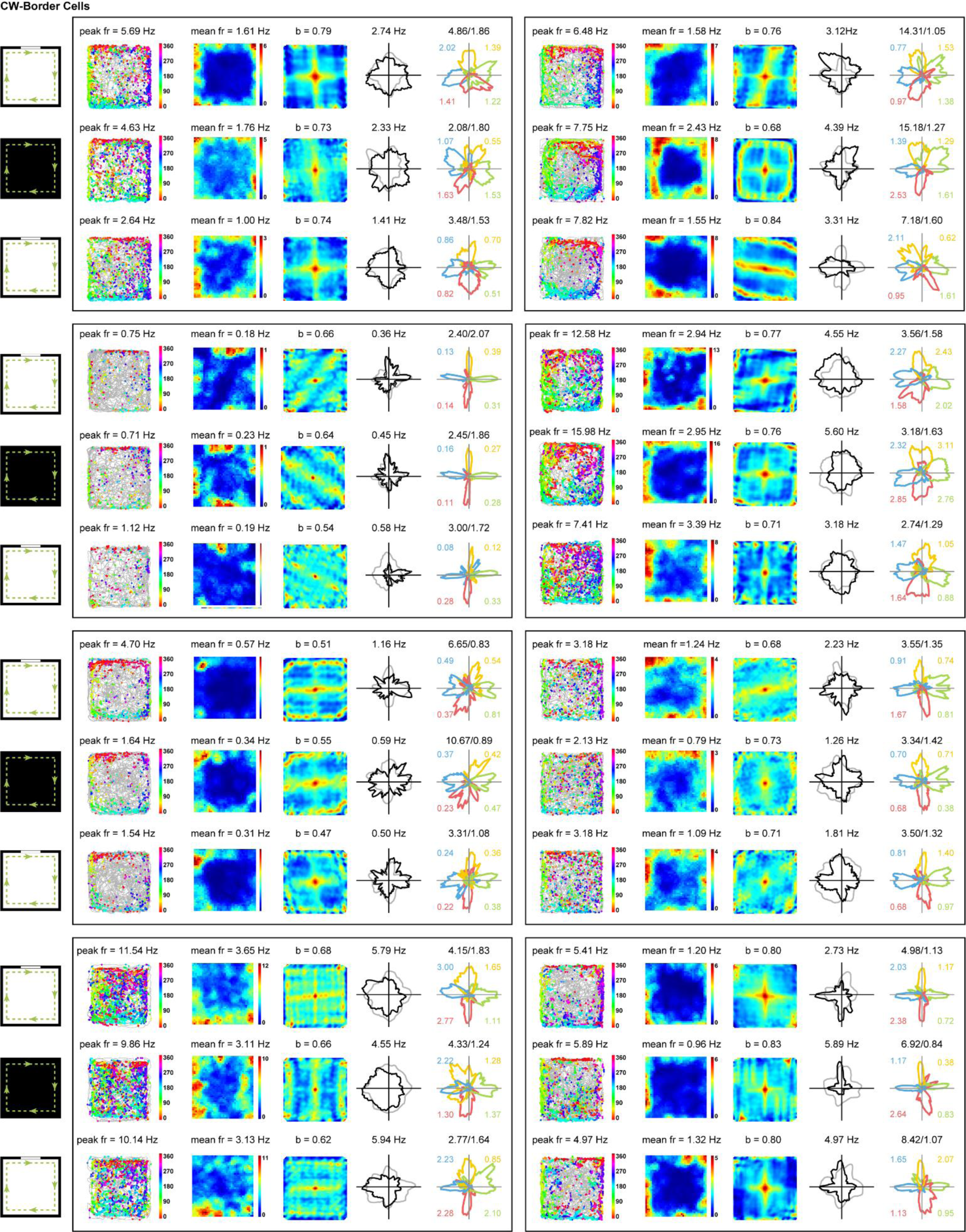
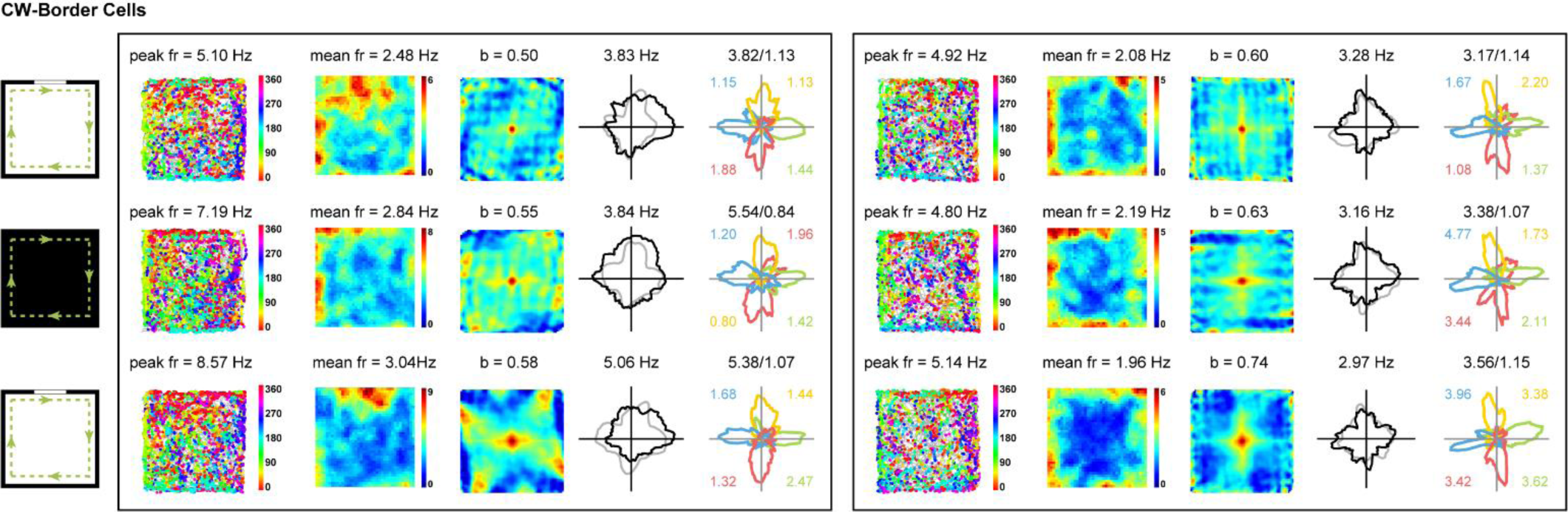
Spatial Firing Patterns for the Entire Sample of CW-Border Cells in the Darkness and Light conditions. Spatial responses for the entire sample of CW-border cells in the light (L), darkness (D) and light (L’) conditions. Notations and symbols are similar to **Figure 1**.

**Figure S13.**
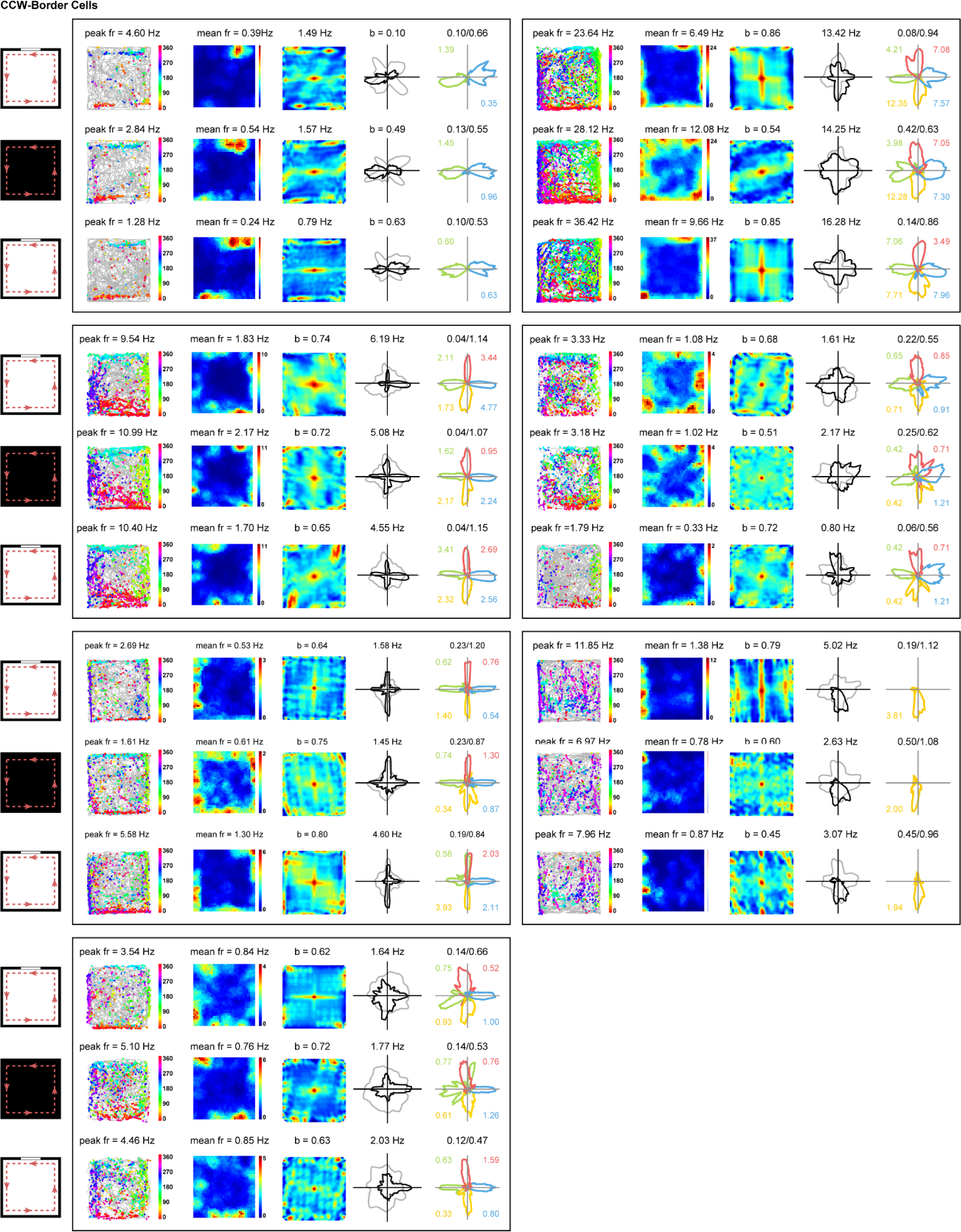
Spatial Firing Patterns for the Entire Sample of CCW-Border Cells in the Darkness and Light conditions. Spatial responses for the entire sample of CCW-border cells in the light (L), darkness (D) and light (L’) conditions. Notations and symbols are similar to **Figure 1**.

**Figure S14.**
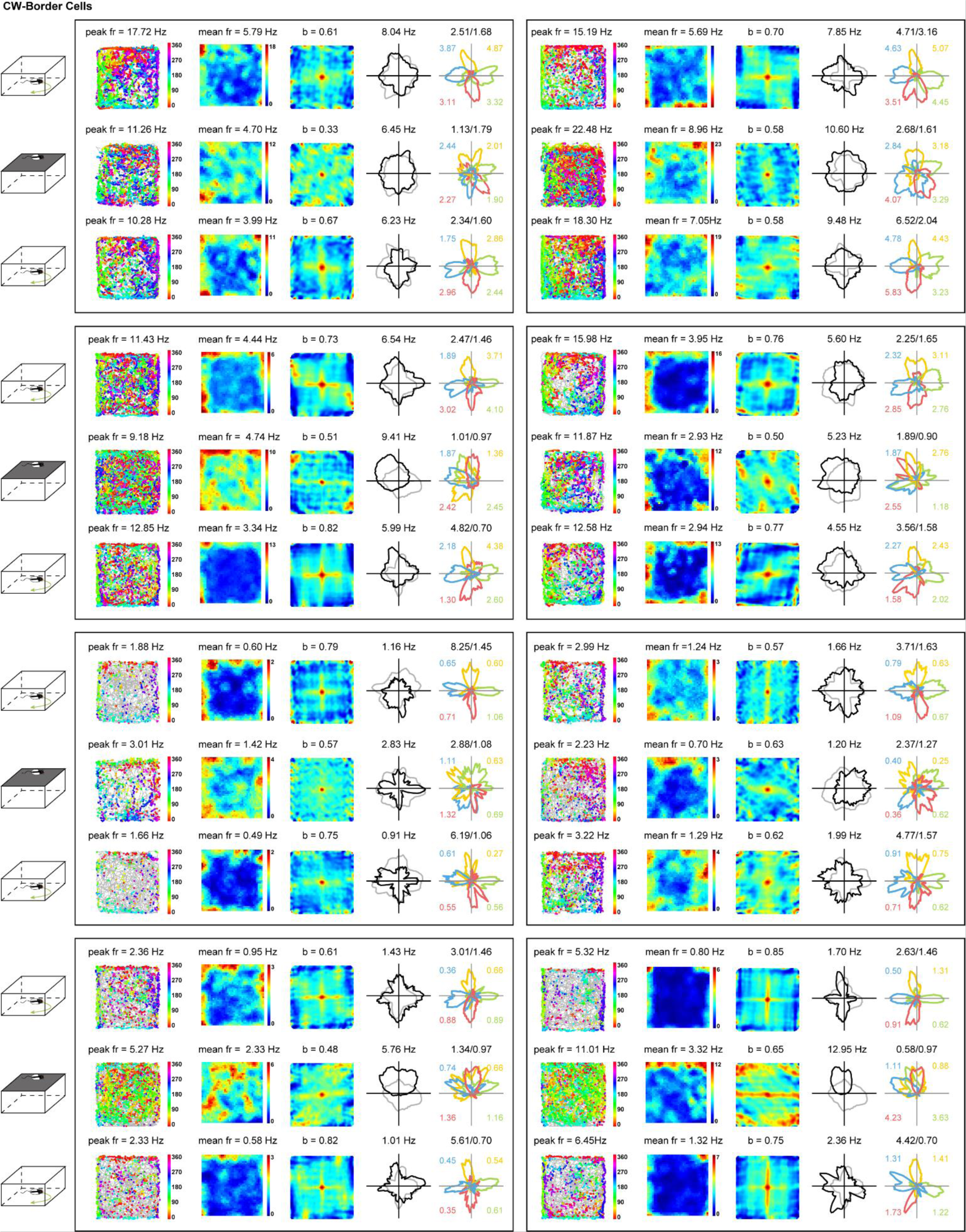
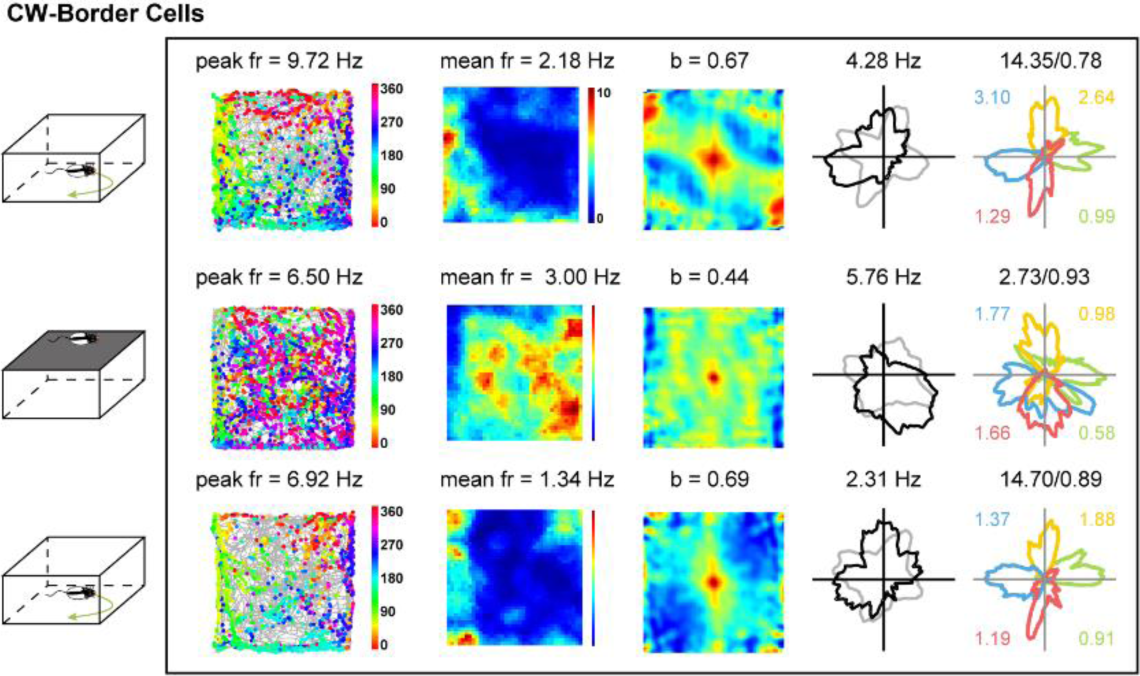
Spatial Firing Patterns for the Entire Sample of CW-Border Cells in Elevated Wall-less Platform. Spatial responses for the entire sample of CW-border cells, respectively, in the baseline (B), elevated wall-less platform (E) and baseline (B’) conditions. Notations and symbols are similar to **Figure 1**.

**Figure S15.**
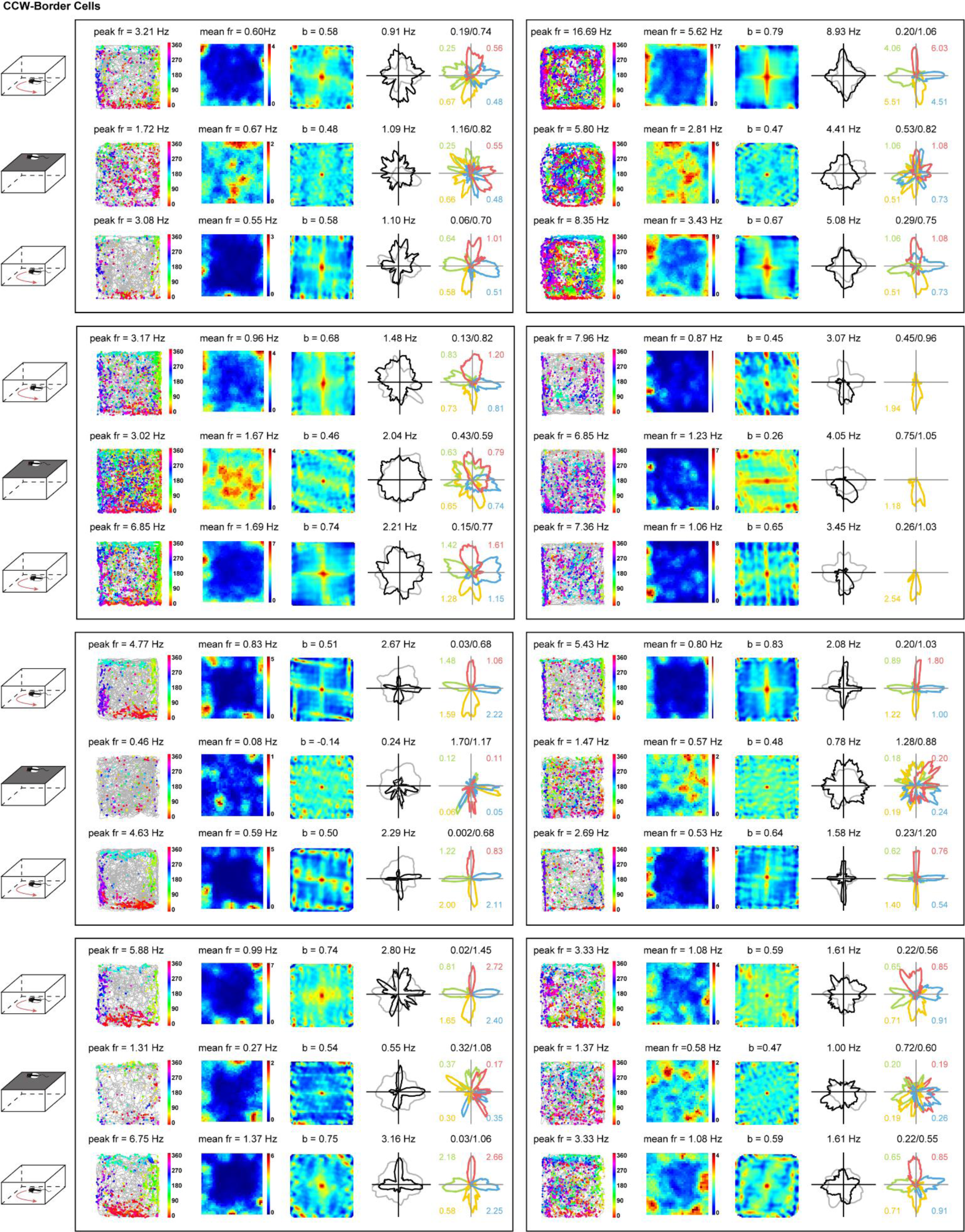
Spatial Firing Patterns for the Entire Sample of CCW-Border Cells in Elevated Platform without Physical Walls. Spatial responses for the entire sample of CCW-border cells, respectively, in the baseline (B), elevated wall-less platform and baseline (B’) conditions. Notations and symbols are similar to **Figure 1**.

**Figure S16.**
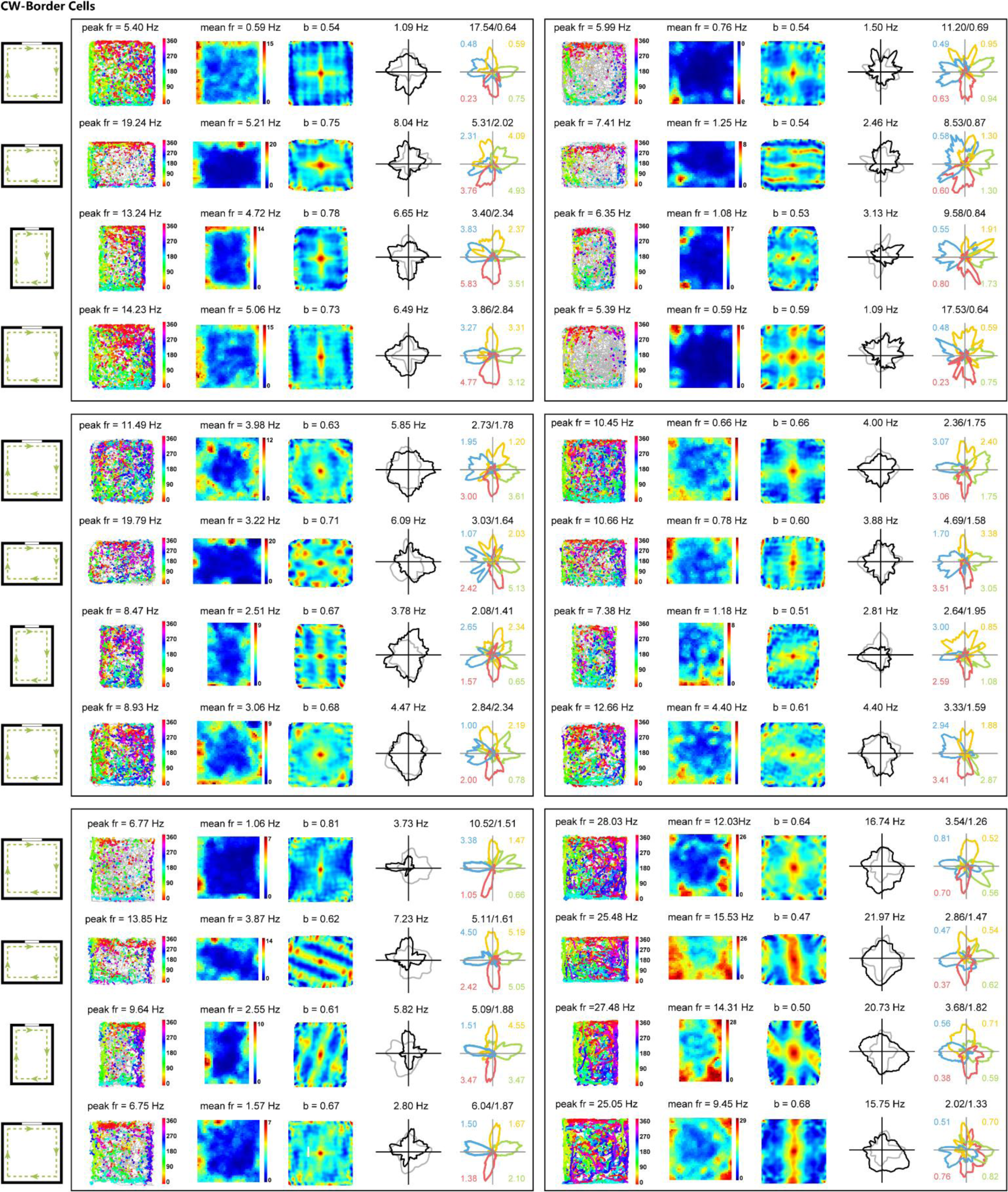
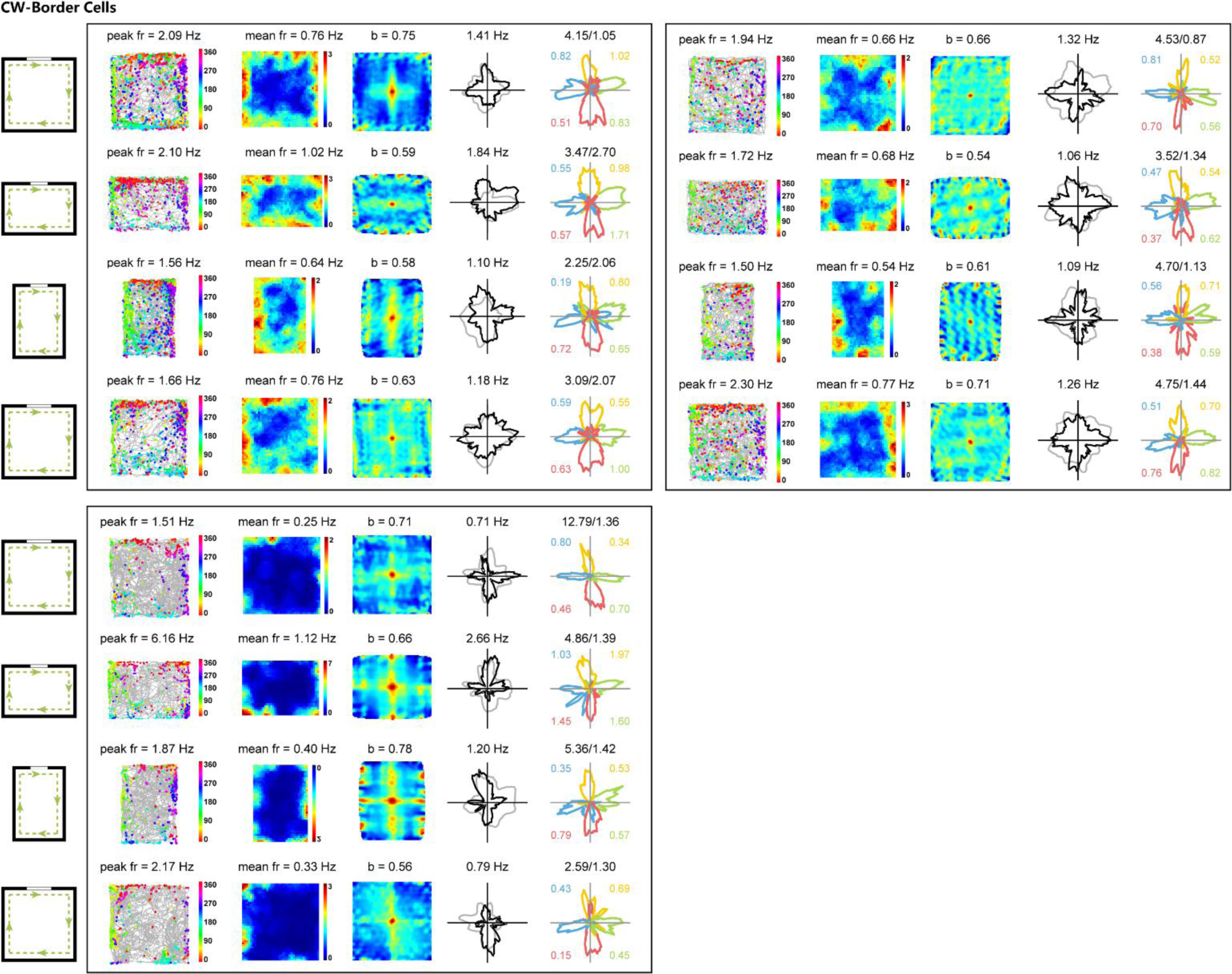
Spatial Firing Patterns for the Entire Sample of CW-Border Cells in Rectangular Boxes. Spatial responses for the entire sample of CW-border cells in the square (S), R_h_ rectangle, R_v_ rectangle and square (S’). Notations and symbols are similar to **Figure 1**.

**Figure S17.**
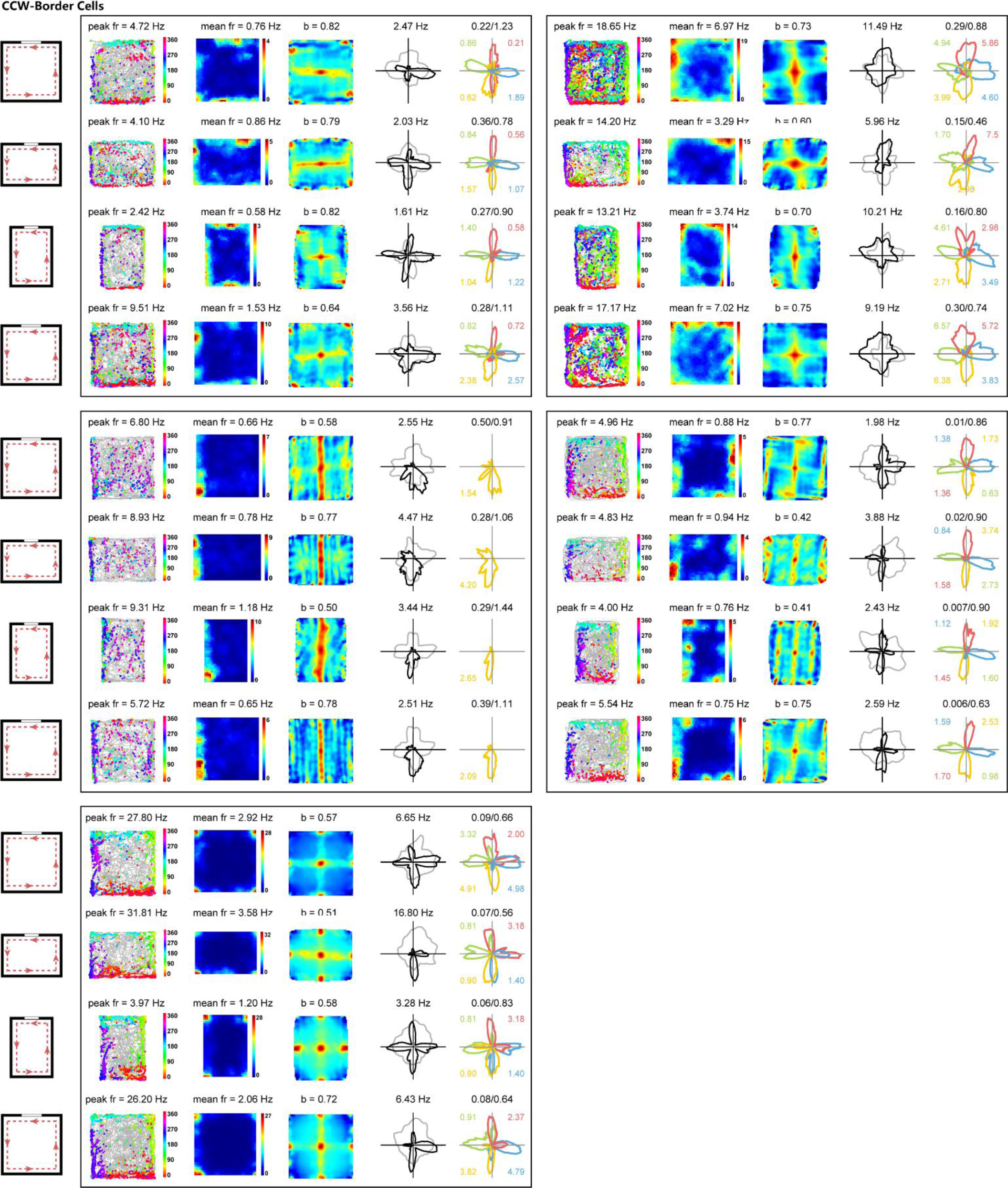
Spatial Firing Patterns for the Entire Sample of CCW-Border Cells in Rectangular Boxes. Spatial responses for the entire sample of CCW-border cells in the square (S), R_h_ rectangle, R_v_ rectangle and square (S’). Notations and symbols are similar to Figure 1. Notations and symbols are similar to **Figure 1**.

**Figure S18.**
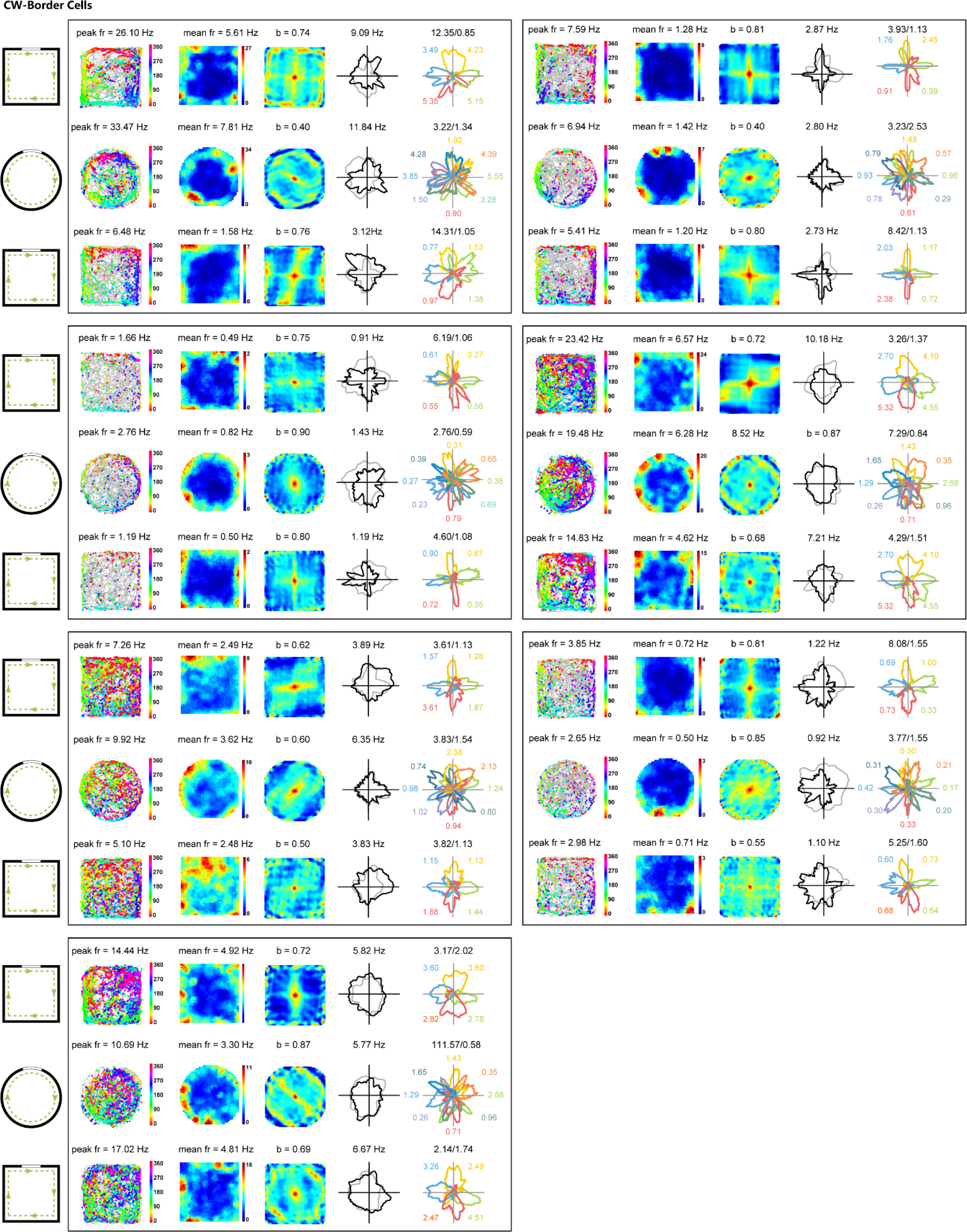
Spatial Firing Patterns for the Entire Sample of CW-Border Cells in the Circular Arena. Spatial responses for the entire sample of CW-border cells in the square (S), Circle (C) and square (S’). Notations and symbols are similar to **Figure 1**.

**Figure S19.**
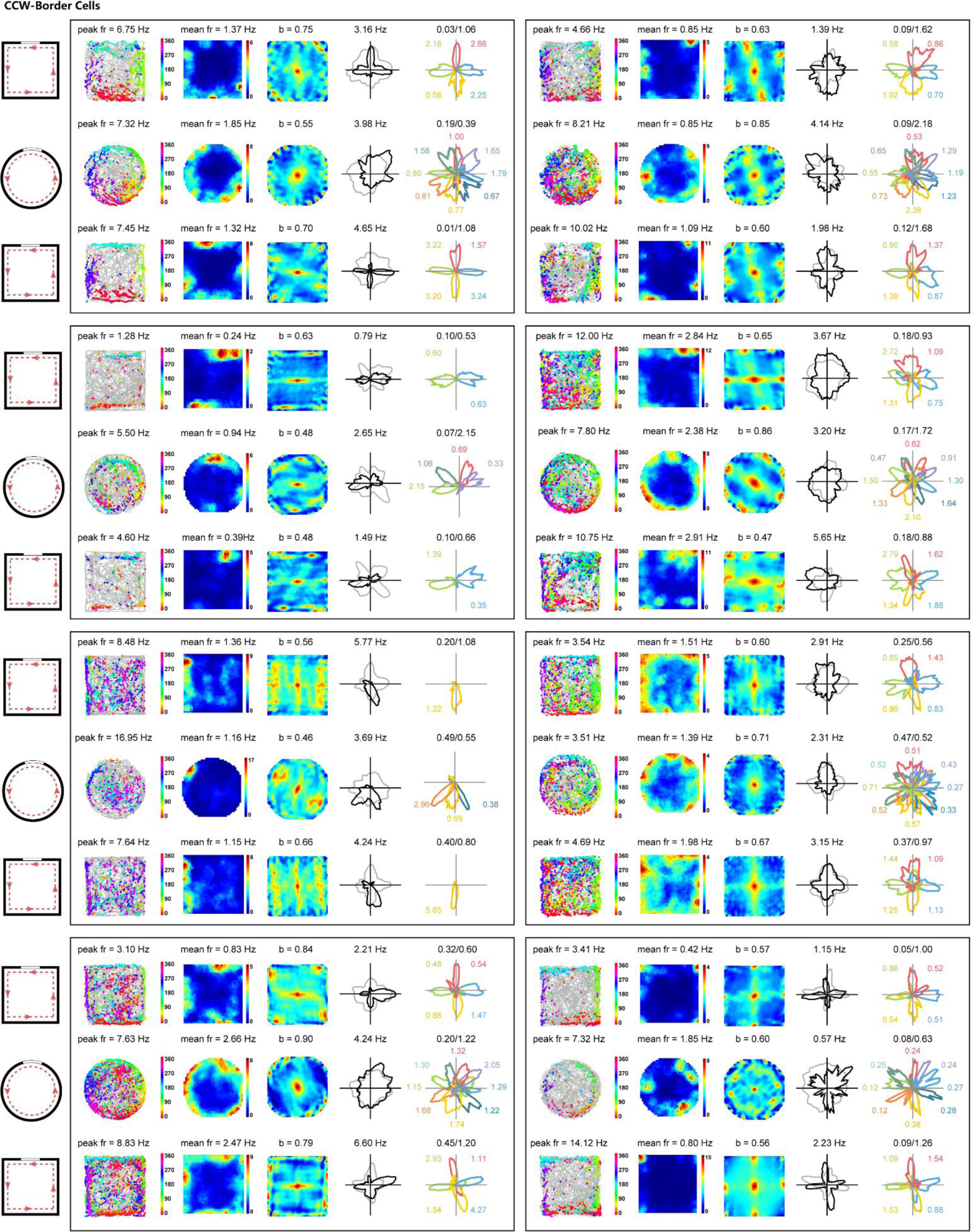
Spatial Firing Patterns for the Entire Sample of CCW-Border Cells in the Circular Arena. Spatial responses for the entire sample of CCW-border cells in the square (S), Circle (C) and square (S’). Notations and symbols are similar to **Figure 1**.

**Figure S20.**
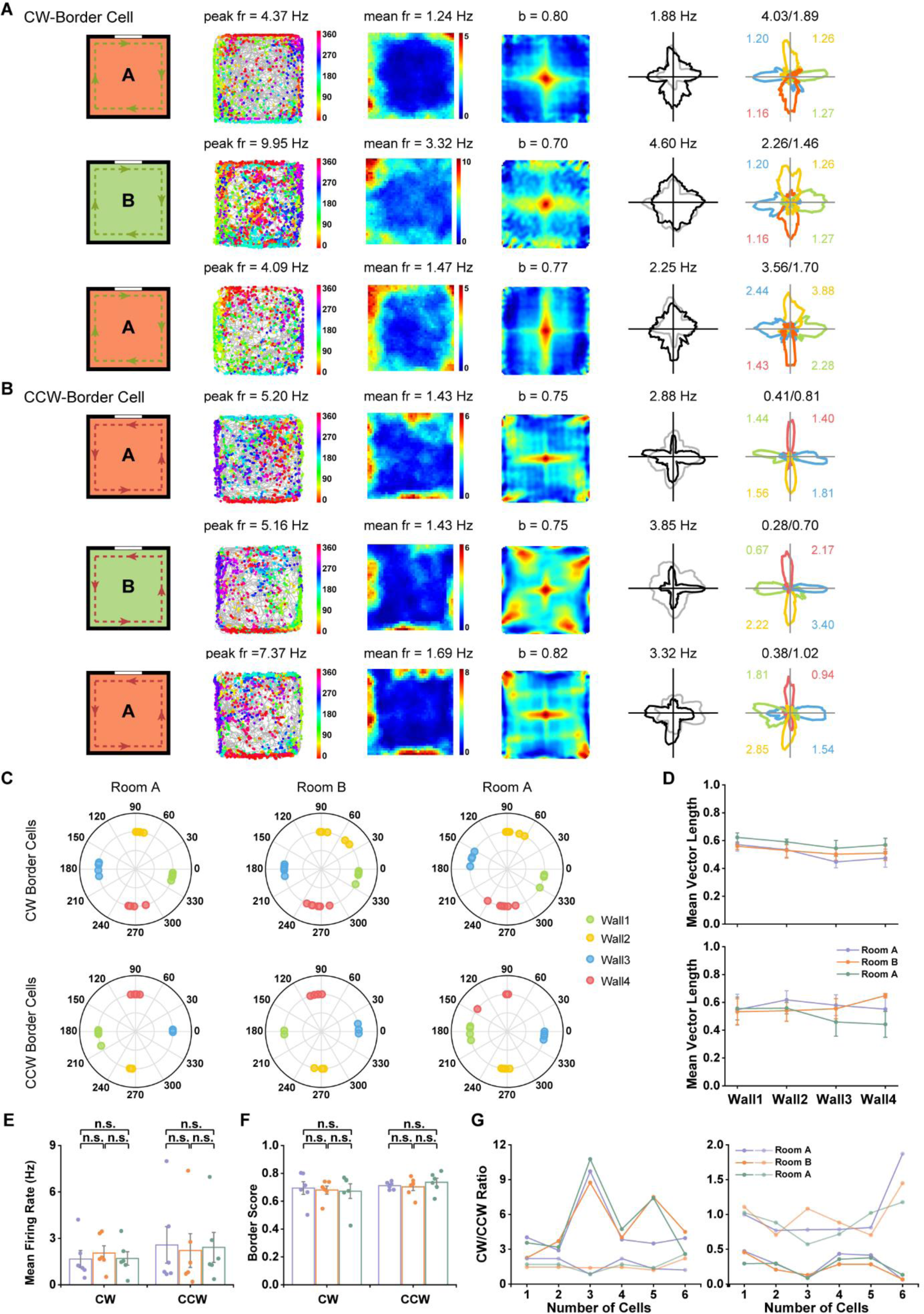
CCW- and CW-Border Cells Were Stable in Novel Environment. (**A**, **B**) Spatial responses of representative cortical CW- and CCW-border cells, respectively, in the familiar (A), novel (B) and back in the familiar environment (A’). Notations and symbols are similar to **Figure 1**. **(C)** Distribution of peak directions for four color-coded sides in the baseline, elevated wall-less platform and baseline sessions. **(D)** Mean vector length showed that the head direction maintained the tuning properties along four sides for both CW- (upper panel) and CCW- (lower panel) border cells in the novel environment. **(E)** Mean firing rate showed no significant change in the novel environment. **(F)** Border score was not statically different between distinct rooms. **(G)** CW- and CCW-border cells, respectively, maintained CW/CCW ratio in firing rate in the novel environments.

**Figure S21.**
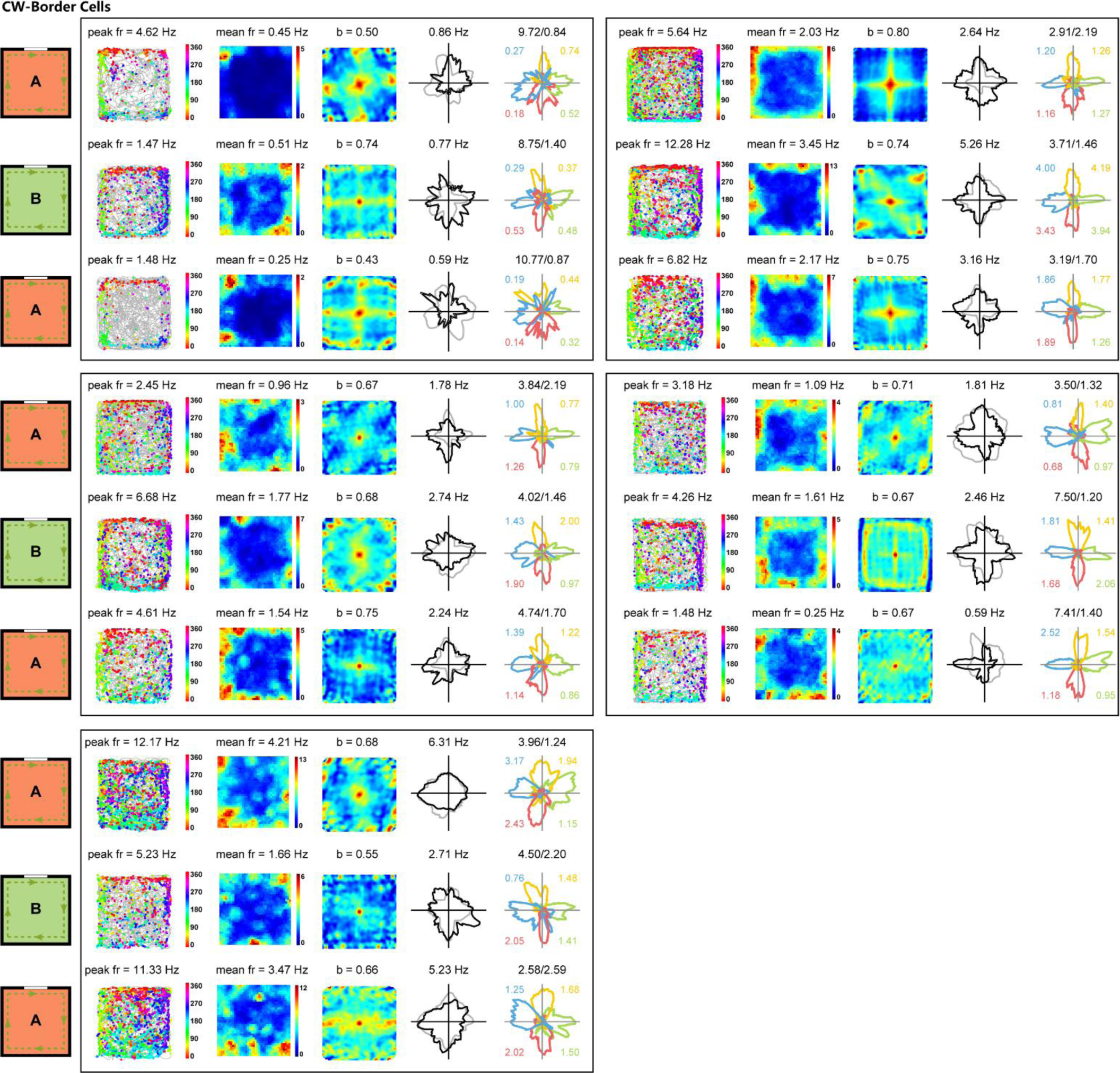
Spatial Firing Patterns for the Entire Sample of CW-Border Cells in Different Environments. Spatial responses for the entire sample of CW-border cells in the A-B-A’ environments. Notations and symbols are similar to **Figure 1**.

**Figure S22.**
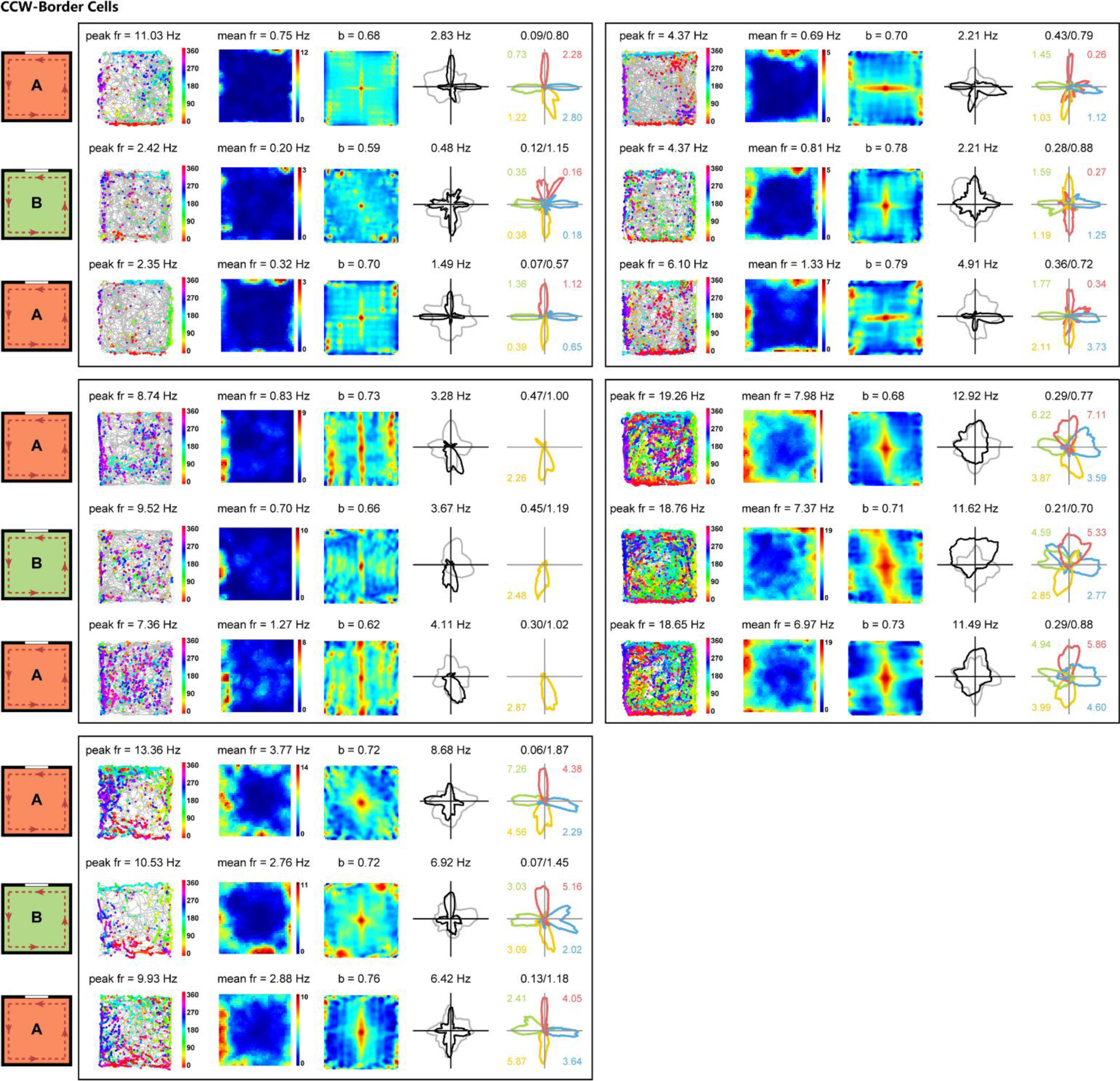
Spatial Firing Patterns for the Entire Sample of CCW-Border Cells in Different Environments. Spatial responses for the entire sample of CCW-border cells in the A-B-A’ environments. Notations and symbols are similar to **Figure 1**.

**Figure S23.**
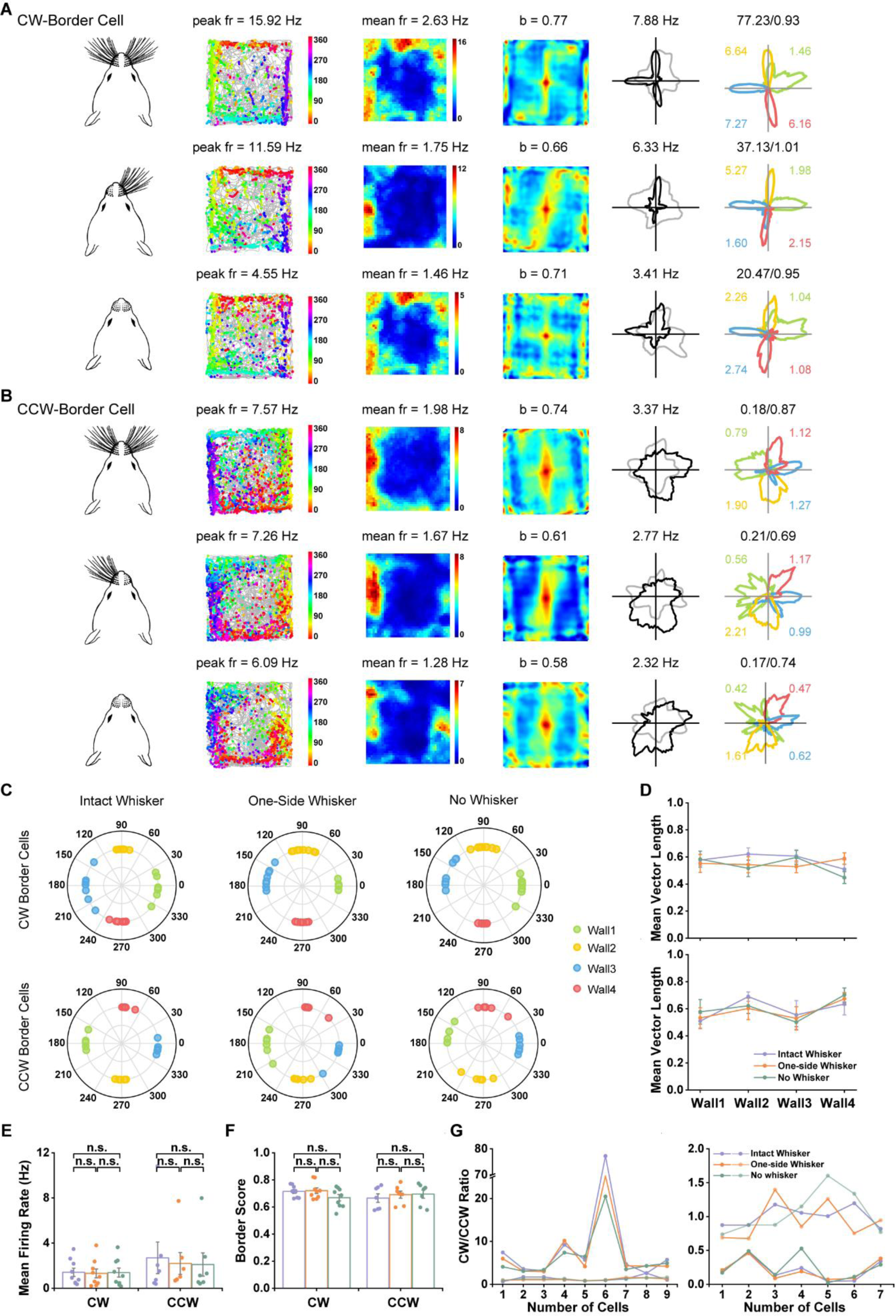
Spatial Firing Patterns of CCW- and CW-Border Cells Were Independent of Whisker Stimulation. (**A**, **B**) Spatial responses of representative cortical CW- and CCW-border cells, respectively, in the presence of whisker (W), one-side whisker (OW) and in the absence of whisker (NW). Notations and symbols are similar to **Figure 1**. **(C)** Distribution of peak directions for four color-coded sides in the presence of whisker (W), one-side whisker (OW) and in the absence of whisker (NW). **(D)** Mean vector length showed that the head direction maintained the tuning properties along four sides for both CW- (left) and CCW- (right) border cells after whisker trimming. **(E)** Mean firing rate showed no significant change after one- or two-side of whisker trimming. **(F)** Border score was not statically different by whisker trimming. **(G)** CW- and CCW-border cells, respectively, maintained CW/CCW ratio in firing rate after one-side whisker trimming and in the absence of whisker.

**Figure S24.**
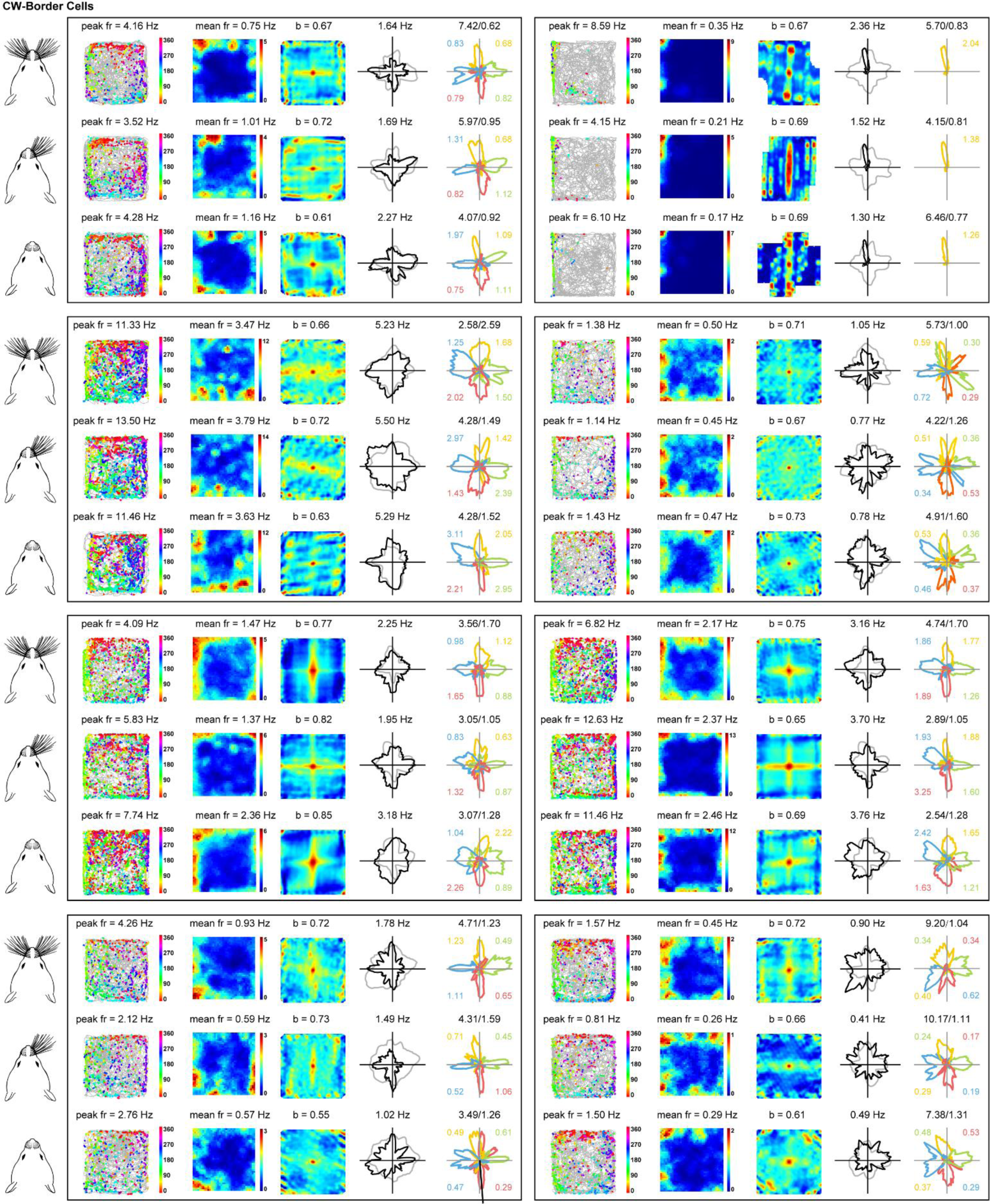
Spatial Firing Patterns for the Entire Sample of CW-Border Cells by Whisker Deprivation. Spatial responses for the entire sample of CW-border cells with intact whisker, one-side whisker trimming and in the absence of whisker. Notations and symbols are similar to **Figure 1**.

**Figure S25.**
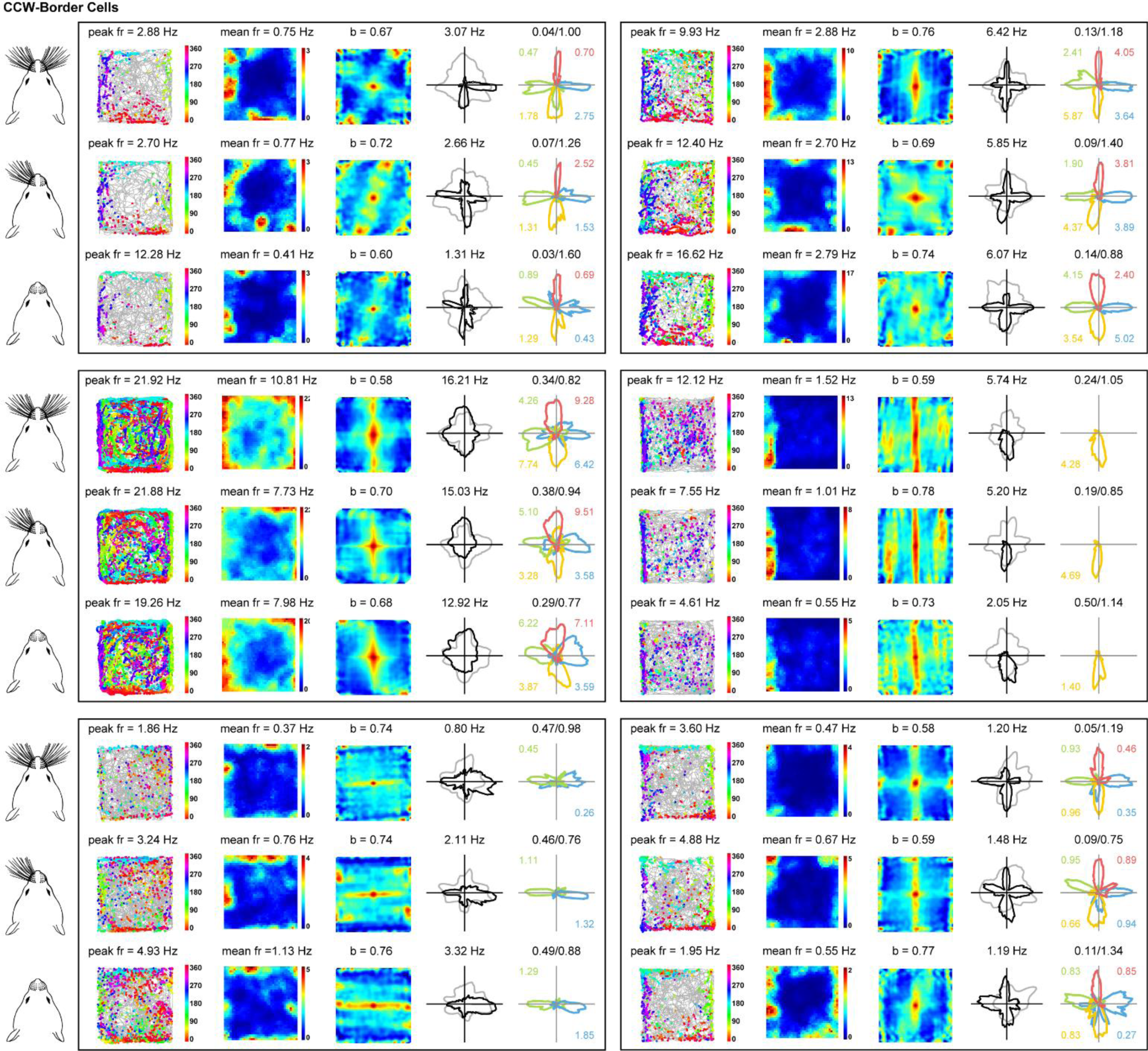
Spatial Firing Patterns for the Entire Sample of CCW-Border Cells by Whisker Deprivation. Spatial responses for the entire sample of CCW-border cells with intact whisker, one-side whisker trimming and in the absence of whisker. Notations and symbols are similar to **Figure 1**.

## REFERENCES

1. Alexander, A.S., Carstensen, L.C., Hinman, J.R., Raudies, F., Chapman, G.W., and Hasselmo, M.E. (2020). Egocentric boundary vector tuning of the retrosplenial cortex. Science advances 6, eaaz2322.

2. Andersen, R.A., Essick, G.K., and Siegel, R.M. (1985). Encoding of spatial location by posterior parietal neurons. Science 230, 456–458.

3. Andersen, R.A., and Mountcastle, V.B. (1983). The influence of the angle of gaze upon the excitability of the light- sensitive neurons of the posterior parietal cortex. The Journal of neuroscience : the official journal of the Society for Neuroscience 3, 532–548.

4. Barry, C., Lever, C., Hayman, R., Hartley, T., Burton, S., O’Keefe, J., Jeffery, K., and Burgess, N. (2006). The boundary vector cell model of place cell firing and spatial memory. Reviews in the neurosciences 17, 71–97.

5. Bartho, P., Hirase, H., Monconduit, L., Zugaro, M., Harris, K.D., and Buzsaki, G. (2004). Characterization of neocortical principal cells and interneurons by network interactions and extracellular features. Journal of neurophysiology 92, 600–608.

6. Bicanski, A., and Burgess, N. (2020). Neuronal vector coding in spatial cognition. Nature reviews Neuroscience 21, 453–470.

7. Byrne, P., and Becker, S. (2008). A principle for learning egocentric-allocentric transformation. Neural computation 20, 709–737.

8. Byrne, P., Becker, S., and Burgess, N. (2007). Remembering the past and imagining the future: a neural model of spatial memory and imagery. Psychological review 114, 340–375.

9. Csicsvari, J., Hirase, H., Czurko, A., and Buzsáki, G. (1998). Reliability and state dependence of pyramidal cell- interneuron synapses in the hippocampus: an ensemble approach in the behaving rat. Neuron 21, 179–189.

10. Deshmukh, S.S., and Knierim, J.J. (2013). Influence of local objects on hippocampal representations: Landmark vectors and memory. Hippocampus 23, 253–267.

11. Duhamel, J.R., Bremmer, F., Ben Hamed, S., and Graf, W. (1997). Spatial invariance of visual receptive fields in parietal cortex neurons. Nature 389, 845–848.

12. Gofman, X., Tocker, G., Weiss, S., Boccara, C.N., Lu, L., Moser, M.B., Moser, E.I., Morris, G., and Derdikman, D. (2019). Dissociation between Postrhinal Cortex and Downstream Parahippocampal Regions in the Representation of Egocentric Boundaries. Current biology : CB 29, 2751–2757 e2754.

13. Hafting, T., Fyhn, M., Molden, S., Moser, M.B., and Moser, E.I. (2005). Microstructure of a spatial map in the entorhinal cortex. Nature 436, 801–806.

14. Hartley, T., Burgess, N., Lever, C., Cacucci, F., and O’Keefe, J. (2000). Modeling place fields in terms of the cortical inputs to the hippocampus. Hippocampus 10, 369–379.

15. Hartley, T., and Lever, C. (2014). Know your limits: the role of boundaries in the development of spatial representation. Neuron 82, 1–3.

16. Hawkins, J., Lewis, M., Klukas, M., Purdy, S., and Ahmad, S. (2018). A Framework for Intelligence and Cortical Function Based on Grid Cells in the Neocortex. Frontiers in neural circuits 12, 121.

17. Hinman, J.R., Chapman, G.W., and Hasselmo, M.E. (2019). Neuronal representation of environmental boundaries in egocentric coordinates. Nature communications 10, 2772.

18. Hoydal, O.A., Skytoen, E.R., Andersson, S.O., Moser, M.B., and Moser, E.I. (2019). Object-vector coding in the medial entorhinal cortex. Nature 568, 400–404.

19. Jercog, P.E., Ahmadian, Y., Woodruff, C., Deb-Sen, R., Abbott, L.F., and Kandel, E.R. (2019). Heading direction with respect to a reference point modulates place-cell activity. Nature communications 10, 2333.

20. Killian, N.J., Jutras, M.J., and Buffalo, E.A. (2012). A map of visual space in the primate entorhinal cortex. Nature 491, 761–764.

21. Kinkhabwala, A.A., Gu, Y., Aronov, D., and Tank, D.W. (2020). Visual cue-related activity of cells in the medial entorhinal cortex during navigation in virtual reality. eLife 9.

22. Kunz, L., Brandt, A., Reinacher, P.C., Staresina, B.P., Reifenstein, E.T., Weidemann, C.T., Herweg, N.A., Tsitsiklis, M., Kempter, R., Kahana, M.J., et al. (2021). A neural code of egocentric spatial information in human medial temporal lobe. bioRxiv, 10.1101/2020.1103.1103.973131.

23. LaChance, P.A., Todd, T.P., and Taube, J.S. (2019). A sense of space in postrhinal cortex. Science 365, eaax4192.

24. Latuske, P., Toader, O., and Allen, K. (2015). Interspike Intervals Reveal Functionally Distinct Cell Populations in the Medial Entorhinal Cortex. The Journal of neuroscience : the official journal of the Society for Neuroscience 35, 10963–10976.

25. Lever, C., Burton, S., Jeewajee, A., O’Keefe, J., and Burgess, N. (2009). Boundary vector cells in the subiculum of the hippocampal formation. The Journal of neuroscience : the official journal of the Society for Neuroscience 29, 9771–9777.

26. Long, X., Deng, B., Cai, J., Chen, Z.S., and Zhang, S.-J. (2021). A compact spatial map in V2 visual cortex. bioRxiv, 10.1101/2021.1102.1111.430687.

27. Long, X., Young, C.K., and Zhang, S.-J. (2020). Sharp tuning of head direction by somatosensory fast-spiking interneurons. bioRxiv, 10.1101/2020.1102.1103.933143.

28. Long, X., and Zhang, S.J. (2021). A novel somatosensory spatial navigation system outside the hippocampal formation. Cell research, 10.1038/s41422-41020-00448-41428.

29. Madl, T., Chen, K., Montaldi, D., and Trappl, R. (2015). Computational cognitive models of spatial memory in navigation space: a review. Neural networks : the official journal of the International Neural Network Society 65, 18–43.

30. McNaughton, B.L., Knierim, J.J., and Wilson, M.A. (1995). Vector encoding and the vestibular foundations of spatial cognition: Neurophysiological and computational mechanisms. In MS Gazzaniga (Ed), The Cognitive Neurosciences (The MIT Press, Cambridge), pp. 585–595.

31. O’Keefe, J., and Dostrovsky, J. (1971). The hippocampus as a spatial map. Preliminary evidence from unit activity in the freely-moving rat. Brain research 34, 171–175.

32. Olson, J.M., Li, J.K., Montgomery, S.E., and Nitz, D.A. (2020). Secondary Motor Cortex Transforms Spatial Information into Planned Action during Navigation. Current biology : CB 30, 1845–1854 e1844.

33. Peyrache, A., Dehghani, N., Eskandar, E.N., Madsen, J.R., Anderson, W.S., Donoghue, J.A., Hochberg, L.R., Halgren, E., Cash, S.S., and Destexhe, A. (2012). Spatiotemporal dynamics of neocortical excitation and inhibition during human sleep. Proceedings of the National Academy of Sciences of the United States of America 109, 1731–1736.

34. Peyrache, A., and Duszkiewicz, A.J. (2021). A spatial map out of place. Cell research.

35. Peyrache, A., Schieferstein, N., and Buzsáki, G. (2017). Transformation of the head-direction signal into a spatial code. Nature communications 8, 1752.

36. Poulter, S., Lee, S.A., Dachtler, J., Wills, T.J., and Lever, C. (2021). Vector trace cells in the subiculum of the hippocampal formation. Nature neuroscience 24, 266–275.

37. Rolls, E.T., and Mills, P. (2019). The Generation of Time in the Hippocampal Memory System. Cell reports 28, 1649–1658 e1646.

38. Sarel, A., Finkelstein, A., Las, L., and Ulanovsky, N. (2017). Vectorial representation of spatial goals in the hippocampus of bats. Science 355, 176–180.

39. Sargolini, F., Fyhn, M., Hafting, T., McNaughton, B.L., Witter, M.P., Moser, M.B., and Moser, E.I. (2006). Conjunctive representation of position, direction, and velocity in entorhinal cortex. Science 312, 758–762.

40. Savelli, F., Yoganarasimha, D., and Knierim, J.J. (2008). Influence of boundary removal on the spatial representations of the medial entorhinal cortex. Hippocampus 18, 1270–1282.

41. Solstad, T., Boccara, C.N., Kropff, E., Moser, M.B., and Moser, E.I. (2008). Representation of geometric borders in the entorhinal cortex. Science 322, 1865–1868.

42. Stangl, M., Topalovic, U., Inman, C.S., Hiller, S., Villaroman, D., Aghajan, Z.M., Christov-Moore, L., Hasulak, N.R., Rao, V.R., Halpern, C.H., et al. (2021). Boundary-anchored neural mechanisms of location-encoding for self and others. Nature 589, 420–425.

43. Stewart, S., Jeewajee, A., Wills, T.J., Burgess, N., and Lever, C. (2014). Boundary coding in the rat subiculum. Philosophical transactions of the Royal Society of London Series B, Biological sciences 369, 20120514.

44. Taube, J.S., Muller, R.U., and Ranck, J.B., Jr. (1990). Head-direction cells recorded from the postsubiculum in freely moving rats. I. Description and quantitative analysis. The Journal of neuroscience : the official journal of the Society for Neuroscience 10, 420–435.

45. van Wijngaarden, J.B., Babl, S.S., and Ito, H.T. (2020). Entorhinal-retrosplenial circuits for allocentric-egocentric transformation of boundary coding. eLife 9.

46. Wang, C., Chen, X., and Knierim, J.J. (2020). Egocentric and allocentric representations of space in the rodent brain. Current opinion in neurobiology 60, 12–20.

47. Wang, C., Chen, X., Lee, H., Deshmukh, S.S., Yoganarasimha, D., Savelli, F., and Knierim, J.J. (2018). Egocentric coding of external items in the lateral entorhinal cortex. Science 362, 945–949.

48. Whitlock, J.R., Pfuhl, G., Dagslott, N., Moser, M.B., and Moser, E.I. (2012). Functional split between parietal and entorhinal cortices in the rat. Neuron 73, 789–802.

49. Wilber, A.A., Clark, B.J., Forster, T.C., Tatsuno, M., and McNaughton, B.L. (2014). Interaction of egocentric and world-centered reference frames in the rat posterior parietal cortex. The Journal of neuroscience : the official journal of the Society for Neuroscience 34, 5431–5446.

## Supplementary references

50. Boccara, C.N., Sargolini, F., Thoresen, V.H., Solstad, T., Witter, M.P., Moser, E.I., and Moser, M.B. (2010). Grid cells in pre- and parasubiculum. Nature neuroscience 13, 987–994.

51. Long, X., and Zhang, S.J. (2021). A novel somatosensory spatial navigation system outside the hippocampal formation. Cell research, 10.1038/s41422-41020-00448-41428.

52. Paxinos, G., and Watson, C. (2007). The rat brain in stereotaxic coordinates, Vol 552 (Amsterdam: Elsevier).

53. Sargolini, F., Fyhn, M., Hafting, T., McNaughton, B.L., Witter, M.P., Moser, M.B., and Moser, E.I. (2006). Conjunctive representation of position, direction, and velocity in entorhinal cortex. Science 312, 758–762.

54. Schmitzer-Torbert, N., Jackson, J., Henze, D., Harris, K., and Redish, A.D. (2005). Quantitative measures of cluster quality for use in extracellular recordings. Neuroscience 131, 1–11.

55. Solstad, T., Boccara, C.N., Kropff, E., Moser, M.B., and Moser, E.I. (2008). Representation of geometric borders in the entorhinal cortex. Science 322, 1865–1868.

56. Zhang, S.J., Ye, J., Miao, C., Tsao, A., Cerniauskas, I., Ledergerber, D., Moser, M.B., and Moser, E.I. (2013). Optogenetic dissection of entorhinal-hippocampal functional connectivity. Science 340, 1232627.

